# Genome resequencing reveals rapid, repeated evolution in the Colorado potato beetle, *Leptinotarsa decemlineata*

**DOI:** 10.1101/2021.02.09.430453

**Authors:** Benjamin Pélissié, Yolanda H. Chen, Zachary P. Cohen, Michael S. Crossley, David J. Hawthorne, Victor Izzo, Sean D. Schoville

## Abstract

**Background:** Insecticide resistance and rapid pest evolution threatens food security and the development of sustainable agricultural practices. An improved understanding of the evolutionary mechanisms that allow pests to rapidly adapt to novel control tactics will help prevent economically damaging outbreaks. The Colorado potato beetle (CPB), *Leptinotarsa decemlineata,* is a global super-pest that rapidly evolves resistance to insecticides. Using whole genome resequencing and transcriptomic data focused on its ancestral and pest range in North America, we assess evidence for three, non-mutually exclusive models of rapid evolution: pervasive selection on novel mutations, rapid regulatory evolution, and repeated selection on standing genetic variation.

**Results:** Population genomic analysis demonstrates that CPB is geographically structured, even among recently established pest populations. Pest populations exhibit only modest reductions in nucleotide diversity, relative to non-pest ancestral populations, and show evidence of recent demographic expansion. Genome scans of selection provide clear signatures of repeated adaptation across different CPB populations, with especially strong evidence that insecticide resistance involves selection of different genes in different populations. Similarly, analyses of gene expression show that constitutive upregulation of candidate insecticide resistance genes drives distinctive population patterns.

**Conclusion:** CPB evolves insecticide resistance repeatedly across agricultural regions, and oftentimes at the same loci, supporting a prominent role of polygenic evolution from standing genetic variation. Despite expectations, we do not find support for strong selection on novel mutations, or rapid evolution from selection on regulatory genes. An important future goal will be to understand how polygenic resistance phenotypes spread among local pest populations, in order to refine integrated pest management practices to maintain the efficacy and sustainability of novel control techniques.

## Background

Herbivorous pests cause an estimated 18-20% damage to crops and cost nearly $470 billion annually on a global scale [1]. The ability of insect pests to evolve resistance to insecticides threatens food security and the development of sustainable agricultural practices, especially when their rate of evolution outstrips the development of novel control strategies [2–4]. This is the case with insect ‘super-pests,’ which repeatedly evolve insecticide resistance even as they are faced with completely novel insecticides, thus perpetuating the arms race that defines the pesticide treadmill [5]. Curiously, particular super-pest species or even select populations are more likely to adapt to new compounds, suggesting that there is a genetic basis in the propensity to evolve resistance [6]. Yet, despite more than 60 years of research on the evolution of resistance [7], the relative importance of alternative mechanisms that underlie the evolutionary potential for pesticide resistance evolution are still unclear [8, 9]. Although a considerable effort has been placed on understanding the proximal molecular control of resistance [10], broader questions about the genetic complexity of resistance, mode of selection and geographical extent of adaptation have rarely been studied [11–13]. While population genetic models of resistance management have been highly effective in certain management scenarios [14], observed patterns of insecticide resistance evolution defy many of the assumptions of our evolutionary models [15]. Recent genomic resequencing datasets suggest that resistance evolution is sometimes geographically and genetically complex [16–19].

A key goal should be to understand the evolutionary processes that allow species to become pests, particularly the mechanisms underlying phenotypic shifts that result in economically damaging pest outbreaks [8, 20]. A prevalent view is that pests, including invasive species, often retain substantial genetic diversity that facilitates evolution in agroecosystems [21–23]. However, there is increasing recognition that evolution can be rapid irrespective of levels of standing genetic diversity [24]. While not all pests exhibit rapid rates of adaptation to insecticides [6], insect super-pests often demonstrate repeated rapid evolution [25, 26]. Rapid evolution is defined as a shift in phenotype from underlying variation in exceptionally few generations [27], and can occur as a result of several mechanisms [28]. First, selection can act on novel mutations, which may arise frequently if pests have intrinsically large population sizes (much greater than >>10^6^) and are not mutation-limited [29, 30]. This could lead to repeated evolution of resistance among different populations, most likely with independent mutations at different loci underlying resistance phenotypes. Second, as a special case of the first mechanism, key mutational changes could affect a master regulatory gene [31–33], where mutations drive expression of the same downstream molecular pathways in different populations. Rapid gene regulatory evolution has been raised as a possible mechanism underlying repeated evolution of pesticide resistance in the spider mite *Tetranychus urticae*, where it has been linked to a transcriptional cascade in xenobiotic detoxification [34]. Third, an alternative pathway of rapid evolution would draw on standing genetic variation [7]. Standing genetic variation is increasingly viewed as a common source of rapid adaptive variation [35], because the initial frequency of mutations in a population determine the rate at which populations respond to selection pressures [36, 37]. While population size must typically be large to retain large reservoirs of standing variation, admixture among divergent populations can increase standing variation [38, 39]. Furthermore, standing genetic variation can also be present in the form of redundancy in molecular pathways that are critical to pesticide resistance phenotypes [e.g. in generalist herbivorous insects that specialize on toxic plants: 40, 41, 42], rather than allelic diversity *per se*. It should be emphasized that these mechanisms of adaptation need not be exclusive, yet it remains unclear how each contributes to the evolutionary success of the top arthropod super-pests. Emerging genomic datasets provide the opportunity to detect and quantify the importance of different mechanisms underlying rapid evolutionary change by screening for genomic signatures of selection [8].

The Colorado potato beetle (CPB), *Leptinotarsa decemlineata,* is a global super-pest and an especially tractable exemplar of rapid evolution to insecticides. CPB has evolved resistance to over 50 different insecticides in all the major classes, in some cases within the first year of use [43]. CPB has demonstrated an ability to rapidly evolve in response to a wide range of environmental pressures, including host-plant defenses and climatic variability [44, 45]. This super-pest originated in the Great Plains region of the U.S. [46], following a host shift to potato (an introduced crop) in the mid-19th century (around 1866) that allowed for rapid spatial expansion from Nebraska to the Eastern U.S. in a 20 year period and colonization of Eurasia by the early 1900s [47–49]. Despite rapid spatial expansion, populations are genetically differentiated [49, 50] and insecticide resistance is geographically heterogeneous [51], even over local landscape scales [52]. In particular, beetles from Long Island, New York are known to have the highest levels of baseline resistance and are typically the first populations to develop resistance to all compounds [53], while populations in the Pacific Northwest remain susceptible to insecticides despite an equivalent duration of usage and comparable treatment practices [54, 55]. Non-pest populations are found in the Great Plains and Mexico, where they use ancestral host plants (primarily *Solanum rostratum*) [56]. Closely related congeners in the genus *Leptinotarsa* are sympatric in the southern part of CPB’s geographical range [57]. By integrating across this diversity, CPB can serve as a model for understanding evolutionary mechanisms that facilitate and constrain rapid evolution.

Here we leverage the recent publication of the CPB genome [58] to investigate whether repeatable patterns of evolution occur in highly resistant pest populations. We compare CPB genomic and transcriptomic variation across populations in the U.S., Mexico, and Europe, as well as closely related *Leptinotarsa* species, to assess three competing models of rapid evolution: pervasive selection on *de novo* mutation (independent hard selective sweeps in geographically separate populations), rapid regulatory evolution, or repeated selection on standing genetic variation (see **Table 1** for predictions). We also provide detailed description of genomic diversity patterns, evolutionary relationships, and the population history of CPB pest lineages, in order to understand how expansion history has influenced geographical variation in insecticide resistance. Over the long-term, by improving our understanding of the evolutionary processes and genomic mechanisms underlying the ability to repeatedly evolve insecticide resistance in super-pests, integrated pest management strategies can be developed to provide more sustainable agricultural practices [2, 22].

**Table 1.** Predictions from alternative mechanisms of rapid evolution to insecticide resistance in Colorado potato beetle.

| Evolutionary mechanism | Geographical pattern in resistant populations | Genome scans of selection | Haplotype-based selection scan | Differential gene expression |
| --- | --- | --- | --- | --- |
| *De novo* mutation | Independent hard selective sweeps | A few statistically significant candidate genes | Long haplotype blocks around selected loci | Differential gene expression limited to key pathways |
| Rapid regulatory evolution | Selection in key regulatory genes leading to repeated upregulation of resistance pathways | A few statistically significant regulatory genes | Long haplotype blocks around regulatory genes encoding transcription factors | Differential expression of key transcription factors and constitutive expression differences in insecticide resistance pathways |
| Standing genetic diversity | Repeated selection on candidate insecticide resistance genes | Numerous statistically significant candidate genes | Short haplotype blocks around selected loci | Multiple differentially expressed genes in the same molecular pathways and constitutive expression differences in insecticide resistance pathways |

## Results

### Extensive Genomic Diversity within CPB

We examined short-read whole genome sequences for a geographically dispersed set of 85 samples, including six geographically proximate pairs of susceptible and resistant samples, as well as nine additional *Leptinotarsa* species (**Fig. 1** and **Additional File 1: Table S1** and **Fig. S2**). Employing best practices in genotype ascertainment, we sequenced each sample (most resulted in coverage >4x, **Fig. S3**), and found that CPB shows considerable genomic diversity, with 76,647,868 single nucleotide polymorphisms (SNPs) recovered from the nuclear genome (**Additional File 1: Table S2**; the CPB genome is estimated to be ∼670 Mb in size). Following variant recalibration, there were a total of 47,969,460 SNPs, of which 30,973,249 were in intergenic regions.

**Figure 1.**
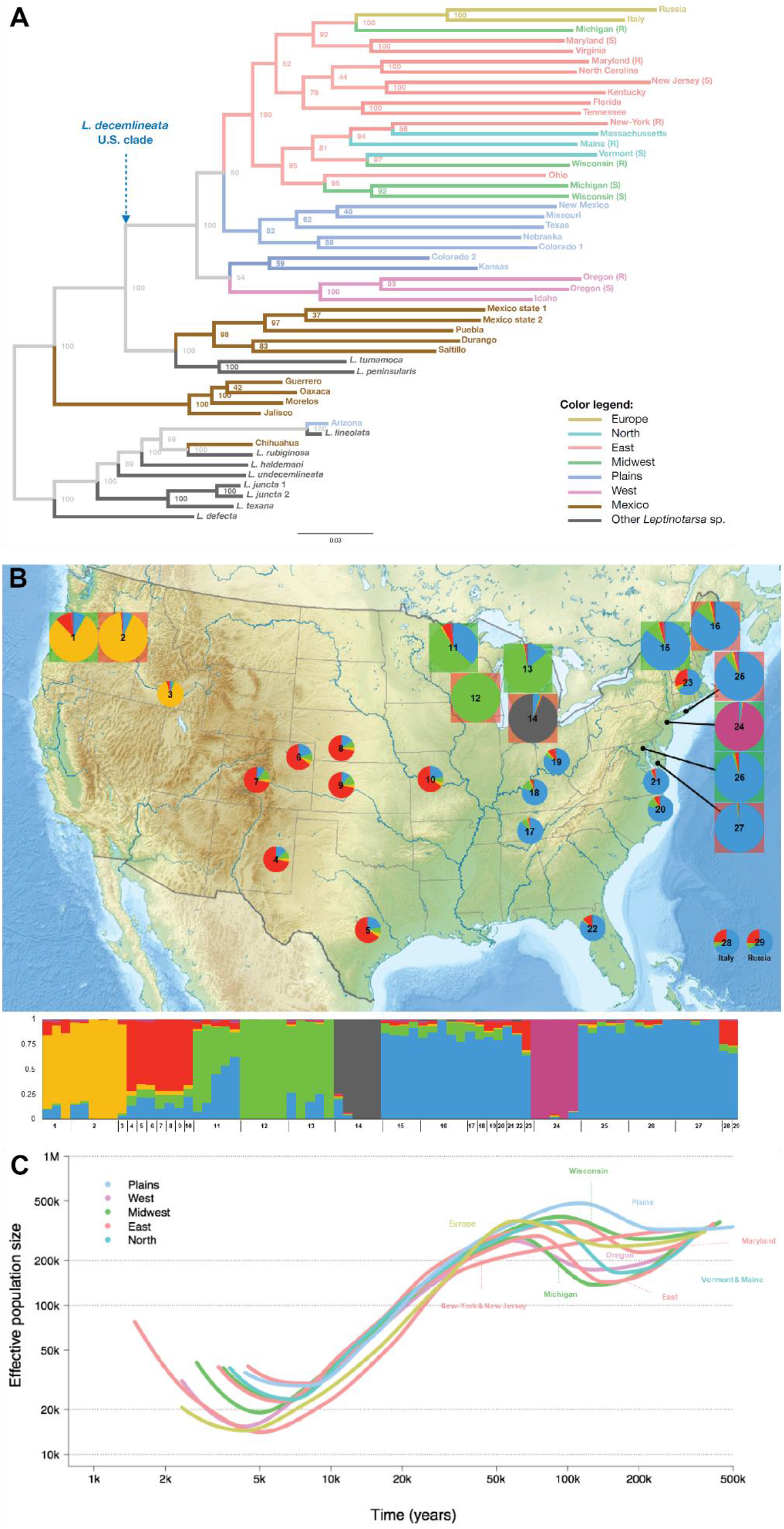
**A)** Unrooted phylogenetic tree of *Leptinotarsa* species obtained with SNPhylo, based on 35,838 SNPs (after LD-based reduction). Node labels represent bootstrap values. The blue arrow highlights a monophyletic clade comprising CPB samples collected in the U.S. and Europe. (S): imidacloprid susceptible population; (R): imidacloprid resistant population. **B)** Geographical sampling of *Leptinotarsa decemlineata* and estimated admixture coefficients. Admixture proportions were estimated with *SNMF* on the intergenic “CPB” dataset for k=6 clusters, and are shown as both pie charts and an individual bar-plot. Each pie chart represents a sampled location (small charts for single samples; large ones for populations of five individuals), referenced as a number. Colored boxes around large pie charts differentiate susceptible (green) vs. resistant samples (red). **C)** Population demographic histories (median Ne only) estimated from SMC++. Colors correspond to geographical regions.

We estimated genome-wide nucleotide diversity (π) using a 10 Kb sliding window. Within-population genetic diversity of non-pest CPB samples from the inferred ancestral source population, the U.S. Plains region, was high with an average π = 0.005. Comparison of nucleotide diversity between the Plains and U.S. pest populations shows a clear reduction in nucleotide diversity (average π = 0.0028 and 0.003 for co-located pairs of pesticide resistant and susceptible populations, respectively; **Additional File 1: Fig. S4**). However, pooling co-located pairs of samples increased nucleotide diversity in a given agricultural region by 40% and eroded the difference from the Plains samples, showing that high levels of variation have been retained in agricultural regions. Individual heterozygosity appeared to be reduced relative to expectations under random mating, as measured by the inbreeding coefficient *F_IS_*. Inbreeding was higher in pest populations (on average *F_IS_* = 0.603 and 0.557 for susceptible and resistant populations, respectively; **Fig. S5**), relative to Plains individuals (on average *F_IS_* = 0.531). The New Jersey lab population, which was maintained as a breeding colony for pesticide assays, showed the highest level of inbreeding (*F_IS_* = 0.723). Susceptible and resistant pest samples had a comparable number of private alleles (93,407 vs. 89,599, respectively; **Additional File 1: Fig. S6** and **S7**), but Plains samples had three times as many private alleles (262,140 vs. 91,503, respectively). Combining observations of nucleotide diversity, inbreeding, and private alleles suggests that pest lineages lost genetic variation as they expanded into agricultural habitats, but do not appear to have suffered from a strong genetic bottleneck.

### Evolutionary Diversification and Demographic History

*Leptinotarsa decemlineata* population structure is clearly driven by geographic isolation. Phylogenetic reconstruction (**Fig. 1A**) clearly separated the Mexican samples from all U.S. samples. All CPB populations from potato growing regions in the U.S. and Eurasia formed a well-supported monophyletic group, while separate clades contained samples in the Pacific Northwest, Southwest U.S. and the U.S. Plains region. The Arizona CPB sample was found outside the CPB clade, next to the *L. lineolata* sample, suggesting high divergence and possibly cryptic species status. Interestingly, Mexican CPB and other *Leptinotarsa* species were mixed together, also suggesting that Mexican CPB belong to an unidentified cryptic *Leptinotarsa* species. Due to their significant genetic divergence, Mexican CPB, the Arizona sample, and other *Leptinotarsa* species were removed from downstream analyses of population structure and demographic change, as their inclusion would violate statistical assumptions in those approaches. Examining U.S. CPB samples, genetic divergence among populations was modest (average pairwise *F_ST_* = 0.09; **Additional File 1: Fig. S8** and **Table S3**), but exceeded >0.1 when New Jersey or the Michigan resistant population were compared. Principal component and population ancestry analyses converged in showing clear geographical patterns of population genetic structure (**Fig. 1B**; **Additional File 1: Figs. S9**-**S12**). Admixture-based clustering supported six populations, which represent the New-Jersey population, a western population (Oregon plus Idaho), a Plains population, a Midwestern population, a distinctive Michigan resistant population, and an Eastern U.S. population (which includes the introduced European samples). Admixture tests using the *D* statistic were examined among CPB pest and non-pest populations, as well as with other *Leptinotarsa* species. These tests provided limited evidence of admixture contributing to genetic diversity of pest lineages (**Fig. 2**), ruling out the hypothesis of standing genetic variation increasing in the pest lineage as a result of hybridization. The highest *D_min_* values suggest historical admixture from Mexico to the Plains (0.036) and Mexico to the Western populations (0.044), and limited ongoing gene flow between New York and the Michigan susceptible population (*D* = 0.015). Similarly, an assessment of admixture with other *Leptinotarsa* species (**Additional File 1: Fig. S13**) does not suggest recent gene flow into pest populations.

**Figure 2.**
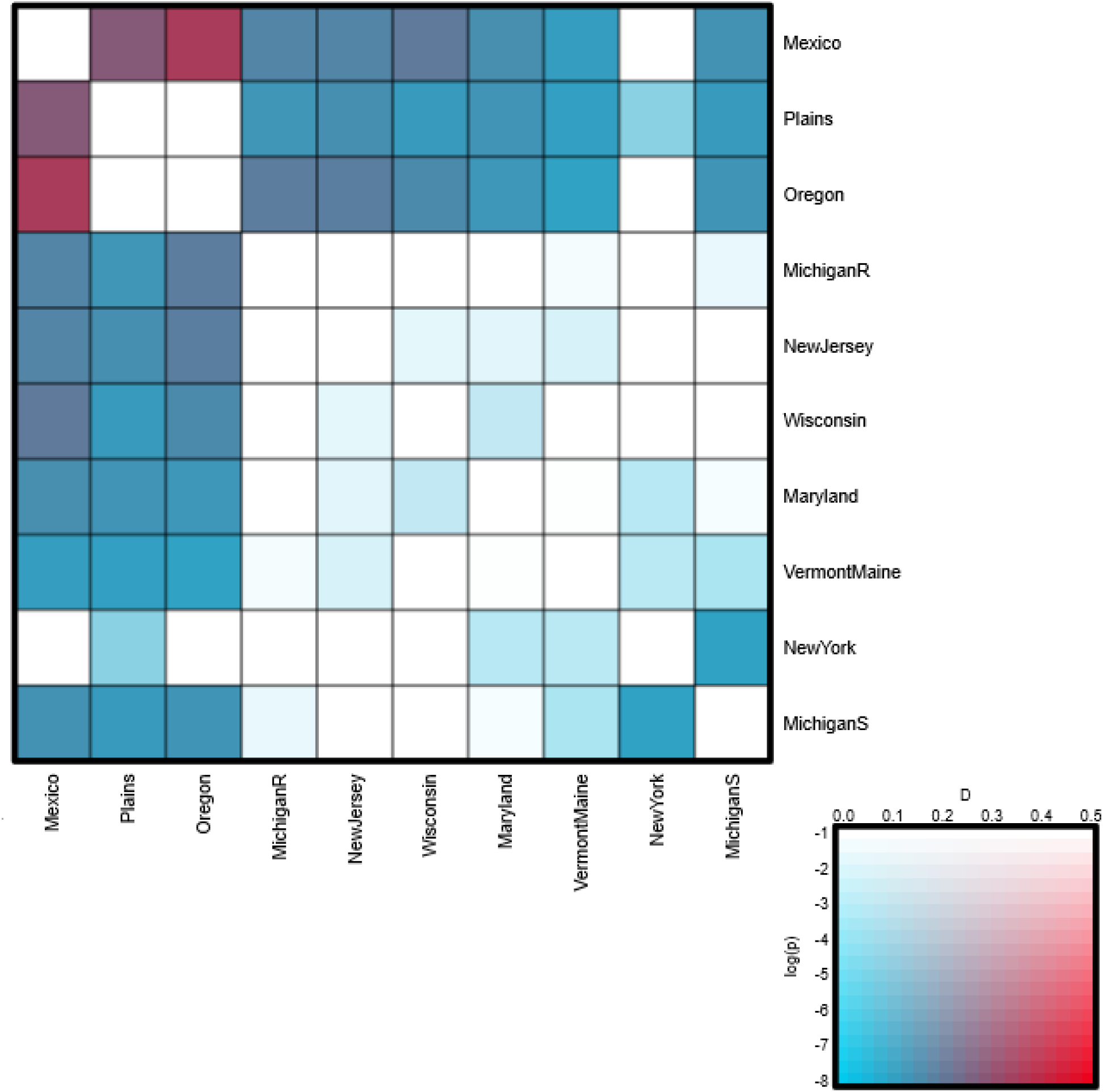
Heatmap of *D*-statistics, showing the introgression patterns among CPB populations. The color of the heatmap cell indicates the most significant *D_min_* found with every population pairs: red colors indicate higher *D*-statistics, and generally more saturated colors indicate higher *P*-values. The complete biallelic dataset was analyzed.

Demographic reconstruction of CPB populations using SMC++ and Stairway plot analysis (**Fig. 1C** and **Additional File 1: Fig. S14-S15**) showed consistent population size fluctuations through time, with similar trajectories for all pest populations. Nearly all agricultural populations exhibited recent population size increases in the SMC++ analysis, most notably in the Eastern U.S. The split time analysis suggested an early split of the western populations from the Plains region, and subsequent near-simultaneous split times of Midwest and Eastern populations (**Additional File 1: Fig. S16)**. Applying a mutation rate from other insect taxa (2.1 × 10^-9^ substitutions per site per generation) suggests that populations contracted between 300k and 100k years ago, expanded between 200k and 70k years, and declined again until 10k to 5k years ago. The splits of most pest populations from the sampled populations in the Plains occurred between 21k and 11k years, during the transition from the late Pleistocene to early Holocene.

### Genome-wide Patterns of Natural Selection

Genomic diversity across CPB’s geographical range was scanned for evidence of natural selection by identifying outlier SNPs in comparisons of population differentiation and SNPs that were highly correlated to environmental predictor variables. Population differentiation tests identified 0.37% of all SNPs as outliers (i.e. 65,815 out of 17,599,906, with a false discovery rate, or FDR, of 0.01%). A total of ∼32% of the outlier SNPs could be assigned to 8,760 known genes (**Additional File 1**: **Table S4** and **Table S5**; gene list provided in **Additional File 2**). Of these genes, 336 were linked to candidate insecticide resistance genes, including 205 genes involved in detoxification pathways, 91 target-sites, and 40 genes involved in cuticular development. The well-known voltage-sensitive sodium channel gene (LDEC011942) that provides knockdown resistance to pyrethroids was included among the target-site genes. Based on a gene set enrichment analysis, over-represented gene ontology terms were linked to insecticide resistance and/or stress (**Additional File 1**: **Fig. S17**). For biological processes, GO terms included oxidation-reduction process and response to oxidative stress (and multiple nested terms), among others. For cellular components, terms linked to insecticide resistance included voltage-gated sodium channel and acetylcholine-gated channel complexes, presynaptic active zone and synapse, and integral component of the membrane. Among the molecular functions, terms such as heme and zinc ion binding (including iron ion binding), extracellular ligand-gated ion channel activity (including voltage-gated sodium channel and acetylcholine receptor activity), glutathione transferase activity, ABC transporter activity via the term ATPase activity coupled to transmembrane movement, and peroxidase activity (including CYP monooxygenase activity) were associated with insecticide resistance.

To test for patterns of natural selection driven by environmental factors, we also employed gene environment association analysis to examine five different, ecologically-relevant predictors of natural selection on the genome: latitude, elevation, precipitation, minimum temperature in the coldest month and potato land cover. Only 0.02% of the analyzed SNPs (4,098 out of 17,599,906 SNPs, with an FDR of 0.01%) were significantly associated with at least one environmental variable (**Additional File 1**: **Table S6** and **Fig. S18**; gene list provided in **Additional File 2**). A total of 67.6% of the SNPs were associated with precipitation, 15.5% with latitude, 10.7% with potato land cover, 2.8% with elevation, and 3.7% with temperature. Of all significant SNPs, 29% were found in 816 known genes, including 42 resistance-related genes (28 involved in metabolic detoxification mechanisms, 3 in cuticle development, and 11 target-site genes; **Additional File 1**: **Table S7**). Based on a gene set enrichment analysis, over-represented gene ontology terms were associated with insecticide resistance and/or stress (**Additional File 1**: **Fig. S19**). Among the biological processes, terms included chemical synaptic transmission, oxidation-reduction process, proteolysis, defense response, and DNA repair. Among the cellular components, terms included synapse and presynaptic active zone, as well as integral component of the membrane. Among the molecular functions, terms included heme and iron ion binding, carboxylic ester hydrolase activity, extracellular ligand-gated ion channel activity, oxidoreductase activity (including monooxygenase activity), and ubiquitin binding.

The outlier-based and environmental-association genome scans leverage different models to detect selection, but a comparison of the results shows that a total of 557 genes (see **Additional File 2**) were shared in both tests, and 29 of these are candidate insecticide resistance genes (**Table 2**). The largest group represents xenobiotic detoxification genes, with nine ABC transporters, seven CYP genes, two esterase genes, one MFS gene and one GST gene represented. Target-site genes included four voltage-dependent channel genes and two nicotinic acetylcholine receptors, and three cuticle genes overlap in both tests. Gene set enrichment analysis of significant genes identified as overlapping in the two genome scan tests (**Additional File 1**: **Fig. S20**) showed enrichment of gene ontology terms associated with insecticide resistance and stress. Among biological processes, terms included chemical synaptic transmission, oxidation-reduction process, proteolysis, DNA repair, and chloride transport, transmembrane transport, and ion transport. Terms associated with cellular components included integral component of membrane, synapse, and presynaptic active zone. Terms associated with molecular functions included pathways such as heme and iron ion binding, ATPase activity coupled to transmembrane movement, GABA and G-protein coupled receptor activity, and monooxygenase and oxidoreductase activity.

**Table 2.**
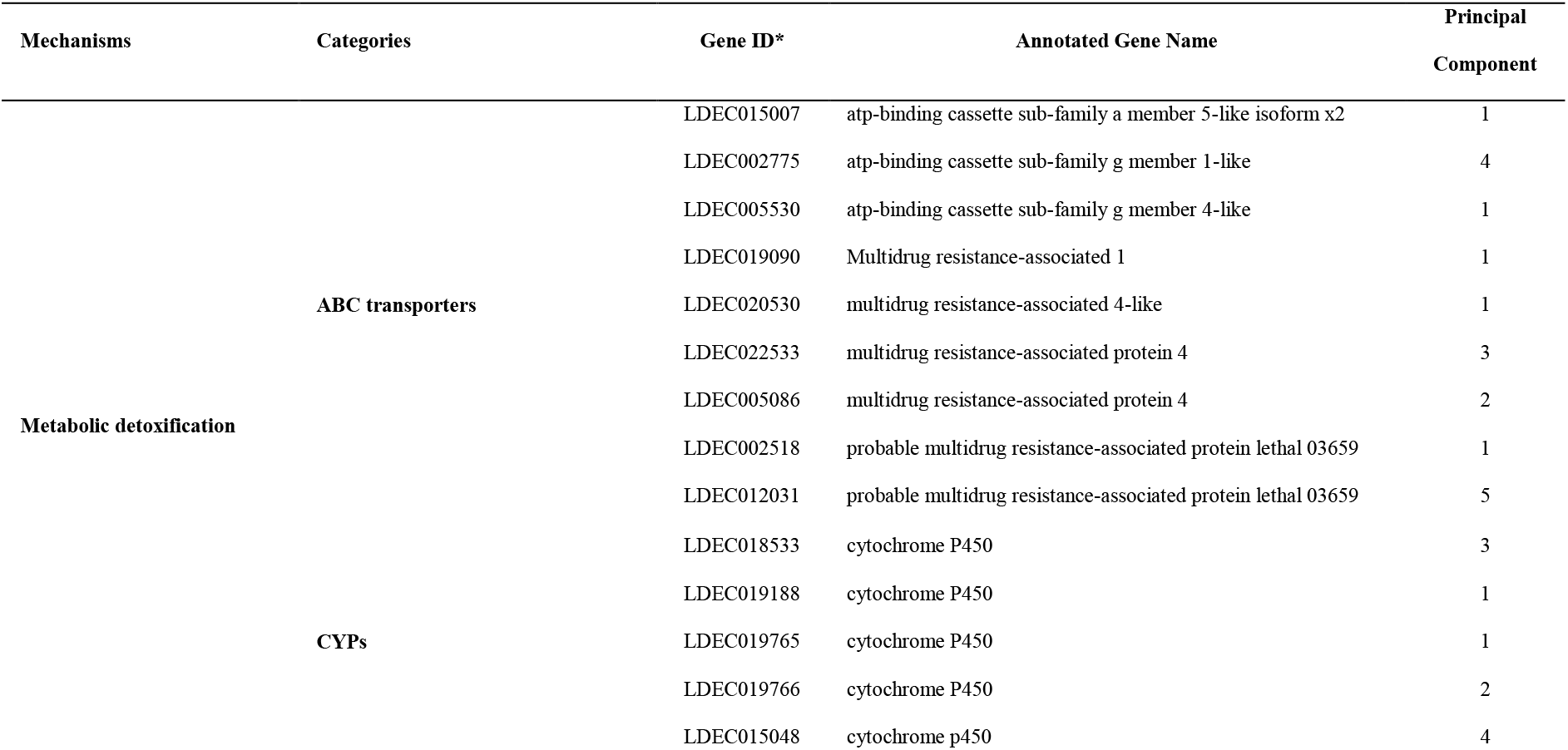
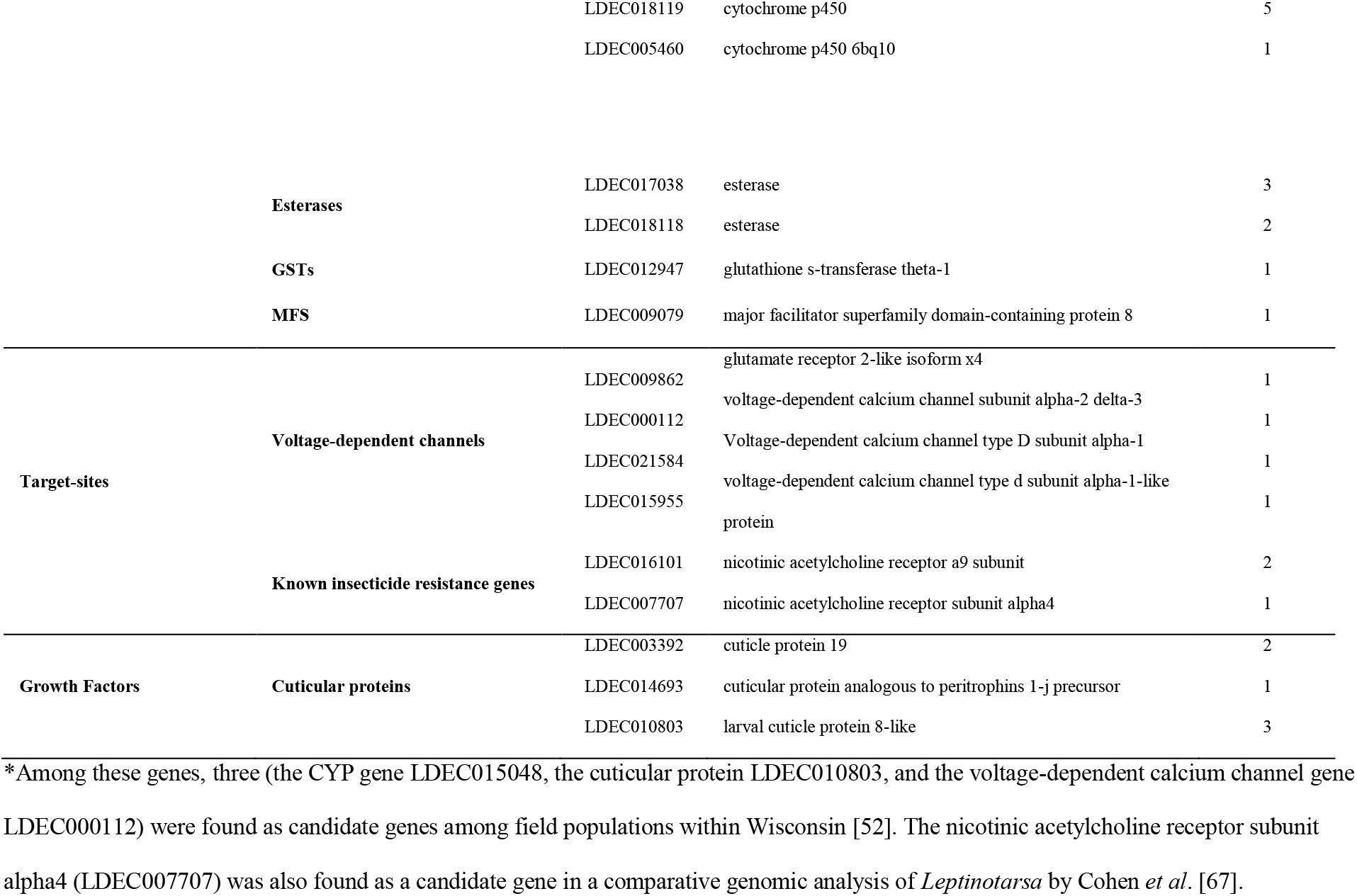
Candidate resistance genes identified in both PCAdapt and LFMM. The loading of each gene on a principal component is indicated (see **Additional File 1: Fig. S9**).

### Local Adaptation to Insecticides

To examine the geographical occurrence of selection events, we employed a haplotype-based approach that examines shifts in haplotype frequency along branches of a population tree. Due to the fragmented nature of the reference genome, we examined haplotype frequencies on the longest 95 genomic scaffolds (encompassing ∼21% of the genome, all >1 Mb). Our analyses show that 1.1% (72,386 SNPs), 0.01% (7,826 SNPs) and 0.16e^-3^% (1,106 SNPs) of the markers were significant at α = 0.01, 0.001 and 0.0001, respectively (see list in **Additional File 2**). SNPs were grouped into regions, where each region was separated by at least 1 Kb up- and downstream. This resulted in 1,169 selection regions at α = 0.01, 140 regions at α = 0.001, and 24 regions at α = 0.0001 (**Additional File 1**: **Table S8**). Excluding one extremely long region (35 Kb in length), the average length of the most significant haplotypes (α = 0.0001) was 3.1 Kb; **Additional File 1: Fig. S21**). On average (across α levels and branch association thresholds), these regions in the genome were repeatedly selected in multiple populations (on average, in 6.06 branches of the population tree, and only 4.87% of these selection events were singularly associated with one population; **Additional File 1: Table S9**). These singular regions tended to be short (1.1 Kb on average). Out of all selected regions (α = 0.01), 319 were found in 224 genes and only 3.8% were singular (**Additional File 1: Fig. S22**). Recalling that this only represents ∼21% of the genome, 24 regions comprising 16 genes were candidate insecticide resistance-associated genes, including six ABC transporters, two esterases, one olfactory receptor, one nicotinic acetylcholine receptor, two genes associated with glutamate pathways and four growth factors (**Additional File 1: Table S10**).

Most haplotype-based selection events in the candidate insecticide resistance genes (19 out of 24 regions) were less than 1 Kb long (**Additional File 1: Fig. S23**) and present (22 out of 24 regions) on nine to eleven branches of the population tree (**Fig. 3**; **Additional File 1: Fig. S24**). Selection at these candidate regions (19 out of 24) were shared between Western and Eastern lineages (**Additional File 1: Fig. S25**), which were the most genetically distinct and geographically isolated populations. Furthermore, several of these candidate genes (LDEC004355, LDEC005089, and LDEC002775) appeared to have multiple regions under selection, with population-specific patterns. The observed patterns at insecticide resistance candidates suggests that repeated selection on the same set of protein-coding genes is prevalent among populations. Over 150 genes (68.5%) identified in the haplotype-based test were also identified in the outlier test (**Additional File 1: Fig. S26**), including 11 of the 16 candidate insecticide resistance genes. Seven of these were ABC transporters, and notably three were significant in all selection tests (the ABC subfamily C/multidrug associated gene LDEC002518, and two ABC subfamily G genes, LDEC002775 and LDEC005530). The other ABC genes included: the subfamily F gene LDEC004565, and three additional multidrug associated genes LDEC005089, LDEC003183, and LDEC002116. The remaining genes included one target site gene, the acetylcholine receptor β subunit (LDEC002850), one cuticle protein (LDEC003397), one transient receptor potential (TRP) gene (LDEC003216), and one odorant binding gene (LDEC003898).

**Figure 3.**
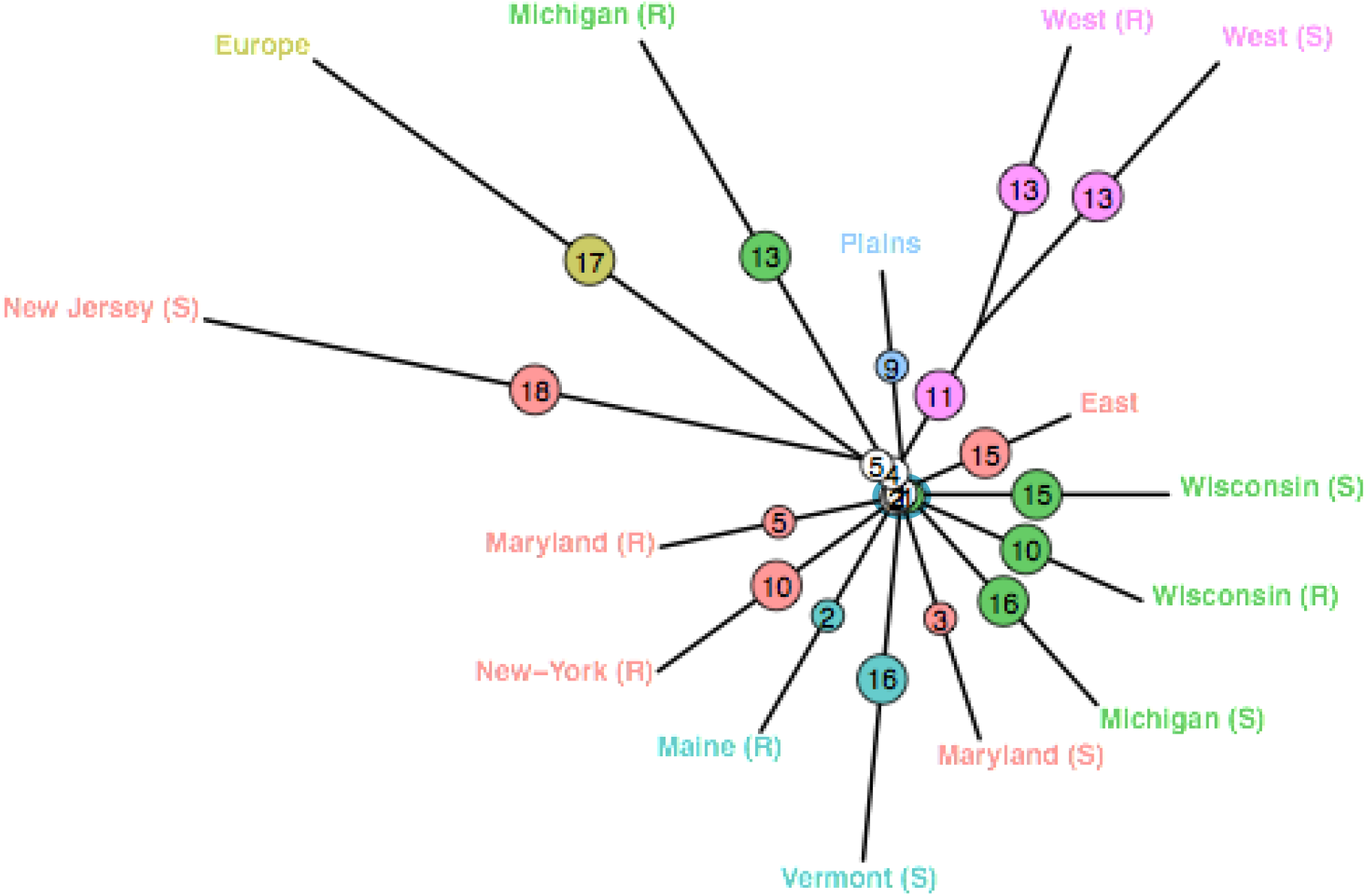
Population tree showing the distribution of 24 resistance-associated selection events identified with hapFLK in the first 95 genomic scaffolds. Colors refer to geographical location. Internal branches show few selection events (one or two events in four branches, no selection event in five branches).

We examined gene expression profiles to test whether patterns of gene regulatory evolution were shared among geographical regions for a subset of previously published CPB population samples (**Additional File 1: Table S11**). For quality control, we first assessed whether differences in experimental design influenced the expression of candidate insecticide resistance genes (see detailed results in **Additional File 1)**. Based on these comparisons, we determined that regional population differences could be compared for adults from field populations irrespective of generation sampled, but lab reared larvae needed to be compared separately. For the larval comparison, we removed samples representing an insecticide induction treatment, focusing our analysis on constitutive differences in gene expression. These comparisons showed strong geographical differences in overall gene expression profiles (**Additional File 1: Fig. S27** and **Fig. S28**). Focusing on significantly differentially expressed candidate insecticide resistance genes, local populations showed divergent patterns of constitutive upregulation among populations (**Fig. 4** and **Additional File 1: Fig. S29**; see gene list in **Additional File 2**). Seven differentially expressed candidate insecticide resistance genes were found among the adults and 84 among larvae (**Additional File 1: Fig. S29** and **Table S12**), with only one esterase (LDEC019310) and one ABC transporter (LDEC004154) common to both datasets. A gene set enrichment analysis of the differentially expressed gene list in among-population comparisons of adults (**Additional File 1: Fig. S30**) showed enrichment of terms associated with insecticide detoxification and/or stress, including heme and ion iron binding, oxidoreductase activity and dioxygenase activity, proteolysis, transport, defense response, and integral component of the membrane. Among larvae (**Additional File 1: Fig. S31**), there was enrichment of terms associated with regulatory changes to gene networks underlying insecticide detoxification or stress, such as glutathione transferase activity, hexachlorocyclohexane metabolism, oxidoreductase and monooxygenase activity, gap junction channel activity, proteolysis, substrate-specific transmembrane transporter activity, heme and iron ion binding, innate immune response, and integral component of the membrane.

**Figure 4.**
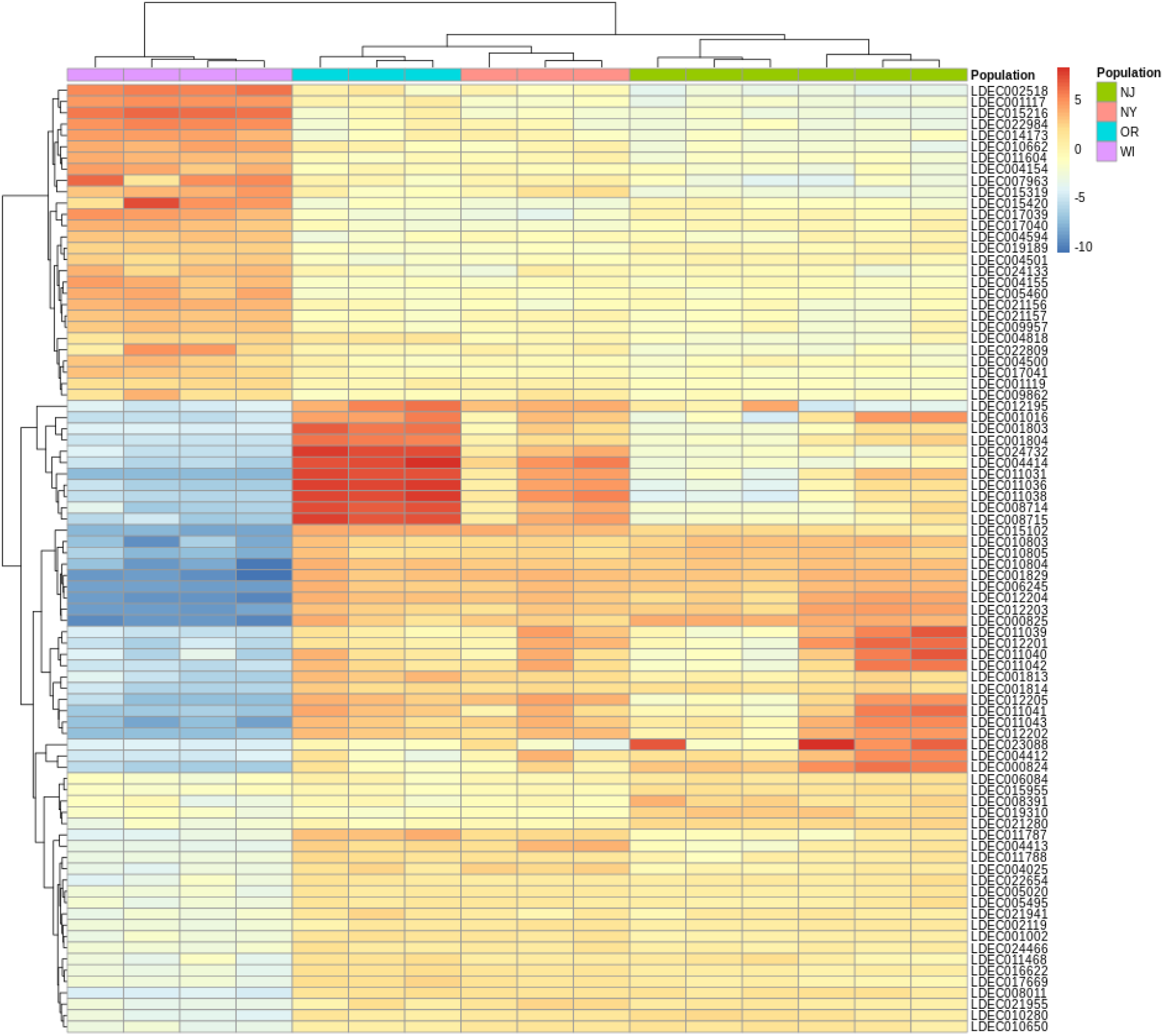
Gene expression heatmap among four populations of CPB larvae, showing divergent constitutive expression of 84 differentially expressed candidate insecticide resistance genes. Colors of expression levels correspond to log-fold change. See **Table S15** for the functional annotation of these genes.

Although 26 transcription factors were significantly differentially expressed among the four larval populations (**Additional File 1: Fig. S32** and **Table S13**), they are not known to be associated with detoxification pathways. These results do not support rapid evolution of regulatory genes, as different genes, and in some cases different molecular pathways, are favored in each regional population. Gene set enrichment analysis of the overlapping set of significant genes in larvae and adult differential expression analysis gene sets (**Additional File 1: Fig. S33**) showed shared regulatory changes in gene networks linked to insecticide detoxification and stress, such as oxidoreductase and dioxygenase activity, heme and iron ion binding, proteolysis and integral component of membrane. The shared enrichment of lipid metabolism might also be related to insecticide detoxification (through an interaction with oxidation-reduction or membrane-transport processes), rather than metabolism per se.

Finally, we examined a global intersection of gene ontology terms that were enriched in both the genome scans and differential expression datasets (**Additional File 1: Fig. S34**). Six gene ontology terms were identified, each of which has been related to insecticide resistance in prior studies: biological process terms include oxidation-reduction process and proteolysis, while cellular components include integral component of the membrane, and molecular functions include oxidoreductase activity acting on paired donors with incorporation or reduction of molecular oxygen, heme binding, and iron ion binding.

## Discussion

Population genomics is increasingly providing insight into the evolutionary mechanisms that give rise to super-pests and holds promise for improving pest management practices [8]. Our study provides the first comprehensive genomic and geographical assessment of genome-wide patterns of genetic variation for a super-pest in the center of its origin. Whole genome variation demonstrates that the Colorado potato beetle (CPB) is geographically structured, including among pest populations, and corroborates evidence from microsatellite markers that CPB pest populations are most closely related to populations in the Great Plains instead of Mexico [46, 59, 60]. In evaluating the competing (but not mutually-exclusive) mechanisms of rapid evolution, our data suggest that different loci are selected across growing regions, and among different populations within regions, with statistical tests providing a consistent pattern of repeated evolution at many candidate insecticide resistance genes. Below we discuss in turn the evidence that supports polygenic evolution from standing genetic variation, selection on *de novo* mutation, and rapid regulatory evolution. We close by discussing the pest management consequences of these modes of rapid evolution.

### Evidence for repeated selection on standing genetic variation

Population clustering analyses demonstrate that regional CPB pest populations (western US, eastern US, multiple lineages in the Midwestern US, and Europe) are genetically distinct, with D-statistics suggesting limited ongoing gene flow. This alone supports previous observations that insecticide resistance evolves locally among CPB populations [43, 51, 52, 55]. However, genome scans, using both outlier-based and environmental association-based methods, provide clear signatures that adaptation occurs repeatedly across different populations of CPB. Focusing on candidate genes of insecticide resistance, the considerable overlap among the two methods suggests different genes are selected in different populations (see PC loadings in **Table 2**). Similarly, candidate insecticide resistance genes are constitutively upregulated, but in distinct patterns in different populations (**Fig. 4**). Selected genes also encompass multiple resistance mechanisms, including metabolic detoxification, target site resistance, and cuticular proteins, which is broadly supported by gene set enrichment analyses across multiple tests (**Additional File 1: Fig. S34**). These results are consistent with previous genetic studies that have documented some of the same genes (see **Table 2; Additional File 2**) or mechanisms in CPB resistance [43, 61–64]. We also identify compelling new candidates, such as the multiple ABC transporters, as well as acetylcholine receptor β subunit (LDEC002850), which are likely linked to neonicotinoid resistance phenotypes [65, 66]. Although our analysis focuses on pesticide resistance candidates, we note that other interesting genes emerged from our study. In particular, an octopamine receptor (LDEC006841), which was identified as a gene associated with pest behavior in a comparative genomics analysis of CPB [67], is recovered as a significant target in contrasts of Plains and pest populations in both *LFMM* and *PCAdapt*.

The observed patterns of repeated local adaptation are most consistent with polygenic evolution from standing genetic variation. Using a more sophisticated haplotype-based method (focusing on ∼21% of the genome), we found 24 highly significant (*p <* 0.0001) haplotype blocks suggestive of selective sweeps. Only one selected region exceeded 4 Kb in length (35 Kb), while the remaining regions averaged 3.1 Kb in length. These shorter sweep lengths are similar to those found in at least one other insect pest, *Spodoptera frugiperda* [on average, sweeps are 4.1 Kb: 68], but much smaller than the strong selective sweeps found in other prominent cases of insecticide resistance [18, 69, 70]. Sweep lengths typically scale inversely with the rate of recombination as a product of increasing effective population size, so we might have expected sweep lengths similar to those found in insects with large population size (typically > 10 Kb in *D. melanogaster*) [71]. Instead, nearly all strong candidate selective events from *hapFLK* are more consistent with soft sweeps that recur in multiple geographically distant populations. Most importantly, the sweep regions in CPB occur on multiple branches of the population tree (on average, 8.16 branches). For soft sweeps to be a reasonable mechanism in the evolution of insecticide resistance, however, populations must maintain high levels of genomic variation. The host-shift to potato, coupled with agroecosystem invasion over a broad geographical scale, has been presumed to reduce genetic variation in CPB [47, 72]. However, despite having recently invaded agroecosystems, CPB pest populations exhibit only modest reductions in nucleotide diversity (though the loss of private alleles is more pronounced), and SMC++ reconstructions show recent population expansion. Dense population sampling of CPB at a landscape scale in the Midwestern U.S. also supports high levels of standing variation at a regional level [52].

Early population genetic research on insecticide resistance by J.F. Crow suggested that resistance evolution arises from polygenic standing variation [7]. Quantitative variation in insecticide tolerance, among populations and individuals within populations, is frequently observed, even under controlled laboratory conditions [73, 74]. Increasingly, genomics-based analyses are documenting how multiple genes contribute to quantitative resistance phenotypes [16, 75–77]. A combination of new mutations and recruitment of standing genetic variation probably occurs in many cases of adaptation, as it is evident that phenotypic traits are typically quantitative in nature (influenced by multiple loci), large-effect mutations often require compensatory fitness changes, and the most likely outcome of evolution in theoretical models is polygenic adaptation [78]. At the same time, the action of multiple genes and their regulatory elements can be difficult to detect using population genomics methods, as it results in modest changes in allele frequencies (under soft or partial selective sweeps) [79–81]. Further noise is added because tests of selection are prone to false positives [82]. For example, *hapFLK* tends to show a high false positive rate under population models with continuous migration or in those that experience strong bottlenecks [83, 84]. However, while chance false positives might contribute to our observed patterns, it is unlikely that so many would occur in candidate insecticide resistance genes. Our results are in broad support with the emerging view that polygenic architecture is common in insecticide resistance [10, 24, 85, 86]. Certainly of course, expanded and more continuous population sampling of CPB will be needed to improve support for this mode of selection and the complex genetic architecture underlying insecticide resistance.

### Selection on de novo mutation: is CPB mutation limited?

Selection on novel mutations could lead to repeated evolution of resistance in pests with intrinsically large population sizes (much greater than >>10^6^) that are not considered mutation-limited [29, 30]. Analyses of nucleotide diversity in protein-coding genes is known to range widely in animal species and is most strongly correlated with reproductive strategy, with highly fecund (r-selected) species like CPB having the greatest diversity [87]. Schoville *et al*. [58] found evidence for a high rate of polymorphism in protein-coding regions (nucleotide diversity, π, was ∼0.01), suggesting a high level of standing genetic diversity relative to other insects. Furthermore, CPB shows a higher rate of positive selection and greater levels of standing variation compared to other species in the genus *Leptinotarsa* [88]. In this genome-wide resequencing dataset, we identify an average nucleotide diversity of 0.005 within CPB, irrespective of pest or non-pest status across different geographical regions. While this rate of nucleotide diversity remains high relative to most vertebrates (median 0.0025), it is not exceptional among arthropods [median 0.0125 all sites, 0.00204 synonymous sites; 89]. Despite its super-pest status, CPB nucleotide diversity falls within the range of insect species (0.0023-0.0288). Among Coleoptera, CPB falls within the range of nucleotide diversity found in the bark beetles *Dendroctonus ponderosae* and *D. brevicornis* [0.0023 and 0.008, respectively; 90, 91] and is similar to the horned scarab beetle *Onthophagus taurus* [0.0056, 92]. Lepidopteran pest genomes are more polymorphic; five species of *Helicoverpa* range in nucleotide diversity from 0.004 to 0.010 in autosomal regions, with the super-pest *H. armigera* as the most polymorphic [93], while two sympatric stains of the pest species *Spodoptera frugiperda* range from 0.043 to 0.044 [94] and the closely related pest *S. litura* has nucleotide diversity as high as 0.016 [95].

These results suggest that CPB is not exceptional in terms of standing genetic diversity and the potential for rapid evolution. However, there is an upper limit on nucleotide diversity in species with large effective population sizes (e.g*. Drosophila melanogaster*), as nucleotide variation at neutral sites is removed as a result of selection on nearby linked sites [96]. In fact, patterns of reduced variation in species with large effective population sizes might reflect selection from recurrent adaptive mutation [97]. On the other hand, there is considerable debate about this interpretation, as reduced nucleotide diversity might alternatively arise from purifying selection acting on nearly neutral sites in the form of background selection [98, 99]. Distinguishing among these alternatives will require direct estimates of genome-wide mutation and recombination in CPB, in addition to improved sampling, as it is not yet clear that CPB is mutation-limited.

### Are key regulatory shifts contributing to resistance evolution

Overexpression of multiple CYP, GST and ABC transporter genes is often associated with insecticide resistance and several studies have shown that trans-acting transcription factors may simultaneously regulate the expression of these targets [100–103]. Known trans-acting transcription factors involved in the xenobiotic detoxification pathway include *CncC*, *Maf-S*, *AhR*, *ARNT*, and *Met*. *CncC* forms a heterodimer with *Maf-S* to regulate multiple detoxification loci in many insects [104], including the highly resistant Long Island population of CPB [63] and deltamethrin resistant populations of the beetle *Tribolium castaneum* [105]. However, in comparing gene expression profiles of CPB populations throughout several growing regions, we found that expression levels of insecticide detoxification genes vary across populations, suggesting that varied transcriptional patterns are most-likely achieved through cis-regulatory evolution [106, 107]. In a similar analysis, comparison of transcriptomic profiles of *Anopheles gambiae* across Africa revealed the recruitment of many population-specific candidate insecticide resistance genes [108], suggesting cis-regulatory evolution of these pathways may be common among insect pests. Our data don’t support the role of a single master-regulatory switch driving insecticide resistance, as we see no evidence for differential expression of key transacting transcription factors despite comparing insecticide resistant and susceptible populations. Furthermore, the role of different genes in metabolic resistance in CPB has been demonstrated by RNAi experiments where knockdown of different upregulated CYP genes restores susceptibility in different pesticide resistant CPB populations [109, 110]. Altogether, our results suggest that a simple upstream shift in *Cap-n-collar* expression is not sufficient to explain all cases of metabolic resistance in CPB and that, instead, additional cis-regulatory changes are required to account for the heterogeneity and diversity of resistance pattern among populations [111]. This is consistent with widespread evidence that cis-regulatory evolution is more common in adaptive evolution, while trans-acting gene regulation is typically constrained by strong stabilizing selection [112, 113].

### Implications for pest management and novel control tactics

Current resistance management strategies assume that the evolution of resistance is a rare event, caused by simple (single-gene) mutations [114], thereby ignoring the importance of alternative mechanisms involving multiple loci [115]. Resistance management models also assume that resistance is involved in fitness tradeoffs [116], a rationale underlying high dose/refuge strategies where gene flow from susceptible ‘refuge’ populations into insecticide-treated fields delays resistance evolution [117]. However, it is increasingly difficult to understand the rate of pesticide resistance evolution using conventional models of single, large-effect mutations in conferring a resistance phenotype [118]. Rates of insecticide resistance evolution in CPB are among the highest observed in agricultural pests [6] and often lead to failure unless implemented in an integrated pest management framework [119]. From our results, polygenic evolution from standing variation appears best explains this pattern, although we note alternative mechanisms were not investigated and could contribute to rapid evolutionary change. Notably, recent work in CPB has shown that changes in DNA methylation patterns might drive transgenerational epigenetic mechanisms of regulatory evolution that lead to pesticide resistance [120, 121].

As society faces the challenge of global food security, there is a prevailing view that insect pests have won the arms race involving conventional chemical pesticide control [3]. Novel chemical modes of action are needed to avoid target-site resistance, yet there is a high cost to such efforts, both in terms of development costs and environmental impacts. In addition, the widespread emergence of cross-resistance, especially through metabolic detoxification, suggests novel modes of action may have limited efficacy and durability [122]. Although gene drives have been raised as promising novel control tactics [123, 124], most recent work in in CPB has focused on gene-targeted insecticides via the RNAi pathway [125]. Gene knock-down via RNAi could allow for highly effective, species-specific management if multiple genes are targeted simultaneously, and such products are currently under development [126]. How do genome-wide population genetic data shed light on RNAi implementation? Drawing on the mechanisms of evolution described in this paper, where standing variation is substantial, the likelihood of resistance evolution to RNAi might be high unless population-specific approaches are developed. One target site mutagenesis experiment in CPB has shown that mismatch rate of 3% or less still allows for effective gene target suppression [127], but clearly some CPB target genes would be problematic. Additionally, alternative pathways of RNAi resistance might emerge. For example, experiments with cell lines of CPB have shown that resistance evolving from mutations altering the uptake and transport of dsRNA [128]. CPB is known to utilize both *sid-1* transmembrane channel-mediated uptake and clathrin-mediated endocytosis in processing dsRNA [129], suggesting that there are multiple targets for resistance evolution in the RNAi pathway. Comparison of dsRNA efficacy among European CPB populations also suggested variation in the RNAi pathway itself (involving the multiple homologs of *dicer*, *argonaut*, and *staufen*) was more likely to evolve than at target site loci [130]. However, knockdown experiments in other Coleopteran pests have shown that loss of function of genes in the RNAi pathway impair development and reduce reproductive fitness [131], thus the inherent trade-off in resistance may be too great. One interesting RNAi resistance pathway involves selection on gut nuclease activity that alters the sensitivity of CPB to RNAi [132]. In other insects, such as Lepidopteran pests [133], a single nuclease is responsible for dsRNA tolerance. Though clearly dsRNA provides a novel mode of action for controlling CPB pests, the propensity to draw on reservoirs of standing genetic variation to rapidly evolve suggests that multiple mechanisms of resistance are likely to occur.

## Conclusions

Understanding the molecular mechanisms underlying pesticide adaptation has become increasingly important because of the widespread occurrence of the “pesticide treadmill” phenomenon in agricultural pests [134], wherein the repeated and escalating use of pesticides, and the search for new chemistries [119], is required to keep pace with pest evolution. We provide clear evidence that polygenic resistance drawn from standing variation could explain how insects rapidly overcome multiple classes of pesticides [135]. While the importance of polygenic evolution from standing genetic variation remains mostly theoretical in the insecticide resistance literature and has proven challenging to identify in empirical case studies [8], polygenic resistance has also been broadly implicated in the evolution of herbicide resistance in agricultural weeds [85] and antibiotic resistance in bacterial pathogens [136]. Here we provide evidence that CPB evolves insecticide resistance repeatedly across agricultural regions, and often at the same loci. An important future goal will be to understand how polygenic resistance phenotypes spread among local pest populations, in order to refine integrated pest management practices to maintain the efficacy and sustainability of novel control techniques.

## Methods

### Study Design and Aim

We collected a geographically dispersed set of 88 samples was selected to maximize information about genomic differentiation across the range of CPB (**Additional File 1: Fig. S1** and **Table S1**). An additional 10 samples comprising nine species of *Leptinotarsa* were also collected for relevant information on outgroup variation and possible sources of hybridization. Within the 88 CPB samples, we sampled six geographically proximate pairs of resistant (R) and susceptible (S) populations, 5 beetles per R/S site: Maine (R) and Vermont (S), New York (R) and New Jersey (S), Maryland (R and S), Michigan (R and S), Wisconsin (R and S), Oregon (R and S). The resistance status of beetles was ascertained by topical exposure to an insecticide (imidacloprid) or from published records at those sites (see **Additional File 1** for detailed methods). However, all pest populations have potentially evolved resistance to other insecticides, as insecticide use was widespread starting in the late 1940s. As each diploid individual represents N=2 genomes, our sample size exceeds the requirements for most population genomic tests and allows for accurate estimation of frequencies for all but the most rare (and therefore, presumably, less important) alleles in key potato growing regions of the United States [137].

### Genomic Resequencing, Quality Control and Variant Calling

High quality genomic DNA was isolated from adult beetle thoracic muscle tissue using DNeasy Blood & Tissue kits (Qiagen) and then submitted to the University of Wisconsin-Madison Biotechnology Center. Libraries were sequenced using paired-end, 125bp sequencing on a HiSeq2500 sequencer (see **Additional File 1** for detailed methods). We predetermined sequencing effort to yield >6x average coverage for each of our CPB genomes, a quantity sufficient to identify SNPs with reasonable accuracy [138].

Each sample was demultiplexed prior to downstream analysis, and we followed GATK’s “Best Practices” guidelines (https://software.broadinstitute.org/gatk/best-practices/). Using the *L. decemlineata* reference genome v1.0 [GCA_000500325.1; 58], we aligned demultiplexed reads using BWA v0.7.101 [139], and converted SAM files to BAM format using SAMTOOLS v1.3.12 [139]. We generated one uBAM file (i.e., unmapped BAM file) per forward-reverse pair of the fastq raw reads using FastqToSam and then marked Illumina adapters with MarkIlluminaAdapters, both functions available from PICARD v2.2.4 (https://github.com/broadinstitute/picard). We then reverted BAM files to fastq format with PICARD’s SamToFastq, aligned the new fastq files to the reference genome with the BWA-mem algorithm and merged all alignments into one BAM file per sample with PICARD’s MergeBamAlignment tool. We marked PCR and optical duplicates using PICARD’s MarkDuplicates tool, but some of our samples were sequenced on multiple sequencer lanes. For these samples, we marked duplicates first at the lane level (i.e., per replicate), then at the sample level (merging duplicates into a unique BAM output). Finally, we realigned reads around insertions and deletions with GATK’s RealignerTargetCreator and IndelRealigner tools. In order to assess the quality of our BAM files, we used GATK’s Flagstat and DepthOfCoverage tools. Among our 88 CPB samples, three samples (two susceptible samples from Oregon: CPBWGS_59 and CPBWGS_63, and one susceptible sample from Vermont: CPBWGS_93) had few successfully mapped reads and were removed (**Additional File 1**: **Fig. S2** and **Table S1**).

Genotyping was split into two steps: per-individual variant calling, followed by joint genotyping. Variant calling was conducted with GATK’s *HaplotypeCaller* tool, which generates a likelihood score for all reference sites (-ERC GVCF option), including non-variant sites. Two different joint genotyping procedures were performed: one excluding non-CPB samples (”CPB” dataset; N=85), and one including non-CPB samples, but keeping only one susceptible and one resistant sample (chosen at random) for populations from Oregon, Wisconsin, Michigan, Maryland, New-Jersey/New-York and Vermont/Maine (”*Leptinotarsa*” dataset; N=50; **Additional File 1: Table S2)**. For joint genotyping, we employed variant quality score recalibration (VQSR) using a training dataset. VQSR is based on applying machine learning algorithms and clustering methods to examine the overlap of the raw call set and a training dataset. It is composed of two steps: 1. *ApplyRecalibration* describes the multi-dimensional annotation profile of variants and calculates (for each variant in both datasets) a new, well-calibrated quality score called VQSLOD (for “variant quality score log-odds”). 2. *ApplyRecalibration* uses VQSLOD to apply a new cutoff to retain only high likelihood variants from the call set, based on a proportion of the variants in the training set that are present in the call set (e.g. 99.9% to enhance sensitivity or 90% to enhance specificity). As this approach requires a well-validated, independent dataset to be used as a training set, we used 41,454 SNPs generated from a published genotyping-by-sequencing (GBS) experiment [52]. These data represent 188 samples from 24 Midwestern populations (**Additional File 1: Fig. S35)**, which were hard filtered for depth of coverage ≥10x, polymorphism in at least 30% of the individuals of each population, minor allele frequency ≥5%, and less than 20% missing genotypes across all individuals. We calculated VQSLODs based on the following annotations: QD, MQ, MQRankSum, ReadPosRankSum, FS, SOR, DP and InbreedingCoeff. The recalibrated score provides a continuous estimate for the probability of each variant, which can then be partitioned into quality tranches. Tranche plots for the “CPB” and “*Leptinotarsa*” datasets are based on a 90% threshold that maximizes specificity over sensitivity (**Additional File 1: Fig. S36**). Finally, we used GATK’s *VariantsToTable* tool to assess the quality of our inferred SNP dataset. We plotted the improvement in the distribution of QualByDepth (QD) following the VQSR procedure for the “CPB” dataset (**Additional File 1: Fig. S37**) using the *ggplot*2 package in R v3.6 [140, 141]. After removing sites representing mitochondrial DNA (1,756 total variant sites), our dataset contained 47,969,460 SNPs in the “CPB” dataset and 69,680,768 in the “*Leptinotarsa*” dataset. Some analyses, like demographic reconstruction, require analyses with neutrally evolving loci. To mitigate the effect of non-neutral loci in these analyses, we created an “intergenic CPB dataset” from our “CPB” dataset, by considering only SNPs located outside of known genes from the *L. decemlineata* Official Gene Set (OGS) v1.1 [58].

### Genomic Diversity

To estimate the genetic diversity of populations, we used the “CPB” dataset (*i.e*. not including Mexican samples), removed multi-allelic SNPs and those SNPs with a MAF <5% (using GATK’s *SelectVariants* tool; final SNP number = 17,599,906). For these analyses we consider paired populations (susceptible/resistant) and grouped individuals from the U.S. Plains region (Colorado, Nebraska, Kansas, Missouri, New Mexico and Texas). We estimated genome-wide nucleotide diversity (π) using a 10 Kb sliding window with VCFtools v0.1.15 [142]. We also estimated heterozygosity by calculating the inbreeding coefficient *F* for each individual, using the method of moments estimator in VCFtools. Individual estimates were then averaged per population, keeping susceptible and resistant individuals separate. Finally, we used the “*singletons”* function in VCFtools to calculate the number of singletons and private doubletons for each sample.

### Evolutionary Divergence

In order to reconstruct the evolutionary origins of CPB populations, we conducted a phylogenomic analysis of the “*Leptinotarsa*” dataset. This dataset comprised 48 samples, including 10 *L. decemlineata* samples from Mexico, 28 *L. decemlineata* from the U.S. and Europe, and 10 samples of closely-related *Leptinotarsa* species. Since the dataset was quite large, we used only the first 100 scaffolds comprising 16,519,065 SNPs. We used SNPhylo v.20160204 [143] to construct a phylogeny, after pruning the SNPs for linkage disequilibrium (LD). In order to ensure the relative independence of the SNPs used in the analysis, we tested several values of the LD threshold parameter and selected 0.5 for downstream analyses, which resulted in 35,838 SNPs. The SNP data were concatenated and then aligned using MUSCLE v3.8.31 [144], and a maximum likelihood phylogeny was estimated using DNAML in PHYLIP v3.6 [145]. Support values for nodes in the tree were determined by bootstrap resampling 100 times.

To estimate population structure, we examined both the “CPB” dataset and the “intergenic CPB” dataset, but the results were biologically consistent. We used three approaches: classical F_ST_ estimates between pairs of populations, principal components analysis using *PCAdapt* [146, 147] and ancestry analysis with *sNMF* (Frichot and François 2016). For the F_ST_ analyses, we consider paired populations (susceptible/resistant) and Plains individuals (Colorado, Nebraska, Kansas, Missouri, New Mexico and Texas). All estimations of Weir and Cockerham’s mean weighted fixation indices (F_ST_) were done using 10 Kb windows in VCFtools [142]. For the PCAdapt analysis, we assessed the number of principal components using Cattell’s rule [148]. We plotted the percentage of variance explained by the first 20 principal components in a screeplot and examined the point of inflection in the plot, above which additional terms provide diminishing returns in terms of explained variance. For *sNMF*, we inferred individual patterns of ancestry by estimating ancestral population allele frequencies and admixture coefficients using the R package *LEA* [149]. We first converted our VCF files to PLINK’s ped format using VCFtools, then to the geno file format using *LEA*’s *ped2geno* tool. We implemented 10 runs per k value and combined the different runs with CLUMPAK (http://clumpak.tau.ac.il/). We plotted cross-entropy values to assess the number of k values.

Finally, to test for possible admixture among ancestral populations during the invasion of CPB into agroecosystems, we assessed evidence of gene flow (i) from Mexican CPB populations and (ii) from other *Leptinotarsa* species, into the pest lineage. We used *Dsuite* [150] to calculate the genome-wide *D*-statistic (*D*) on allele frequency data, estimating the strength of introgression based on the ABBA/BABA test [151]. *D* ranges from zero (no introgression) to one (complete introgression), and is calculated using a set of three focal populations or taxa (P1, P2), P3) with one additional outgroup. For the first test (i), we only considered populations with sample sizes ≥ 5, grouping susceptible and resistant samples from Vermont/Maine, Maryland, Wisconsin and Oregon as F_ST_ between these sub-populations was negligible (**Additional File 1: Table S5**), grouping samples from the “Plains” region (N=7), and grouping all CPB samples from Mexico (N = 10). We tested every possible focal trio of populations, retaining the lowest *D*-statistic for every given trio (*D_min_*; a conservative estimate of *D*). We assessed whether *D* was significantly different from zero by calculating a *p*-value based on jackknifing. We analyzed the complete biallelic SNP dataset with the two *L. juncta* samples as outgroups (the results were almost identical when using *L. undecemlineata* instead; data not shown). For the second test (ii), we included the nine other *Leptinotarsa* species using the same dataset from the SNPhylo analysis, comprising the 100 first scaffolds and 16,519,065 SNPs. The dataset contained one susceptible and one resistant sample for each of the six paired populations. In order to analyze a balanced dataset, we limited the “Plains” population to the samples from Colorado. We also created two Mexican populations, representing the two distinct Mexican clades recovered in our phylogeny: “Mexico_City” (containing two samples) and “Mexico_South” (containing the sample from Oaxaca and the one from Guerrero; **Additional File 1: Table S1**). We used *L. lineolata* as the outgroup, as it was recovered as the most basal and distantly-related taxon to the U.S. CPB clade.

### Demographic Analysis

To reconstruct the population history of CPB, we used the CPB intergenic SNP dataset and employed two coalescent approaches: the stairway plot method [152] and SMC++ [153]. Demographic reconstructions rely on an accurate estimate of the mutation rate. Most estimates of nuclear mutation rate in insects fall into the range of 2 × 10^-9^ to 7 × 10^-9^ substitutions per site per generation [154, 155]. As there is no genome-wide mutation rate estimate for CPB or related beetles, we chose to use a mutation rate of 2.1 × 10^-9^, estimated recently in the non-biting midge [156]. We set the generation time of 0.5/year (*i.e*. 2 generations per year) for all our samples. The Stairway plot approach relies on the calculation of the expected composite likelihood of a given site frequency spectrum (SFS), which reduces the computational burden of inferring population parameters. It is also suitable for estimating recent population histories with low coverage genomic data. We analyzed the resistant and susceptible paired populations, both separately and pooled together, and also considered a pooled sample from the Plains (Colorado, Nebraska, Kansas, Missouri, New Mexico and Texas), East (Florida, Tennessee, North Carolina, Virginia, Kentucky and Ohio) and Europe (Italy, Russia). For each population, we estimated the folded SFS in *dadi* [157], calculated for a genome length of 678Mbp [58], and used 200 bootstraps to assess confidence intervals. We then conducted the Stairway plot analysis with default parameters and plotted the estimated median (and 95% confidence intervals) effective population size (Ne) through time.

We chose SMC++ [153] as an alternative approach, as it incorporates estimates of recombination and linkage disequilibrium (LD) in an SFS framework. Even though this method is relatively computationally efficient, long run times prevented us from using the entire CPB dataset. Due to the fragmented nature of the genome, we analyzed the longest 95 genomic scaffolds of the CPB dataset, including all intergenic and non-intergenic biallelic SNPs. This encompasses ∼21% of the genome (∼140MB out of ∼670M bp, mean length of 1.9MB and all > 1MB) and contains 6,669,259 SNPs. This subset of highly contiguous reference genomic data ensures we can accurately infer demography using the sequential Markov coalescent [158], although we note that analyses considering all >10 Kb scaffolds (covering 95% of the genome) yielded comparable results (i.e. same population sizes, same demographic events, same time-scale; results not shown).

### Selection Analyses

We used three different approaches to study genomic signatures of selection: outlier detection with *PCAdapt* [146, 147], genome-environment association with *LFMM* [159] and haplotype-based tests using *hapFLK* [84]. To detect outlier SNPs using *PCAdapt*, Mahalanobis distances were transformed into *p-values* and then the FDR was controlled by transforming the *p-values* into *q*-values and considering an FDR of 0.01% (α=0.0001). We initially filtered SNPs using a minor allele frequency (MAF) of 0.05 and a conservative setting of K=10, but the number of SNPs suggested a high rate of false positives (see **Additional File 1**). We therefore refined our filtering steps using linkage disequilibrium clumping (choosing a window size of 500 SNPs and a squared-correlation coefficient threshold of 0.2). We examined the screeplot of the principle components and, following Cattell’s rule, selected K=6 as the optimal clustering level. Finally, we adjusted the dataset by setting a more conservative MAF setting of 0.1.

As an alternative genome scan approach, we employed a genome-environmental association method using latent factor mixed models [LFMM; 159]. This test identifies SNP allele frequencies that are significantly associated with environmental predictor variables, while simultaneously modeling the confounding effect of population structure as latent factors. To account for the population structure observed in our data, we modeled k = 6 latent factors, as suggested by our *PCAdapt* and *sNMF* results. We adjusted *p*-values by an empirically-determined genomic inflation factor, while controlling the false discovery rate at 0.01%. We explored five different environmental variables: elevation, precipitation, minimum temperature in the coldest month and potato land cover. We reasoned that genes containing SNPs associated with climate variables could be related to *L. decemlineata* adaptation to northern climates during range expansion, while associations with potato land cover might reveal genes responding to selective pressures faced in potato agroecosystems, such as novel host plants, natural enemy pressure and insecticide exposure. We obtained historic, county-level potato land cover data (between 1850 and 2012), as detailed in Crossley *et al*. [160]. For latitude, elevation, precipitation and minimum temperature in the coldest month, we obtained the data from the PRISM climate group (http://prism.oregonstate.edu). For each environmental variable, we took the average value within a 75 km radius around each sampling site, using functions available in the *rgdal* and *raster* packages in R [161, 162]. For potato land cover, we summarized the average proportion of area planted with potato within a 75 km radius of each sample site [163].

Finally, to integrate information from linked SNPs in tests of selection, we used the haplotype frequency-based method *hapFLK* [84]. This method has been shown to be relatively robust to confounding effects of population structure and variable population size. It also allows selection events to be pinpointed to specific branches of the population tree. For each identified signature of selection, a local tree is re-estimated using significant SNPs, constrained by the overall topology of the population tree. Statistical significance is computed for the difference between the branch lengths estimated from the focal region and from the global tree. We analyzed the first (longest) 95 genomic scaffolds of the CPB dataset (representing ∼21% of the genome), including all biallelic SNPs. From a VCF file produced with GATK’s *SelectVariants* function, we produced a ped file and associated map file with VCFtools’ *--plink* function. In order to use the multi-point linkage disequilibrium model, *hapFLK* needs the number of haplotype clusters (K) to be specified and a population tree. We compared *hapFLK* results for different K values, ultimately selecting K=20 to minimize imputation errors (see **Additional File 1** for detailed methods; **Additional File 1: Fig. S38**). The population tree was estimated from a kinship matrix (**Additional File 1**: **Fig. S39**). We standardized *hapFLK* values and computed corresponding *p-values* from a standard normal distribution, examining results at three nominal levels (α = 0.01, 0.001 and 0.0001; **Additional File 1: Fig. S40**). We grouped significant SNPs into selected regions, where each region was separated by at least 1 Kb up- and downstream.

### Candidate Genes and Gene Network Analysis

For each selection test, we obtained functional information for candidate SNPs using manual annotation of the OGS supplemented by Blast2GO annotations [164], which have been previously published [52, 58]. To develop a list of candidate insecticide resistance genes, Crossley *et al.* [52] identified 664 genes associated with processes or functions potentially linked to known mechanisms of insecticide resistance (**Additional File 1: Table S14**): metabolic detoxification including cytochrome p450s (CYPs), esterases, Glutathione S-transferases (GSTs) and ATP-binding cassette (ABC) transporters [165]; target-site insensitivity including most of the modes of actions classified by the *Insecticide Resistance Action Committee* (IRAC; http://www.irac-online.org/modes-of-action/) such as TRPC channels, sodium and calcium channels, glutamate receptors, acetylcholine receptors, etc.; and reduced cuticular penetration, including genes involved in chitin production or cuticle development.

One frequent concern in analyses of large datasets is the inclusion of false positives (Type I error) resulting from the large number of statistical tests. Frequently, this is resolved by adjusting *p*-values to more conservative values, for example by implementing multiple testing recalibrations such as Bonferroni or the Benjamini-Hochberg false discovery rate. However, in functional genomic studies, gene lists provide objective hypotheses that can be easily assessed in follow-up studies, while removing them based on statistical criteria can sometimes be challenging [166]. Although we employ stringent statistical criteria in all tests, we additionally leverage information by comparing gene lists from different tests. As we expect that both regulatory and structural changes might lead to genetic adaptation, we tested for over-representation of specific gene networks in selection tests, gene expression analyses, and combinations of both approaches. Given multiple data sources, overlap in gene identity and function provides a measure of support for repeated evolution and polygenic adaptation. We curated gene ontology terms associated with significant genes in lists from genome-wide selection tests and differential expression tests, and used a one-sided hypergeometric Fisher’s Exact test [167, 168] to test for over-representation (enrichment) of gene ontology terms, with *p*-value < 0.05 used as the statistical significance threshold. To further refine this analysis, we used REVIGO [169], a clustering algorithm that relies on semantic similarity measures, to summarize the list of enriched gene ontology terms. Gene ontology terms associated with biological processes, cellular components, and molecular function were separately clustered using the simRel score for functional similarity, allowing for redundancy in similar terms up to a value of 0.7 before removal, and then compared to the UniProt database to find the percentage of genes annotated with each gene ontology term. The results were visualized using a CIRGO plot [170]. To provide a context for interpreting these results, we used our list of candidate genes (**Additional File 1: Table S14**) to generate a CIRGO plot of gene ontology terms associated with insecticide resistance (**Additional File 1: Fig. S41**). Major biological processes include response to insecticide/response to oxidative stress, endocytosis, glycerolipid metabolism, sensory perception, and DNA integration. Major cellular components include the plasma membrane/integral component of the membrane and transcription factor complex. Finally, major molecular functions include metallocarboxypeptidase activity, monooxygenase activity, acetylcholine binding, lipid binding, DNA polymerase binding, tetracycline transporter activity, chromatin binding, chitin binding, and structural component of cuticle.

### Gene Expression Analyses

In order to test for regulatory evolution, we compared gene expression data from RNA sequencing (RNAseq) experiments across the geographical range of CPB, including original data from the Plains region (a Colorado population) and previously published pest CPB population samples (see **Additional File 1: Table S11**). The Colorado population was raised under greenhouse conditions on potato plants (∼25°C, 16:8 light:dark cycle), but represents the first generation derived from wild-caught adults feeding on *Solanum rostratum*. RNAseq studies are recognized as robust estimators of whole-genome gene expression profiles [171] that are highly responsive to experimental conditions [172]. As the original experiments varied in their design, we conducted a set of initial analyses to determine if sampling issues might bias geographical comparisons. First, we assessed whether sampling of an overwintering (post-diapause) population versus a summer (non-diapausing) generation at the same field in Wisconsin altered gene expression patterns (1^st^ generation versus 2^nd^ generation). Second, we examined whether direct exposure to imidacloprid was necessary to induce insecticide resistance gene expression responses, by comparing a set of lab-reared individuals from Wisconsin, Oregon and Long Island populations (control versus an imidacloprid-induction treatment). Third, we compared samples of larvae and adults from a susceptible population in Wisconsin. Based on these comparisons, we determined that regional population differences could be compared for adults from field collected populations irrespective of generation sampled, but lab reared larvae needed to be compared separately. We compared constitutive levels of gene expression in six adult populations: Colorado (CO), Wisconsin (WI), Michigan (MI), New York (NY), New Jersey (NJ), and eastern Canada (CAN). We compared constitutive levels of gene expression in four larval populations: Oregon (OR), Wisconsin (WI), New York (NY), and New Jersey (NJ). As some of the adult samples were sequenced as pair-end and single-end reads, we analyzed only the first read of a set of pair-end samples representing CO and WI.

We aligned short read data from each sample to the *L. decemlineata* reference genome using HISAT2 [173]. SAMTOOLS was used to convert *sam* files to *bam* files. Read counts per gene per sample were generated using the function *featureCounts* available in the *Rsubread* package [174], with reference to the *L. decemlineata* OGS. Using the resulting counts, we evaluated evidence for differential gene expression for each region using DESEQ2 [175]. DESEQ2 first estimates the dispersion among a set of replicated samples and then the logarithmic fold change of transcript counts among sample groups. It then employs a generalized linear model based on the negative binomial distribution of transcript counts and a binomial Wald statistic to test for differences among experimental contrasts. We first trimmed the read count matrix to remove genes with less than five reads and then conducted the differential expression analysis. We retained differentially expressed genes if read counts were > 2-fold and the significance level α = 5% was reached. We then adjusted the false discovery rate to 1% level, using a Benjamini-Hochberg correction [176]. Heatmaps of differentially expressed genes were generated in R using the *pheatmap* package [177]. As described previously, we curated gene ontology terms associated with significant genes and used a one-sided hypergeometric Fisher’s Exact test to test for over-representation (enrichment) of gene ontology terms, with a *p*-value <0.05 used as the statistical significance threshold.

## Declarations

The authors wish to thank the reviewers and editorial staff for their assistance with our manuscript. No ethics approval was required for this research and the authors declare no competing interests. All genomic data have been made publicly available at NCBI (Bioproject PRJNA580490). Funding was provided by a USDA NIFA AFRI Exploratory Grant (2015-67030-23495), a USDA-National Potato Council award (58-5090-7-073) and two Hatch Awards (WIS02004 and VT-H02010), in addition to support from Wisconsin Potato and Vegetable Growers Association. The authors thank the University of Wisconsin Biotechnology Center DNA Sequencing Facility for providing facilities and services. Finally, we thank Margarethe Brummerman for help in sampling *Leptinotarsa* species, and an extensive network of collectors that helped us to collect beetles in the U.S, Mexico, and Europe.

## Supplementary Information

**Additional File 1** (.pdf): Supplementary Methods and Results. Contains detailed methodology and results from analytical methods, including Supplemental Figures S1-S44 and Supplemental Tables S1-S15.

**Additional File 2** (.xlsx): Significant genes from selection tests and gene expression tests.

