## Additional File 1 for "Genome resequencing reveals rapid, repeated evolution in the Colorado potato beetle, *Leptinotarsa decemlineata*"

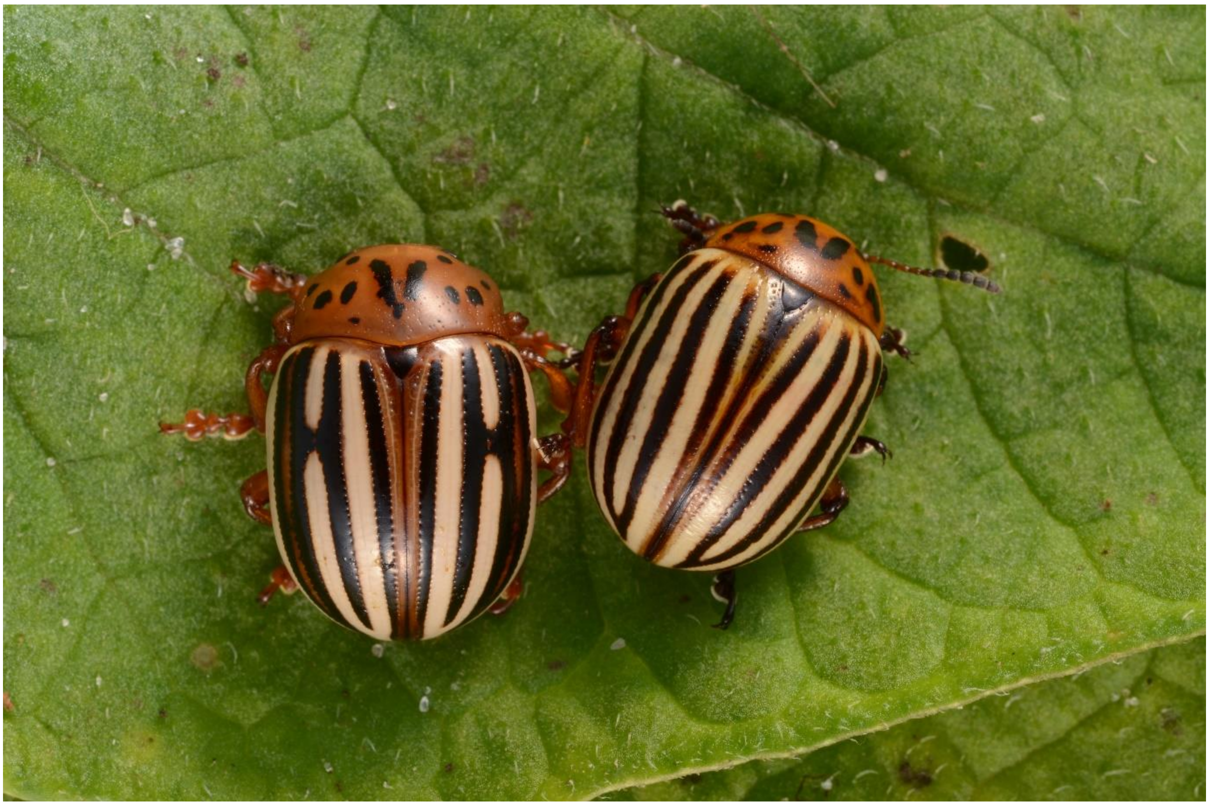

**Figure S1**. Photo of *Leptinotarsa juncta* (left) and *L. decemlineata* (right) adults on a potato plant in Gainesville, Florida. Photo credit: Lyle Bass, April 26, 2015.

### Supplementary Results

#### Sequencing Results

Our resequencing project was designed (**Fig. S2**) to test for geographical variation and the evolution of insecticide resistance in Colorado potato beetle (CPB), *Leptinotarsa decemlineata*. Among our 88 CPB samples (**Table S1**), three samples (two susceptible samples from Oregon: CPBWGS_59 and CPBWGS_63, and one susceptible sample from Vermont: CPBWGS_93) had few successfully mapped reads (8.5%, 9.2% and 13% respectively; see **Fig. S3**), resulting in very shallow average coverage of 0.03, 0.1 and 0.04 respectively. After removing these samples, the average fraction of successfully mapped reads was 96.3±2.8% (90.3±3.4% of the reads were successfully mapped with their pair), leading to an average coverage of 4.82±0.79 reads. Note that the eight *Leptinotarsa* spp. had an average fraction of successfully mapped reads of 91.5±5.2%. Genome-wide depth of coverage was >4x for most of our Colorado potato beetle samples (see **Table S1** and **Fig. S3**).

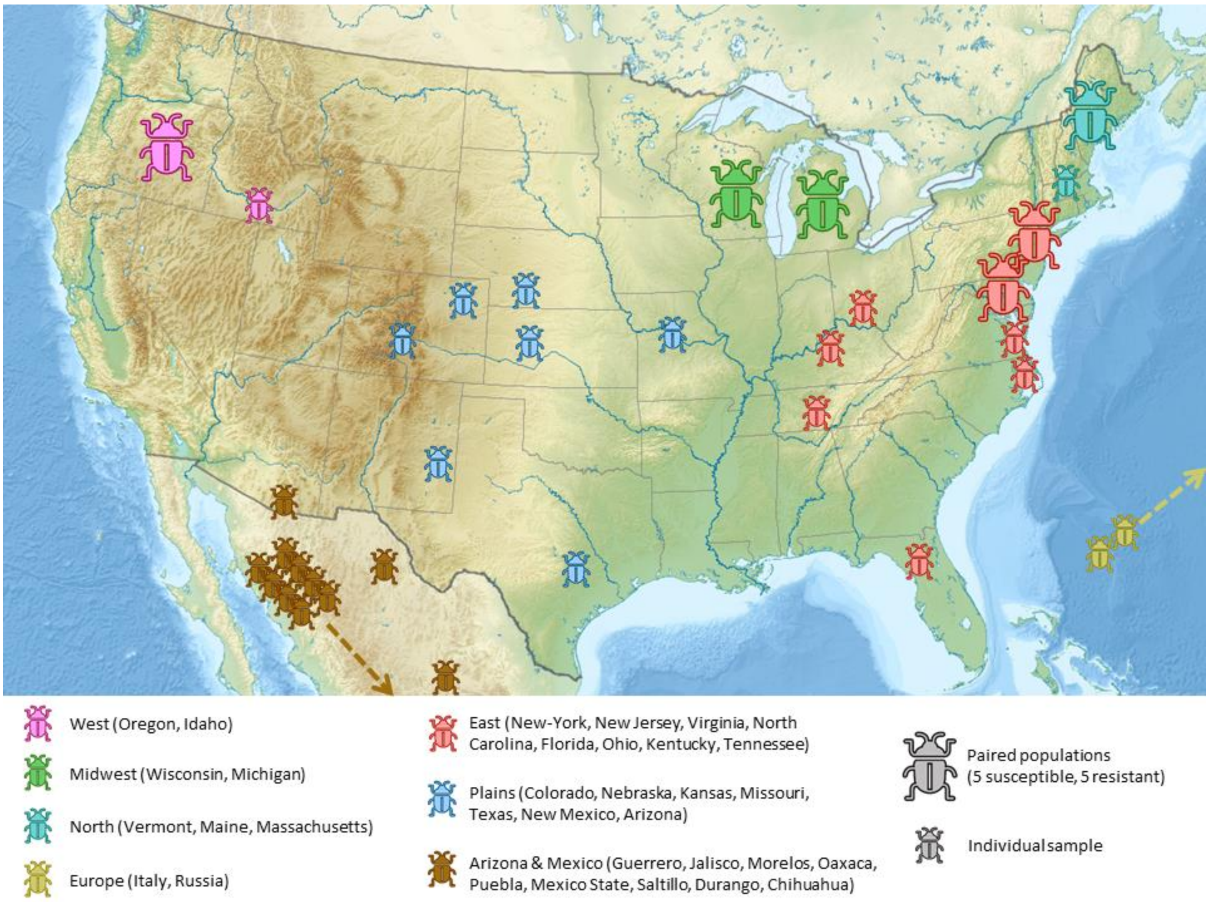

**Figure S2.** Geographical distribution of CPB samples used for genome resequencing. Dotted arrows show samples located outside the map for European and Mexican samples.

**Table S1.** Samples used for genome resequencing. Raw sequencing data and metadata are available for download under NCBI Bioproject PRJNA580490.

| Sample ID | Species | Sex | Location | Latitude | Longitude | Collector | Sequencing Effort | Mean Coverage | NCBI Accession |
|---|---|---|---|---|---|---|---|---|---|
| CPBWGS_01 | *L. peninsularis* | Female | Pima Co., Arizona | 31.824 | -110.927 | Margarethe Brummerman | 20X | 17.0 | SRR10388401 |
| CPBWGS_02 | *L. juncta* | Female | College Park, Maryland | 38.987 | -76.941 | David Hawthorne | 20X | 20.5 | SRR10388400 |
| CPBWGS_03 | *L. juncta* | Female | Gainesville, Florida | 29.65 | -82.32 | Lyle Buss | 20X | 20.3 | SRR10388389 |
| CPBWGS_04 | *L. rubiginosa* | Female | Watson Lake in Prescott, Yavapay Co., Arizona | 34.58 | -112.42 | Margarethe Brummerman | 20X | 19.3 | SRR10388378 |
| CPBWGS_05 | *L. haldemani* | Female | Arizona, Yavapai Co., S. of Prescott | 34.423 | -112.567 | Margarethe Brummerman | 20X | 22.6 | SRR10388367 |
| CPBWGS_06 | *L. texana* | Female | Highway 649, Starr Co., Texas | 26.18729 | -98.37932 | Sean Schoville | 20X | 19.0 | SRR10388356 |
| CPBWGS_07 | *L. defecta* | Female | Mission, Hidalgo Co., Texas | 26.18729 | -98.37932 | Sean Schoville | 20X | 19.2 | SRR10388345 |
| CPBWGS_08 | *L. undecemlineata* | Female | Chiapas, Mexico | 16.560583 | -92.470153 | Bruce Ferguson and Helda Morales | 20X | 21.0 | SRR10388334 |
| CPBWGS_09 | *L. tumamoca* | Female | Pima Co., Arizona | 31.824 | -110.927 | Margarethe Brummerman | 20X | 17.8 | SRR10388323 |
| CPBWGS_10 | *L. lineolata* | Female | Pima Co., Arizona | 31.805 | -111.003 | Margarethe Brummerman | 20X | 19.0 | SRR10388312 |
| CPBWGS_11 | *L. decemlineata* | Female | Copper Canyon S. of Huachuca Mtn., Cochise Co., Arizona | 31.339269 | -110.32869 | Margarethe Brummerman | 6-7X | 2.35 | SRR10388399 |
| CPBWGS_12 | *L. decemlineata* | Female | Hampshire Co., Massachusetts | 42.379461 | -72.596339 | Vic Izzo | 6-7X | 4.02 | SRR10388398 |
| CPBWGS_13 | *L. decemlineata* | Male | Accomack Co., Virginia | 37.584617 | -75.818217 | Vic Izzo | 6-7X | 5.02 | SRR10388397 |
| CPBWGS_14 | *L. decemlineata* | Male | Ellis Co., Kansas | 38.861594 | -99.333558 | Vic Izzo | 6-7X | 6.49 | SRR10388396 |
| CPBWGS_15 | *L. decemlineata* | Male | Yuma, Yuma Co., Colorado | 39.602511 | -102.325383 | Andrew Norton | 6-7X | 4.87 | SRR10388395 |
| CPBWGS_16 | *L. decemlineata* | Male | Burleson Co., Texas | 30.5715 | -96.658367 | Yolanda Chen | 6-7X | 4.87 | SRR10388394 |
| CPBWGS_17 | *L. decemlineata* | Male | Boone Co., Missouri | 38.892703 | -92.200942 | Vic Izzo | 6-7X | 4.99 | SRR10388393 |
| CPBWGS_18 | *L. decemlineata* | Male | Guerrero, Mexico | 18.327083 | -99.499433 | Vic Izzo | 6-7X | 4.48 | SRR10388392 |
| CPBWGS_19 | *L. decemlineata* | Male | Jalisco, Mexicao | 20.90025 | -103.994917 | Vic Izzo | 6-7X | 5.79 | SRR10388391 |
| CPBWGS_20 | *L. decemlineata* | Male | Morelos, Mexico | 18.77845 | -99.299917 | Vic Izzo | 6-7X | 3.95 | SRR10388390 |
| CPBWGS_21 | *L. decemlineata* | Male | Oaxaca, Mexico | 16.919417 | -96.368833 | Vic Izzo | 6-7X | 4.15 | SRR10388388 |
| CPBWGS_22 | *L. decemlineata* | Female | Puebla, Mexico | 18.85595 | -97.5469 | Vic Izzo | 6-7X | 4.92 | SRR10388387 |
| CPBWGS_23 | *L. decemlineata* | Male | Texcoco, Mexico | 19.518983 | -98.88565 | Vic Izzo | 6-7X | 4.76 | SRR10388386 |
| CPBWGS_24 | *L. decemlineata* | Female | Saltillo, Mexico | 25.4383 | -100.9737 | Sergio Sanchez-Peña and Celso Avanti | 6-7X | 6.04 | SRR10388385 |
| CPBWGS_25 | *L. decemlineata* | Female | Durango, Mexico | 25.050458 | -104.489965 | Sergio Sanchez-Peña and Celso Avanti | 6-7X | 4.46 | SRR10388384 |
| CPBWGS_26 | *L. decemlineata* | Female | Chihuahua, Mexico | 28.076221 | -105.355825 | Sergio Sanchez-Peña and Celso Avanti | 6-7X | 3.19 | SRR10388383 |
| CPBWGS_27 | *L. decemlineata* | Male | Putnam Co., Tennessee | 36.1 | -85.5 | Philip Moore | 6-7X | 5.02 | SRR10388382 |
| CPBWGS_28 | *L. decemlineata* | Male | Caprock, Lea Co., New Mexico | 33.391 | -103.712 | David Hawthorne | 6-7X | 4.92 | SRR10388381 |
| CPBWGS_29 | *L. decemlineata* | Male | (On potato) Tlaxcala, Mexico | 19.594683 | -98.61425 | Vic Izzo | 6-7X | 3.61 | SRR10388380 |
| CPBWGS_30 | *L. decemlineata* | Female | Morrow, Ohio | 39.355 | -84.128 | David Hawthorne | 6-7X | 4.63 | SRR10388379 |
| CPBWGS_31 | *L. decemlineata* | Female | Lexington, Kentucky | 38.038 | -84.505 | Ric Bessin | 6-7X | 4.84 | SRR10388377 |
| CPBWGS_32 | *L. decemlineata* | Female | Vernon G James Research Center, Plymouth, North Carolina | 35.87342 | -76.660317 | Dominic Reisig | 6-7X | 5.73 | SRR10388376 |
| CPBWGS_33 | *L. decemlineata* | Male | Florida, Gainesville | 29.65 | -82.32 | Lyle Buss | 6-7X | 4.49 | SRR10388375 |
| CPBWGS_34 | *L. decemlineata* | Female | Twin Falls Co., Idaho | 42.550256 | -114.342597 | Vince Adamson, Amy Carroll, Erik Wenninger | 6-7X | 5.76 | SRR10388374 |
| CPBWGS_35 | *L. decemlineata* | Female | Vogel Canyon, Otero Co., Colorado | 37.75071 | -103.477568 | Whitney Cranshaw | 6-7X | 5.93 | SRR10388373 |
| CPBWGS_36 | *L. decemlineata* | Female | Kearney Co., Nebraska | 40.615747 | -99.137258 | Tim Stortz | 6-7X | 5.54 | SRR10388372 |
| CPBWGS_37 | *L. decemlineata* | Female | Camposampiero, Italy | 45.561119 | 11.947728 | Alessandro Grapputo | 6-7X | 5.07 | SRR10388371 |
| CPBWGS_38 | *L. decemlineata* | Male | Novosibirskaya, Russia | 55.02 | 82.979 | S.E. Tshernyshev | 6-7X | 5.45 | SRR10388370 |
| CPBWGS_39 | *L. decemlineata* | Male | (Susceptible) Arlington, Columbia Co., Wisconsin | 43.30438055 | -89.33189999 | Michael Crossley | 6-7X | 5.29 | SRR10388369 |
| CPBWGS_40 | *L. decemlineata* | Female | (Susceptible) Arlington, Columbia Co., Wisconsin | 43.30438055 | -89.33189999 | Michael Crossley | 6-7X | 5.21 | SRR10388368 |
| CPBWGS_41 | *L. decemlineata* | Female | (Susceptible) Arlington, Columbia Co., Wisconsin | 43.30438055 | -89.33189999 | Michael Crossley | 6-7X | 4.43 | SRR10388366 |
| CPBWGS_42 | *L. decemlineata* | Female | (Susceptible) Arlington, Columbia Co., Wisconsin | 43.30438055 | -89.33189999 | Michael Crossley | 6-7X | 5.19 | SRR10388365 |
| CPBWGS_43 | *L. decemlineata* | Male | (Susceptible) Arlington, Columbia Co., Wisconsin | 43.30438055 | -89.33189999 | Michael Crossley | 6-7X | 3.51 | SRR10388364 |
| CPBWGS_44 | *L. decemlineata* | Female | (Resistant) Hancock, Waushara Co., Wisconsin | 44.119753 | -89.535683 | Michael Crossley | 6-7X | 5.05 | SRR10388363 |
| CPBWGS_45 | *L. decemlineata* | Female | (Resistant) Hancock, Waushara Co., Wisconsin | 44.119753 | -89.535683 | Michael Crossley | 6-7X | 5.09 | SRR10388362 |
| CPBWGS_46 | *L. decemlineata* | Female | (Resistant) Hancock, Waushara Co., Wisconsin | 44.119753 | -89.535683 | Michael Crossley | 6-7X | 5.56 | SRR10388361 |
| CPBWGS_47 | *L. decemlineata* | Female | (Resistant) Hancock, Waushara Co., Wisconsin | 44.119753 | -89.535683 | Michael Crossley | 6-7X | 5.1 | SRR10388360 |
| CPBWGS_48 | *L. decemlineata* | Male | (Resistant) Hancock, Waushara Co., Wisconsin | 44.119753 | -89.535683 | Michael Crossley | 6-7X | 5.68 | SRR10388359 |
| CPBWGS_49 | *L. decemlineata* | Male | (Susceptible) Allegan Co., Michigan | 42.67145278 | -85.40225833 | Michael Crossley | 6-7X | 3.87 | SRR10388358 |
| CPBWGS_50 | *L. decemlineata* | Male | (Susceptible) Allegan Co., Michigan | 42.67145278 | -85.40225833 | Michael Crossley | 6-7X | 5.03 | SRR10388357 |
| CPBWGS_51 | *L. decemlineata* | Male | (Susceptible) Allegan Co., Michigan | 42.67145278 | -85.40225833 | Michael Crossley | 6-7X | 4.61 | SRR10388355 |
| CPBWGS_52 | *L. decemlineata* | Female | (Susceptible) Allegan Co., Michigan | 42.67145278 | -85.40225833 | Michael Crossley | 6-7X | 4.34 | SRR10388354 |
| CPBWGS_53 | *L. decemlineata* | Female | (Susceptible) Allegan Co., Michigan | 42.67145278 | -85.40225833 | Michael Crossley | 6-7X | 4.95 | SRR10388353 |
| CPBWGS_54 | *L. decemlineata* | Male | (Resistant) Allegan Co., Michigan | 42.70099167 | -85.87668611 | Michael Crossley | 6-7X | 3.92 | SRR10388352 |
| CPBWGS_55 | *L. decemlineata* | Female | (Resistant) Allegan Co., Michigan | 42.70099167 | -85.87668611 | Michael Crossley | 6-7X | 4.31 | SRR10388351 |

| | | | | | | | | | |
|---|---|---|---|---|---|---|---|---|---|
| CPBWGS_56 | *L. decemlineata* | Male | (Resistant) Allegan Co., Michigan | 42.70099167 | -85.87668611 | Michael Crossley | 6-7X | 5.11 | SRR10388350 |
| CPBWGS_57 | *L. decemlineata* | Female | (Resistant) Allegan Co., Michigan | 42.70099167 | -85.87668611 | Michael Crossley | 6-7X | 4.87 | SRR10388349 |
| CPBWGS_58 | *L. decemlineata* | Female | (Resistant) Allegan Co., Michigan | 42.70099167 | -85.87668611 | Michael Crossley | 6-7X | 5.16 | SRR10388348 |
| CPBWGS_59 | *L. decemlineata* | Female | (Susceptible) Hermiston, Umatilla Co., Oregon | 45.82063333 | -119.2845028 | Silvia Rondon | 6-7X | 0.03 | SRR10388347 |
| CPBWGS_60 | *L. decemlineata* | Male | (Susceptible) Hermiston, Umatilla Co., Oregon | 45.82063333 | -119.2845028 | Silvia Rondon | 6-7X | 4.01 | SRR10388346 |
| CPBWGS_61 | *L. decemlineata* | Male | (Susceptible) Hermiston, Umatilla Co., Oregon | 45.82063333 | -119.2845028 | Silvia Rondon | 6-7X | 4.57 | SRR10388344 |
| CPBWGS_62 | *L. decemlineata* | Male | (Susceptible) Hermiston, Umatilla Co., Oregon | 45.82063333 | -119.2845028 | Silvia Rondon | 6-7X | 5.46 | SRR10388343 |
| CPBWGS_63 | *L. decemlineata* | Female | (Susceptible) Hermiston, Umatilla Co., Oregon | 45.82063333 | -119.2845028 | Silvia Rondon | 6-7X | 0.1 | SRR10388342 |
| CPBWGS_64 | *L. decemlineata* | Female | (Resistant) Hermiston, Umatilla Co., Oregon | 45.85497083 | -119.5325238 | Silvia Rondon | 6-7X | 4.47 | SRR10388341 |
| CPBWGS_65 | *L. decemlineata* | Male | (Resistant) Hermiston, Umatilla Co., Oregon | 45.85497083 | -119.5325238 | Silvia Rondon | 6-7X | 4.37 | SRR10388340 |
| CPBWGS_66 | *L. decemlineata* | Male | (Resistant) Hermiston, Umatilla Co., Oregon | 45.85497083 | -119.5325238 | Silvia Rondon | 6-7X | 6.1 | SRR10388339 |
| CPBWGS_67 | *L. decemlineata* | Male | (Resistant) Hermiston, Umatilla Co., Oregon | 45.85497083 | -119.5325238 | Silvia Rondon | 6-7X | 6.01 | SRR10388338 |
| CPBWGS_68 | *L. decemlineata* | Female | (Resistant) Hermiston, Umatilla Co., Oregon | 45.85497083 | -119.5325238 | Silvia Rondon | 6-7X | 7.72 | SRR10388337 |
| CPBWGS_69 | *L. decemlineata* | Female | (Susceptible) Prince George's Co., Maryland | 38.86 | -76.78 | David Hawthorne | 6-7X | 5.04 | SRR10388336 |
| CPBWGS_70 | *L. decemlineata* | Female | (Susceptible) Prince George's Co., Maryland | 38.86 | -76.78 | David Hawthorne | 6-7X | 5.15 | SRR10388335 |
| CPBWGS_71 | *L. decemlineata* | Female | (Susceptible) Prince George's Co., Maryland | 38.86 | -76.78 | David Hawthorne | 6-7X | 5 | SRR10388333 |
| CPBWGS_72 | *L. decemlineata* | Male | (Susceptible) Prince George's Co., Maryland | 38.86 | -76.78 | David Hawthorne | 6-7X | 5.32 | SRR10388332 |
| CPBWGS_73 | *L. decemlineata* | Female | (Susceptible) Prince George's Co., Maryland | 38.86 | -76.78 | David Hawthorne | 6-7X | 5.08 | SRR10388331 |
| CPBWGS_74 | *L. decemlineata* | Female | (Resistant) Dorchester Co., Maryland | 38.59 | -75.91 | David Hawthorne | 6-7X | 5.58 | SRR10388330 |
| CPBWGS_75 | *L. decemlineata* | Female | (Resistant) Dorchester Co., Maryland | 38.59 | -75.91 | David Hawthorne | 6-7X | 4.83 | SRR10388329 |
| CPBWGS_76 | *L. decemlineata* | Female | (Resistant) Dorchester Co., Maryland | 38.59 | -75.91 | David Hawthorne | 6-7X | 4.89 | SRR10388328 |
| CPBWGS_77 | *L. decemlineata* | Male | (Resistant) Dorchester Co., Maryland | 38.59 | -75.91 | David Hawthorne | 6-7X | 5.53 | SRR10388327 |
| CPBWGS_78 | *L. decemlineata* | Female | (Resistant) Dorchester Co., Maryland | 38.59 | -75.91 | David Hawthorne | 6-7X | 5.55 | SRR10388326 |
| CPBWGS_79 | *L. decemlineata* | Female | (Susceptible) French Biolabs-USDA-New Jersey Department of Agriculture colony, West Trenton, New Jersey | 40.215 | -74.765 | French Biolabs | 6-7X | 5.28 | SRR10388325 |
| CPBWGS_80 | *L. decemlineata* | Female | (Susceptible) French Biolabs-USDA-New Jersey Department of Agriculture colony, West Trenton, New Jersey | 40.215 | -74.765 | French Biolabs | 6-7X | 4.91 | SRR10388324 |
| CPBWGS_81 | *L. decemlineata* | Male | (Susceptible) French Biolabs-USDA-New Jersey Department of Agriculture colony, West Trenton, New Jersey | 40.215 | -74.765 | French Biolabs | 6-7X | 4.26 | SRR10388322 |
| CPBWGS_82 | *L. decemlineata* | Male | (Susceptible) French Biolabs-USDA-New Jersey Department of Agriculture colony, West Trenton, New Jersey | 40.215 | -74.765 | French Biolabs | 6-7X | 5.76 | SRR10388321 |
| CPBWGS_83 | *L. decemlineata* | Male | (Susceptible) French Biolabs-USDA-New Jersey Department of Agriculture colony, West Trenton, New Jersey | 40.215 | -74.765 | French Biolabs | 6-7X | 4.04 | SRR10388320 |
| CPBWGS_84 | *L. decemlineata* | Female | (Resistant) Long Island, Suffolk Co., New York | 40.905657 | -72.752664 | Sandra Menasha | 6-7X | 3.98 | SRR10388319 |
| CPBWGS_85 | *L. decemlineata* | Female | (Resistant) Long Island, Suffolk Co., New York | 40.905657 | -72.752664 | Sandra Menasha | 6-7X | 3.83 | SRR10388318 |
| CPBWGS_86 | *L. decemlineata* | Female | (Resistant) Long Island, Suffolk Co., New York | 40.905657 | -72.752664 | Sandra Menasha | 6-7X | 4.34 | SRR10388317 |
| CPBWGS_87 | *L. decemlineata* | Female | (Resistant) Long Island, Suffolk Co., New York | 40.905657 | -72.752664 | Sandra Menasha | 6-7X | 4.26 | SRR10388316 |
| CPBWGS_88 | *L. decemlineata* | Female | (Resistant) Long Island, Suffolk Co., New York | 40.905657 | -72.752664 | Sandra Menasha | 6-7X | 4.07 | SRR10388315 |
| CPBWGS_89 | *L. decemlineata* | Female | (Susceptible) Chittenden Co., Vermont | 44.265393 | -72.960289 | Yolanda Chen | 6-7X | 4.47 | SRR10388314 |
| CPBWGS_90 | *L. decemlineata* | Male | (Susceptible) Chittenden Co., Vermont | 44.265393 | -72.960289 | Yolanda Chen | 6-7X | 3.98 | SRR10388313 |
| CPBWGS_91 | *L. decemlineata* | Female | (Susceptible) Chittenden Co., Vermont | 44.265393 | -72.960289 | Yolanda Chen | 6-7X | 3.84 | SRR10388311 |
| CPBWGS_92 | *L. decemlineata* | Female | (Susceptible) Chittenden Co., Vermont | 44.265393 | -72.960289 | Yolanda Chen | 6-7X | 4.22 | SRR10388310 |
| CPBWGS_93 | *L. decemlineata* | Male | (Susceptible) Chittenden Co., Vermont | 44.265393 | -72.960289 | Yolanda Chen | 6-7X | 0.04 | SRR10388309 |
| CPBWGS_94 | *L. decemlineata* | Female | (Resistant) Presque Isle, Aroostook Co., Maine | 46.661319 | -68.020531 | Andrei Alyhokhin | 6-7X | 3.98 | SRR10388308 |
| CPBWGS_95 | *L. decemlineata* | Female | (Resistant) Presque Isle, Aroostook Co., Maine | 46.661319 | -68.020531 | Andrei Alyhokhin | 6-7X | 4.15 | SRR10388307 |
| CPBWGS_96 | *L. decemlineata* | Male | (Resistant) Presque Isle, Aroostook Co., Maine | 46.661319 | -68.020531 | Andrei Alyhokhin | 6-7X | 5.43 | SRR10388306 |
| CPBWGS_97 | *L. decemlineata* | Female | (Resistant) Presque Isle, Aroostook Co., Maine | 46.661319 | -68.020531 | Andrei Alyhokhin | 6-7X | 4.52 | SRR10388305 |
| CPBWGS_98 | *L. decemlineata* | Female | (Resistant) Presque Isle, Aroostook Co., Maine | 46.661319 | -68.020531 | Andrei Alyhokhin | 6-7X | 4.39 | SRR10388304 |

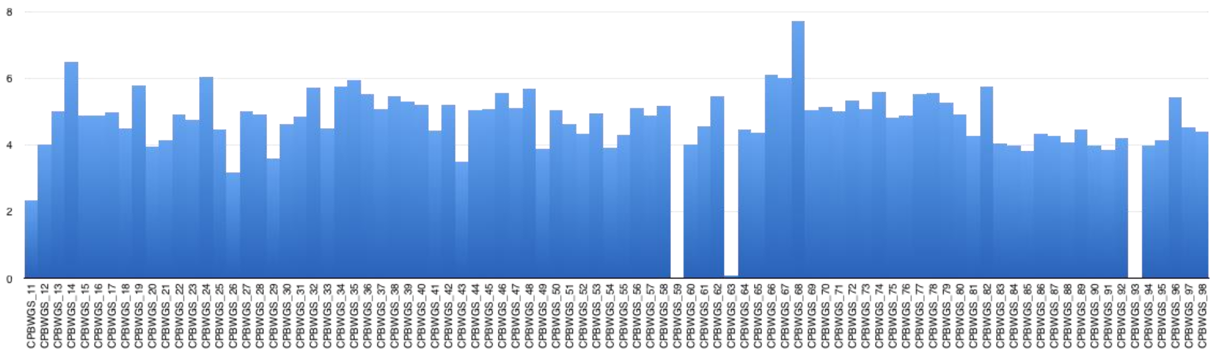

**Figure S3.** Average depth of coverage (number of reads mapped per site), across samples.

**Table S2.** Number of variants (SNPS, insertions and deletions) and SNPs within the "CPB" and "*Leptinotarsa*" datasets.

| | **CPB dataset** | ***Leptinotarsa* dataset** |
|---|---|---|
| **Total Variants** | 88 966 091 | 130 361 272 |
| **SNPs** | 76 647 868 | 109 278 144 |
| **Indels** | 12,318,223 | - |
| **Filtered SNPs (post-VQSR)** | 47,969,460 | - |
| **Intergenic SNPs only** | 30,973,249 | - |

### Genetic diversity

We estimated genome-wide nucleotide diversity ($\pi$) using a 10 Kb sliding window and found it was comparably low between susceptible and resistant pairs of populations (average $\pi$ = 0.0028 and 0.003, respectively), except for Michigan and New York / New Jersey populations (**Fig. S4**). Susceptible individuals from Michigan showed a 20% higher $\pi$ than their resistant counterpart ($\pi$ = 0.0031 vs. 0.0025, respectively). Conversely, susceptible individuals from New Jersey (a long-term laboratory population) showed a 39% lower $\pi$ than individuals from New York ($\pi$ = 0.0019 vs. 0.0031, respectively). When susceptible and resistant individuals were pooled together, estimates increased by 40%, reaching an average $\pi$ = 0.005, identical to the value obtained for the pooled Plains samples. This suggests that low $\pi$ values obtained for susceptible and resistant separately might be an artefact of small sample sizes.

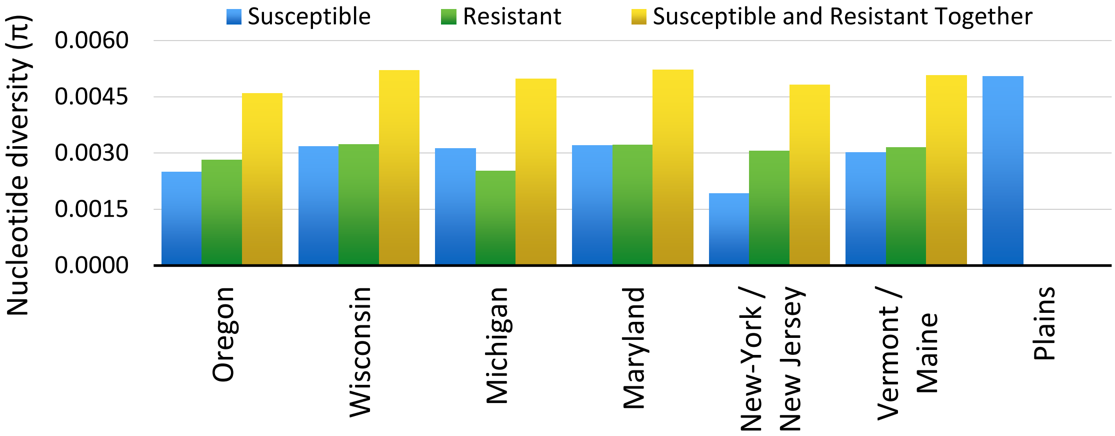

**Figure S4.** Genome-level nucleotide diversity (π) of each CPB population using a 10 Kb sliding window.

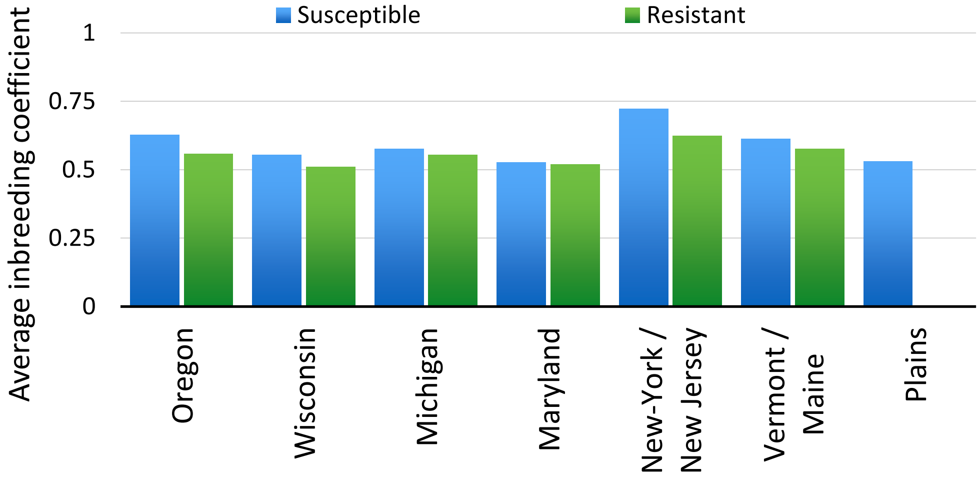

**Figure S5.** Average inbreeding coefficient for each population.

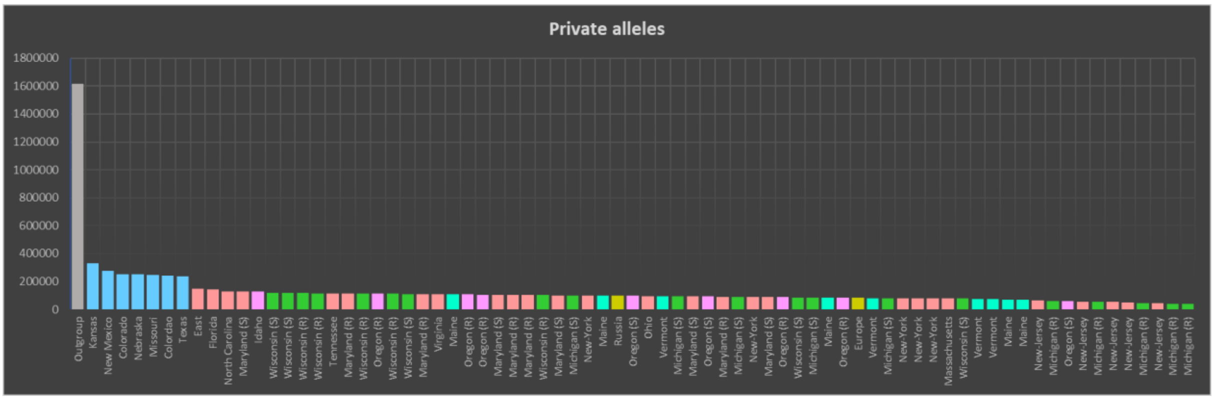

**Figure S6.** Number of singletons and private doubletons for the 75 CPB samples from U.S. and Europe, sorted from highest to lowest. The Arizona CPB sample is shown in grey. Colors correspond to geographical regions: Blue=Plains, Salmon=East, Pink=West, Green=Midwest, Turquoise=North-East, Mustard=Europe. Grey (outgroup) = Arizona

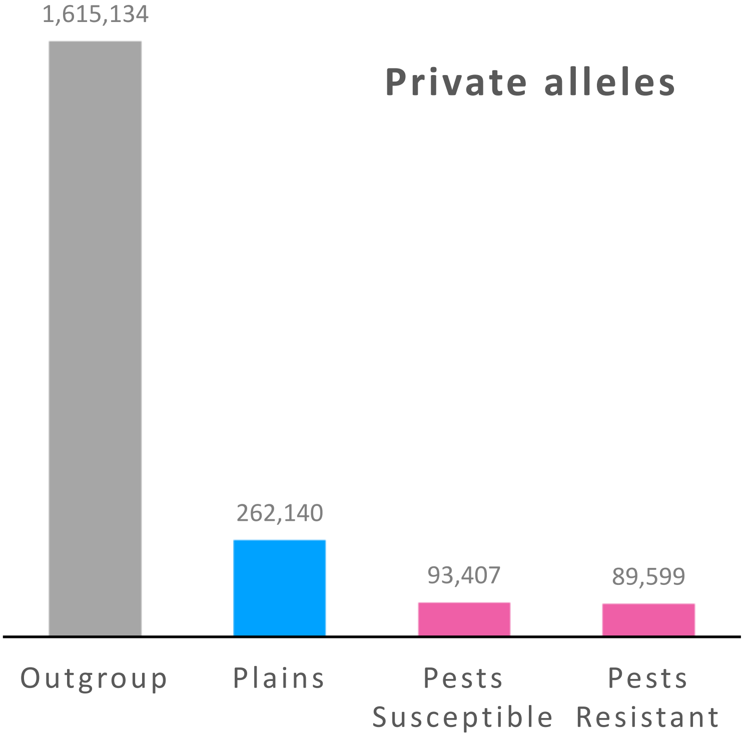

**Figure S7.** Number of private alleles (singletons + private doubletons) per pooled population. Outgroup = Arizona (N=1), Plains (Colorado, Nebraska, Kansas, Missouri, New Mexico, Texas; N=7), Pest Susceptible (N=37), Pest Resistant (N=30).

### Population structure

We assessed population structure using both the "CPB" dataset and the “intergenic CPB” dataset, but the results were biologically consistent. For the $F_{ST}$ analyses, we first calculated $F_{ST}$ between the susceptible and resistant individuals in paired samples from Oregon, Wisconsin, Michigan, Maryland, New York/New Jersey and Vermont/Maine. Only Michigan and New-York/New Jersey populations showed a $F_{ST} > 0.1$ (0.142 and 0.262, respectively), all other populations exhibited null values (**Fig. S8**). We then calculated the $F_{ST}$ between populations, pooling susceptible and resistant populations when $F_{ST} < 0.1$. Overall, $F_{ST}$ values were modest (average $F_{ST} = 0.09$), although Michigan resistant and New Jersey exhibited $F_{ST} > 0.1$ against most other populations (**Table S3**).

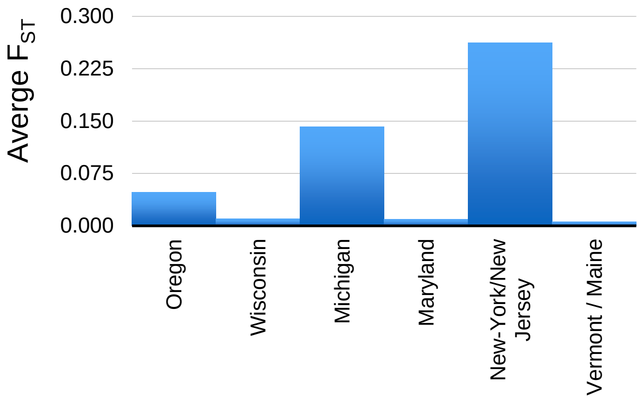

**Figure S8.** Average pairwise $F_{ST}$ between susceptible and resistant populations. The New York, New Jersey estimate may be inflated because the New Jersey population is a long-term laboratory population.

**Table S3.** Average pairwise $F_{ST}$ between populations. Bold values: $F_{ST}$ >0.1.

| | Plains | Oregon | Wisconsin | Michigan suscept. | Michigan resistant | Maryland | New-Jersey | New-York | Vermont / Maine |
|---|---|---|---|---|---|---|---|---|---|
| Plains | - | 0.075 | 0.017 | 0.021 | **0.114** | 0.019 | **0.196** | 0.024 | 0.020 |
| Oregon | - | - | 0.083 | 0.097 | **0.208** | 0.085 | **0.310** | 0.099 | 0.088 |
| Wisconsin | - | - | - | 0.003 | 0.096 | 0.008 | **0.176** | 0.011 | 0.007 |
| Michigan suscept. | - | - | - | - | - | 0.009 | **0.206** | 0.009 | 0.008 |
| Michigan resistant | - | - | - | - | - | 0.094 | **0.297** | **0.112** | 0.097 |
| Maryland | - | - | - | - | - | - | **0.172** | 0.006 | 0.005 |
| New-Jersey | - | - | - | - | - | - | - | - | **0.181** |
| New-York | - | - | - | - | - | - | - | - | 0.006 |
| Vermont / Maine | - | - | - | - | - | - | - | - | - |

For the principal component analysis, we assessed the number of principal components using Cattell's rule, examining the plot of explained variance for inflection points in the first 20 principal components. The resulting screeplots suggested significant diminishing returns beyond k=10 (**Fig. S9**), with Cattell's rule favoring values from k=5 to k=10. In an examination of the first 10 principal components (**Fig. S10**), PC1 differentiates samples from New-Jersey and Western populations (Oregon + Idaho) from the rest of the samples. PC2 confirms PC1's clustering and further differentiates Michigan-resistant population and Plains-population (samples from the South and Central U.S.: Colorado, Nebraska, Kansas, Missouri, Texas and New Mexico) from the rest of the samples. PC3 further separates the Michigan-resistant population and PC4 separates the European samples (Russia and Italy) from other samples. PC5 to PC6 highlight the Plains cluster and a Midwest cluster, but show substantial genetic variance in these groupings. PC7 to PC10 do not show biologically significant population structure.

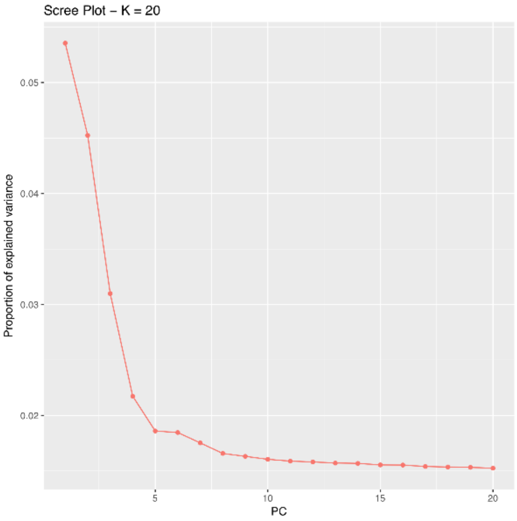

**Figure S9**. Screeplot calculated on the complete nuclear dataset and showing the decreasing proportion of explained variance by the first principal components.

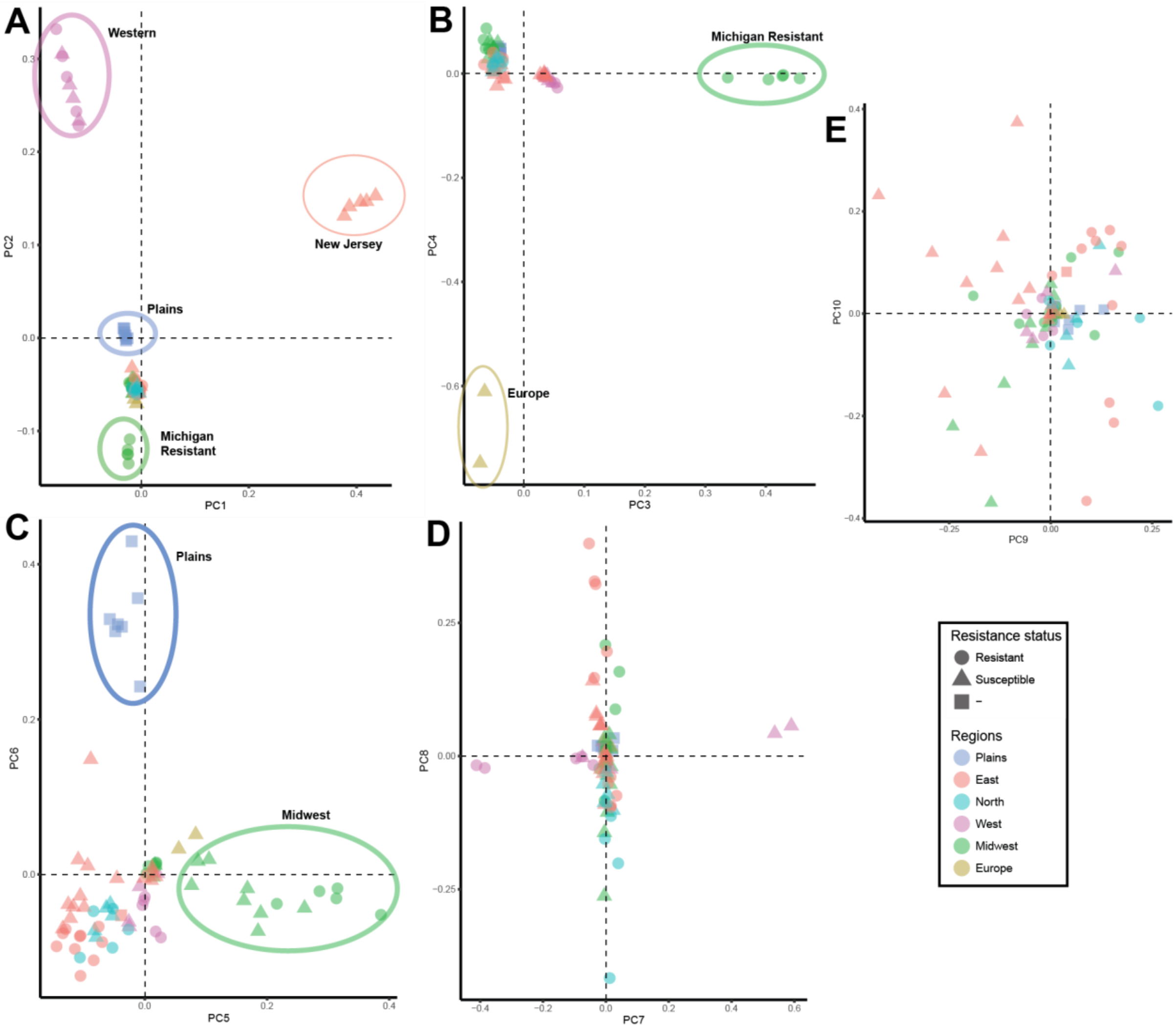

**Figure S10**. PCA plots calculated on the complete "CPB" dataset, for PC 1 to 10.

We inferred individual patterns of ancestry by estimating ancestral population allele frequencies and admixture coefficients using *sNMF*. Although the cross-entropy criterion suggested k=2 as the best number of ancestral population (**Fig.S11**), we chose to use k=6 as a clear geographical pattern of ancestral subdivision is shown in the results and the pattern of population structure is consistent with that estimated from the PCA analysis (**Fig. S12**).

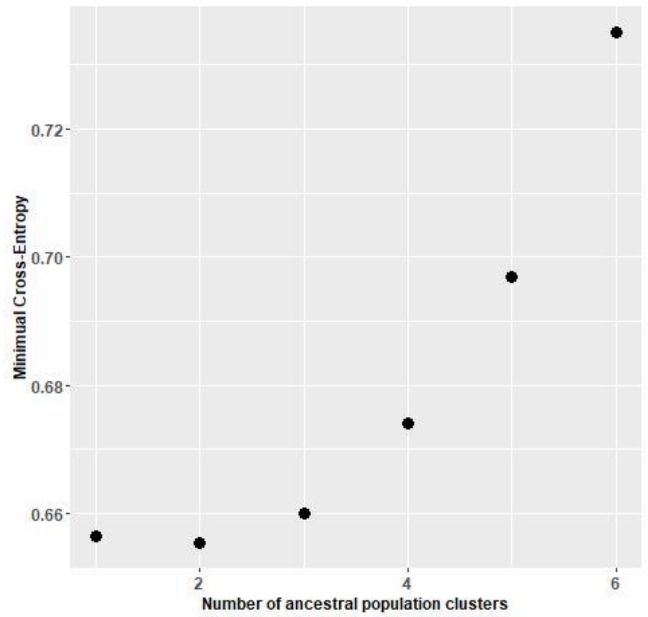

**Figure S11**. Minimal cross-entropy criterion as a function of number of ancestral population (k), for the intergenic dataset. Smaller values theoretically mean better prediction capacity.

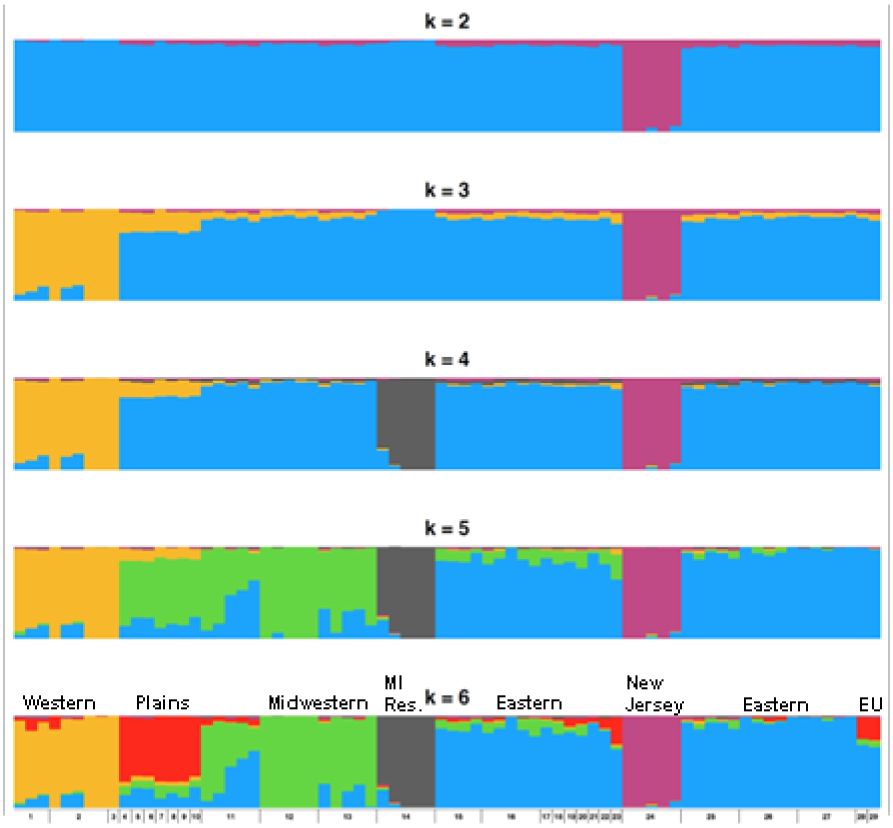

**Figure S12**. Admixture coefficients, estimated with *sNMF* on the intergenic "CPB" for k=2 to k=6. Bars are numbered according to the map in Figure 1B.

In order to test for possible admixture, we calculated the genome-wide *D*-statistic (*D*) on allele frequency data. First, we analyzed our complete biallelic SNP dataset with the two *L. juncta* samples as outgroups (the results were almost identical when using *L. undecemlineata* instead; data not shown). We found the level of introgression among CPB populations from U.S. and Mexico to be very low overall, with a maximum $D_{min}$ value across all comparisons of 0.044 (**Fig. 2**). We detected no introgression between Plains and Oregon populations, which suggests that Western CPB populations are distinct from Plains and Eastern pest lineages. Izzo *et al*. [1] found shared mitochondrial haploytpes in Oregon and Plains populations, but this could be explained by shared ancestral polymorphism rather than recent gene flow. Second, we tested for introgression with the nine other *Leptinotarsa* species using the same dataset from the SNPhylo analysis, comprising the 100 first scaffolds and 16,519,065 SNPs. The dataset contained one susceptible and one resistant sample for each of the six paired populations, chosen randomly, resulting in N = 2 for Vermont/Maine, Maryland, Wisconsin and Oregon, and N = 1 for each sub-population from New-York/New Jersey and Michigan. In order to analyze a balanced dataset, we limited the "Plains" population to the two samples from Colorado. We also created two Mexican populations, representing the two distinct Mexican clades recovered in our phylogeny: "Mexico City" (containing two samples) and "Mexico South" (containing the sample from Oaxaca and the one from Guerrero; **Table S1**). We used *L. lineolata* as outgroup, as it was recovered as the most basal and distantly-related taxon to the U.S. CPB clade. Again, we found no evidence of introgression among pest populations, as well as no introgression between pest populations and either Mexican or Plains populations (**Fig. S13**). $D_{min}$ reached a maximum of 0.318 between both Mexican populations and remained significant among *Leptinotarsa* species, with the notable exception of the species pairs *L. rubiginosa/L. haldemani*, *L. juncta/L. texana* and *L. peninsularis/L. tumamoca*. The *D* values suggest some introgression into CPB from *L. juncta* and, to a lesser extent, *L. undecemlineata*. However, since introgression was detected between these two species and all ancestral CPB populations (Mexico and the Plains), this pattern does not support the hypothesis of standing genetic variation increasing in the pest lineage as a result of hybridization with another *Leptinotarsa* species. Furthermore, the observed patterns of introgression should be interpreted with caution. Although the *D* statistic is generally robust [2], it can have a high false positive rate under some conditions. Most notably, the *D* statistic is sensitive to incorrect tree topologies, admixture from unsampled populations that might affect the 4-population set [3], and ancestral population structure [4], all of which can increase the *D* statistic.

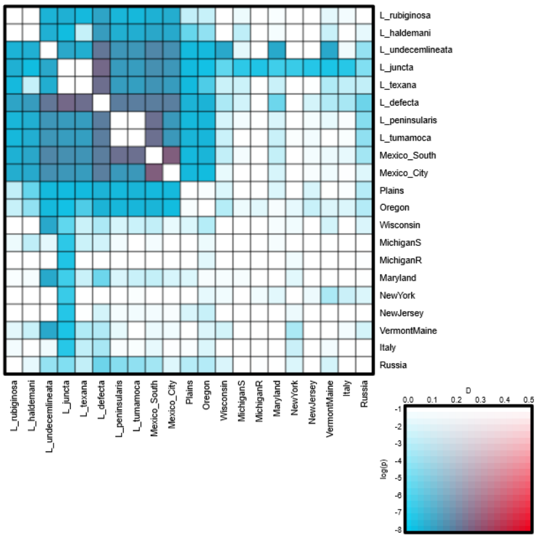

**Figure S13.** Heatmap of *D*-statistics, showing the introgression patterns among CPB populations and other *Leptinotarsa* species. The color of the heatmap cell indicates the most significant $D_{min}$ found with every population pairs: red colors indicate higher *D*-statistic values, and saturated colors indicate higher *p*-values. The dataset contained *L. lineolata* as the outgroup.

### Demographic analyses

Demographic analyses were based on the CPB intergenic SNP dataset, using two 'model-free' skyline plot methods, which have the advantage of exploring larger model space than model-constrained approaches: the stairway plot method [5] and SMC++ [6]. We chose to use a mutation rate of $2.1 \times 10^{-9}$, estimated recently in the non-biting midge [7], as most estimates of nuclear mutation rate in insects fall into the range of $2 \times 10^{-9}$ to $7 \times 10^{-9}$ substitutions per site per generation [8]. We set the generation time to .5/year (*i.e*. 2 generations per year) for all our samples, as this is the norm across most of CPB's geographical range.

**Stairway plots**

The Stairway plot approach relies on the calculation of the expected composite likelihood of a given SNP frequency spectrum (SFS) and is suitable for estimating recent population histories with low coverage genomic data. We analyzed the resistant and susceptible paired populations, both separately and pooled together, and also considered a pooled sample from the Plains (Colorado, Nebraska, Kansas, Missouri, New Mexico and Texas), East (Florida, Tennessee, North Carolina, Virginia, Kentucky and Ohio) and Europe (Italy, Russia). Confidence intervals were almost null when susceptible and resistant populations were analyzed separately (**Fig. S14**). However, when pooled together, the CI remained very narrow and provided more complete reconstructions of population history (<1000 years ago), suggesting an effect of sample size on the analyses.

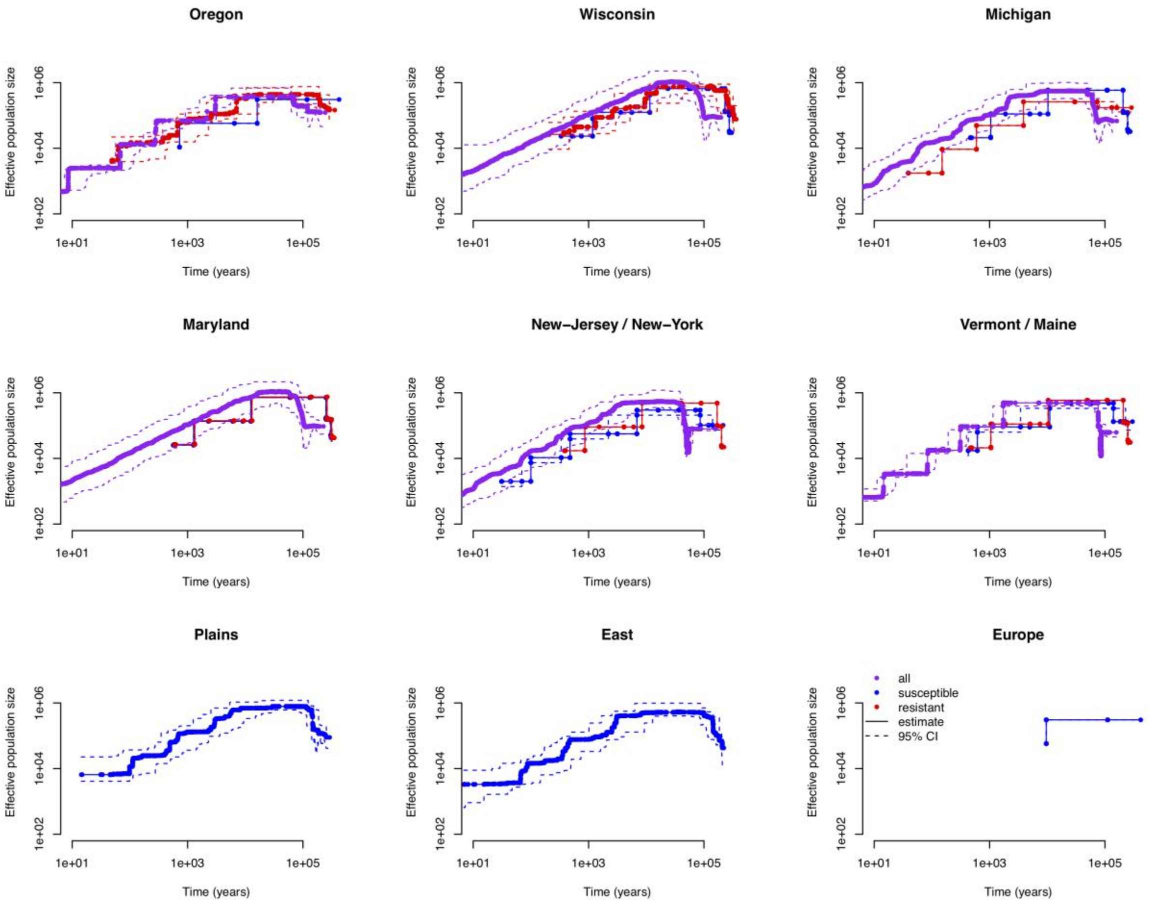

**Figure S14**. Stairway plots depicting demographic histories (median Ne and 2.5–97.5 percentiles) for each population.

When plotted jointly, it is clear that nearly every population displays similar histories (**Fig. S15**). All populations showed a similar ancestral population size of 100,000 individuals around 200,000 years ago. Between 200,000 and 100,000 years, populations from the Midwest, East Coast and Northern U.S experienced a strong decrease in population size, from around 50% in Maryland, Wisconsin and Michigan, up to a 10-fold reduction in New York / New Jersey and Vermont / Maine (event #1 in **Fig. S15**). This was followed by a striking increase in population size for every population, between 150,000 and 70,000 years, with an estimated effective size of up to ~1,000,000 individuals for Wisconsin and Maryland, 790,000 for the Plains, ~490,000 for the Eastern U.S., Michigan and Vermont / Maine, and 375,000 for Oregon (which showed a more gradual increase in Ne compared to other populations; event #2 in **Fig. S15**). Interestingly, the Plains population seem to have expanded first (around 150,000 years ago), quickly followed by the Eastern population. Other population sizes started to increase only 50 (Maryland, Wisconsin), 70 (Michigan, Vermont / Maine) and 100 thousand years later (New-York / New Jersey). After a period of stasis lasting 40,000 to 110,000 years, all population sizes kept decreasing until present down to 500-5,000 individuals. Except for Wisconsin and Maryland, 4 episodes of bottlenecks are identifiable in every population: 5,000-2,000, 700-300, 110-70 and 15-8 years ago (events #3 to #6 in **Fig. S15**).

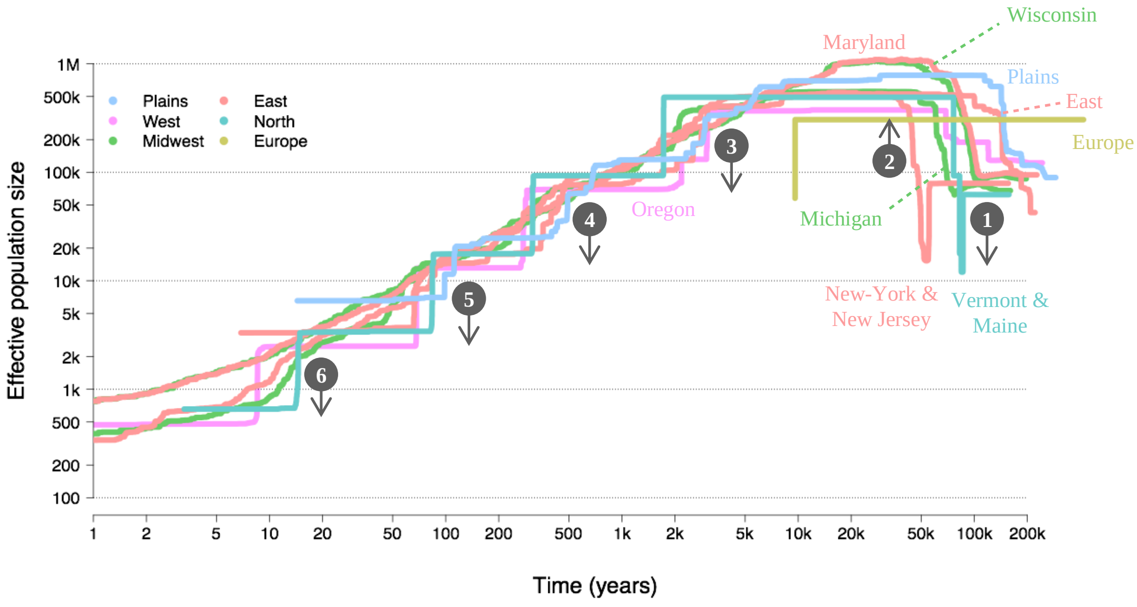

**Figure S15**. Population demographic histories (median Ne only) estimated by Stairway plot analysis. Colors correspond to geographical regions. Numbers and arrows highlight the series of notable demographic events, from 1 to 6 (past to present).

**SMC++**

Demographic reconstruction methods based on SFS assume unlinked segregating sites, which is not true in the case of whole-genome data. SMC++ [6] incorporates estimates of recombination and linkage disequilibrium (LD) in an SFS framework. We analyzed the first 95 scaffolds (>1Mb in length) to ensure we could accurately infer demography using the sequential Markov coalescent, following recommendations by Nadachowska-Brzyska *et al*. [9]. Test results comparing analyses carried on these scaffolds vs. considering all >10 Kb scaffolds (covering 95% of the genome) were comparable (i.e. same population sizes, same demographic events, same time-scale; results not shown). Compared to the Stairway plots, SMC++ reconstructed deeper histories (~500ky vs. 200ky), although more recent histories were not recovered (2-5k years vs. <100 years; **Fig. S16**). Additionally, SMC++ provided a more robust reconstruction of the European samples. SMC++ results were largely consistent with the Stairway plots, showing contracting populations between 300k and 100k years ago, expanding populations between 200k and 70k years, declining population sizes until 10k to 5k years and increasing population sizes more recently. Interestingly, starting population sizes were twice as high and maximum population sizes half as high as those estimated by the Stairway plot method. SMC++ also estimated split times for pairs of populations. Since we hypothesize that CPB colonized the continent from the Plains region, we estimated the split times between the Plains and every other population. When co-estimated with other populations, the Plains' original estimated population size increased from 300k to 900k individuals, reaching a maximum of 1.3M individuals around 100k years (**Fig. S16**). The split with European samples seems inconsistent with other populations, probably due to the very small sample size (N=2). Interestingly, the splits with most populations occurred between 21k and 11k years, during the transition from the late Pleistocene to early Holocene. Historical records, however, place the expansion of pest populations into the east coast in the last two hundred years. The discrepancy in time arises in part because these methods have limited power to reconstruct recent events, but it is also possible that the pest lineages descended from an unsampled population that is genetically distinct from other Plains populations, or that the underlying mutation rate for CPB is unreliable and perhaps higher than the rate of other insects.

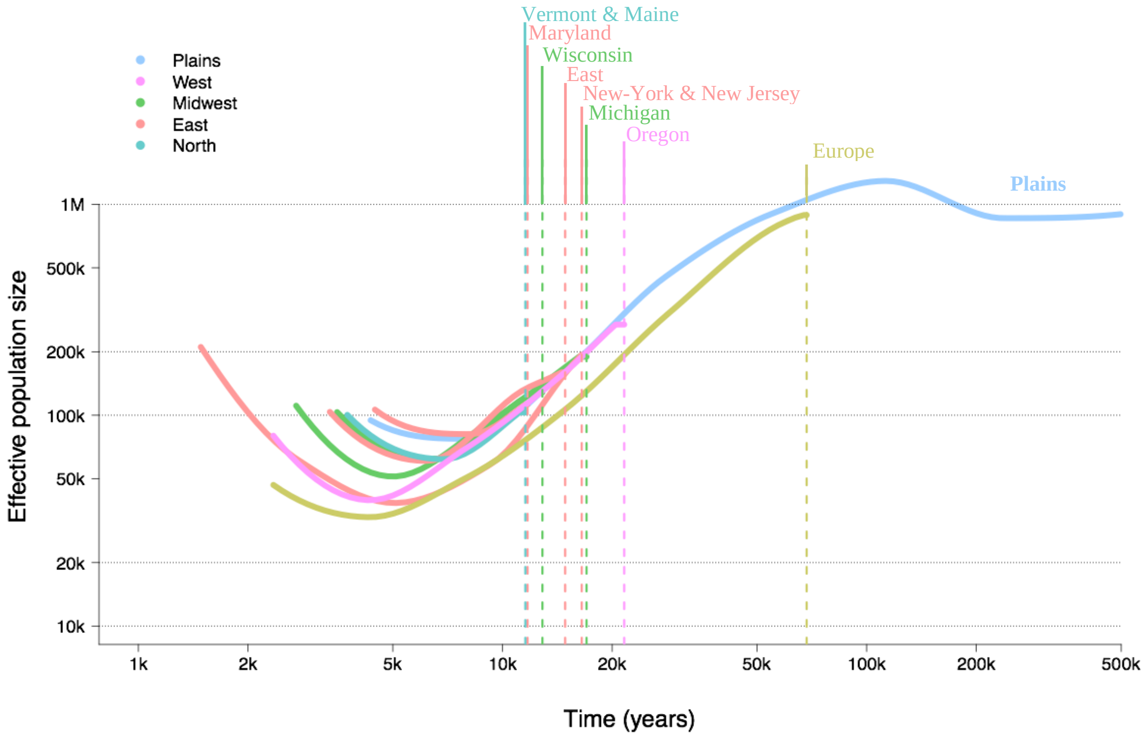

**Figure S16**. Demographic histories (median Ne only) of pest populations and split times from the Plains population, estimated with SMC++. It is assumed that each pest population shared a common demographic history before splits (vertical dotted lines). Colors correspond to geographical regions.

### Selection analyses

We used three different approaches to study genomic signatures of selection: outlier detection with *PCAdapt* [10, 11], genome-environment association with *LFMM* [12] and genome scan with *hapFLK* [13]. These analyses focus on the CPB dataset, with *PCAdapt* and *LFMM* including only U.S. CPB samples (Mexican and European samples have been removed), while *hapFLK* analyses include the European samples.

##### Outlier detection (*PCAdapt*)

To detect outlier SNPs, the false discovery rate (FDR) was controlled at 0.01% (α=0.0001). Filtering at a minor allele frequency (MAF) of 0.05 and choosing a conservative setting of K=10, we identified 1.3% of all SNPs as outliers (i.e. 235,244 out of 17,599,906), of which ~31% belonged to 12,040 known genes (73,495 out of 235,244). As this list suggested a high rate of false positives, we refined our filtering steps using linkage disequilibrium clumping (choosing a window size of 500 SNPs and a squared-correlation coefficient threshold of 0.2). We examined the screeplot of the principal components and, following Cattell's rule, selected K=6 as the optimal clustering level. Finally, we adjusted the dataset by setting the minimum

MAF setting to 0.1. This resulted in 65,814 SNPs at an FDR of 0.01% (α=0.0001), or 0.37% if all SNPs, of which 20,810 (31.6%) occurred within 8,760 known genes (each significant gene had an average and standard deviation of 2.4 ± 2.5 outlier SNPs). Among these genes with signatures of selection, 336 were candidate insecticide resistance genes (**Table S4**). The most notable groups (**Table S5**) included ABC transporters (69 genes), CYP genes (66 genes), and voltage-dependent channel genes (57 genes).

**Table S4**. Summary of outlier SNPs identified by *PCAdapt* and found in known genes.

| | SNPs | Genes |
|---|---|---|
| **In total** | 65814 | - |
| **In known genes** | 20810 | 8760 |
| **Candidate resistance genes** | 862 | 336 |

**Table S5**. Distribution of the candidate genes identified by *PCAdapt* among different mechanisms of insecticide resistance.

| **Mechanisms** | **Categories** | **Genes** |
|---|---|---|
| **Metabolic detoxification** | **Esterases** | 56 |
| | **GSTs** | 5 |
| | **CYPs** | 66 |
| | **ABC transporters** | 69 |
| | **MFS transporters** | 9 |
| **Target-sites** | **Voltage-dependent channels** | 57 |
| | **TRP channel** | 17 |
| | **Acetylcholine receptor (nAChR)** | 16 |
| | **Achetylcholinesterase (AChE)** | 1 |
| **Growth Factors** | **Cuticular proteins** | 40 |

We also examined enriched gene ontology terms generated from *PCAdapt* and found many linked to insecticide resistance and/or stress (**Figure S17**). Among the biological processes (**Fig. S17A**), terms include GO1 oxidation-reduction process, GO2 response to oxidative stress (and multiple nested terms), GO4 chloride and transmembrane transport, GO7 cell redox homeostasis, and protein folding. Among cellular components (**Fig. S17B**), terms include GO2 voltage-gated sodium channel and acetylcholine-gated channel complexes, GO4 integral component of the membrane and membrane, as well as cell junction, presynaptic active zone, and synapse. Among the molecular functions (**Fig. S17C**), terms include GO1 heme binding, GO2 zinc ion binding (including iron ion binding), GO3 extracellular ligand-gated ion channel activity (including voltage-gated sodium channel and acetylcholine receptor activity), GO4 glutathione transferase activity, GO5 ABC transporter activity via the term ATPase activity, coupled to transmembrane movement, GO6 peroxidase activity (including CYP monooxygenase activity), as well as categories such as oxidoreductase activity and structural constituent of the cuticle. Other enriched terms have been implicated in insecticide resistance in some species, such as odorant binding, protein serine/threonine phosphatase activity, and G-protein-coupled receptor (GPCR) signaling pathway [14-16].

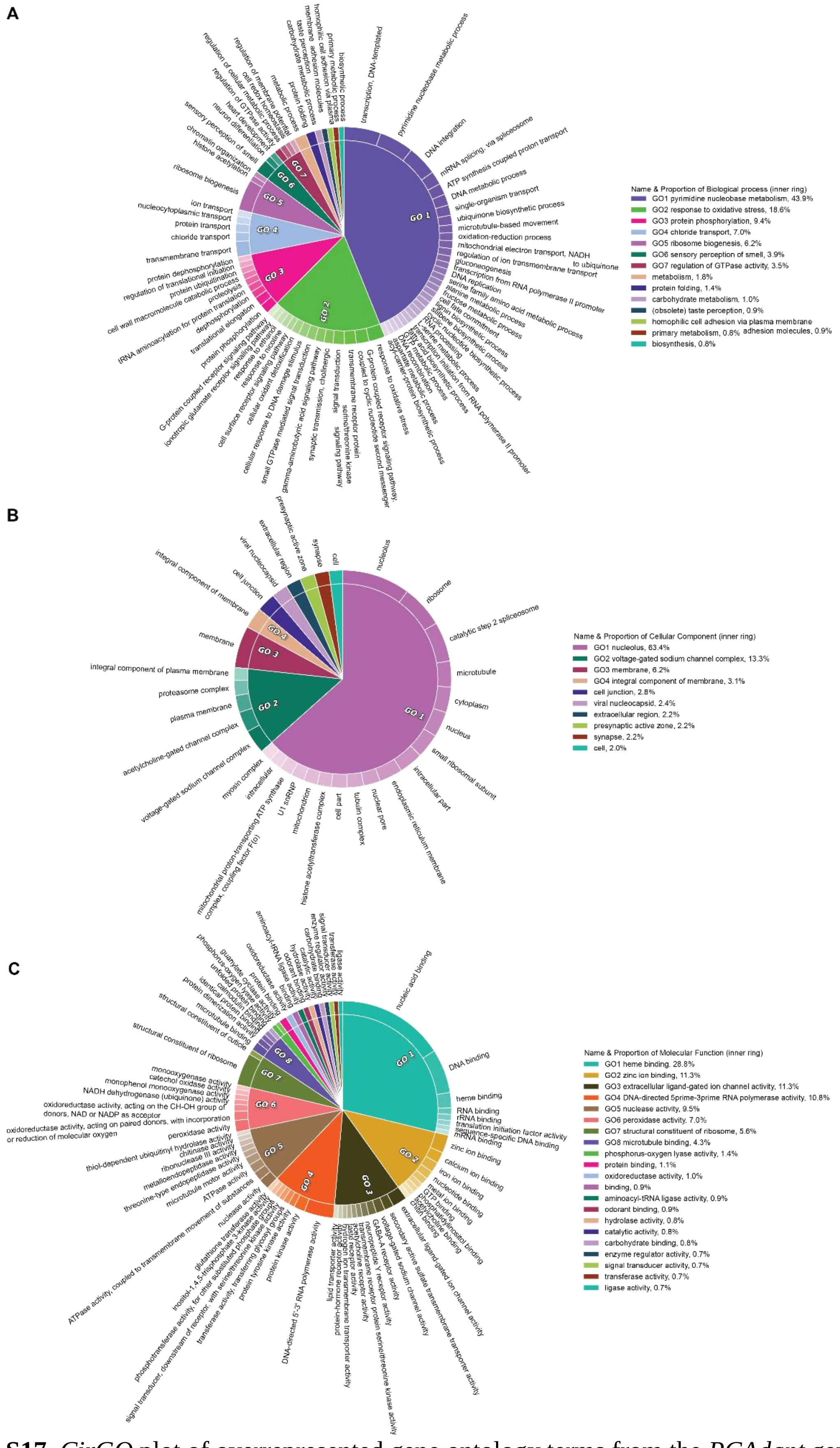

**Figure S17**. *CirGO* plot of overrepresented gene ontology terms from the *PCAdapt* genome scan of selection. Terms are labeled as A) biological processes, B) cellular components, and C) molecular functions. Each box along the outer ring represents a gene ontology term, and these are grouped into 'superclusters' of related terms based on semantic similarity (grouping is indicated by color and highlighted by a GO label, with the relative importance indicated by the size of the pie slice).

**Genome-environment association (*LFMM*)**

A genome-environment association method was used to identify SNPs showing allele frequencies significantly correlated with environmental predictor variables. To account for the population structure observed in our data, we modeled K = 6 latent factors, as suggested by our *PCAdapt* and *sNMF* results. We adjusted *p*-values by an empirically-determined genomic inflation factor, while controlling false discovery rate at 0.01%. We explored five different environmental variables: elevation, precipitation, minimum temperature in the coldest month and potato land cover. We identified 0.02% of the analyzed SNPs as significantly associated to at least one environmental variable (4,098 out of 17,599,906 SNPs; **Table S6**). Out of the 4,098 significant SNPs, 67.6% were associated with the environmental variable *Precipitation*, 15.5% with *Latitude*, 10.7% with *Potato cover*, 2.8% with *Elevation* and 3.7% with *Temperature* (**Fig. S18**), and less than 0.1% (*i.e.* 3 SNPs) with more than one environmental variable. Of all significant SNPs, 28.5% were found in 816 known genes, including 42 candidate insecticide resistance genes (**Table S6**). Genes at target-sites and those involved in metabolic detoxification were most often associated with *Temperature* and *Precipitation*, and to a lesser extent *Latitude* (**Tables S7**).

The gene list from *LFMM* (**Fig. S19**) also shows enrichment of gene ontology terms associated with insecticide resistance and/or stress. Among the biological processes (**Fig. S19A**), terms include GO1 chemical synaptic transmission, GO2 oxidation-reduction process, GO3 intracellular protein transport (transmembrane transport, chloride transport, and ion transport), GO6 proteolysis, defense response, and DNA repair. Among the cellular components **(Fig. S19B**), terms include GO2 synapse and GO6 presynaptic active zone, as well as GO5 membrane and GO8 integral component of the membrane. Among the molecular functions (**Fig. S19C**), terms include GO1 iron ion binding, GO2 carboxylic ester hydrolase activity, GO3 heme binding, GO4 extracellular ligand-gated ion channel activity, GO5 oxidoreductase activity (including monooxygenase activity), GO6 ubiquitin binding. Other possible terms linked to insecticide resistance include the proteasome activator complex [17], pigment binding, and structural constituent of the larval cuticle.

**Table S6**. Summary of SNPs significantly associated with at least one environmental variable by *LFMM*. The number of genes is given in parentheses, next to SNP numbers.

| Mechanisms | | SNPs | Genes | Latitude | Elevation | Precipitation | Temperature | Potato |
|---|---|---|---|---|---|---|---|---|
| Summary | In total | 4098 | - | 633(-) | 115(-) | 2769(-) | 143(-) | 440(-) |
| | In known genes | 1170 | 816 | 195(175) | 53(48) | 735(498) | 52(37) | 136(143) |
| Resistance-related | Metabolic detoxification | 56 | 28 | 9(8) | 1(1) | 18(11) | 25(8) | 3(2) |
| | Cuticle (chitin) | 5 | 3 | 1(1) | 0(0) | 4(2) | 0(0) | 0(0) |
| | Target-sites | 15 | 11 | 3(2) | 1(1) | 9(6) | 1(1) | 1(1) |
| Others | Sensory perception | 11 | 8 | 1(1) | 2(1) | 7(6) | 0(0) | 1(1) |
| | Cellular transport | 135 | 72 | 14(13) | 11(7) | 96(49) | 4(3) | 10(9) |
| | Transposable elements | 75 | 41 | 14(9) | 3(3) | 48(24) | 0(0) | 10(10) |
| | Transcription | 53 | 32 | 9(8) | 2(2) | 29(15) | 0(0) | 13(11) |

| Mechanisms | | SNPs | Genes | Latitude | Elevation | Precipitation | Temperature | Potato |
|---|---|---|---|---|---|---|---|---|
| | **Drug resistance** | 15 | 7 | 4(3) | 1(1) | 9(4) | 1(1) | 0(0) |
| | **Redox** | 4 | 3 | 1(1) | 0(0) | 2(2) | 1(1) | 0(0) |
| | **Immune system** | 1 | 1 | 0(0) | 0(0) | 0(0) | 0(0) | 1(1) |
| | **Nervous system** | 0 | 0 | 0(0) | 0(0) | 0(0) | 0(0) | 0(0) |

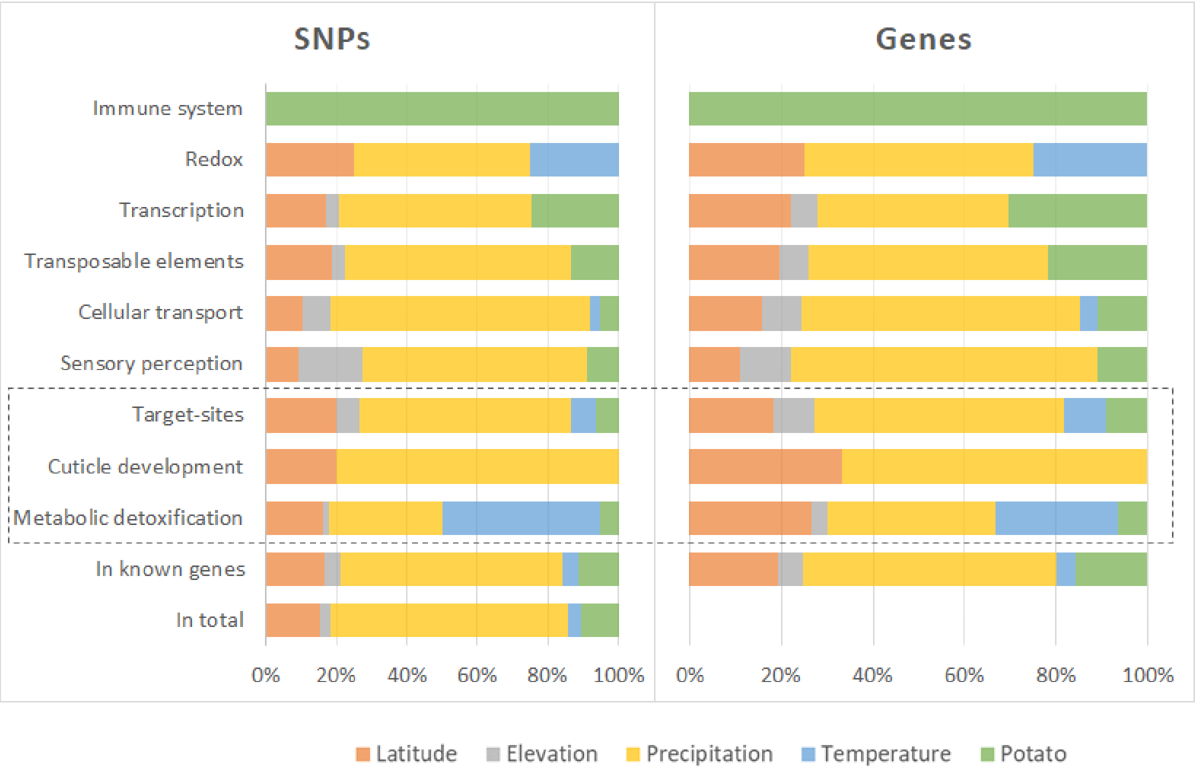

**Figure S18**. Distribution of SNP associations estimated with *LFMM*, for SNPs found in genes. Gene categories include known mechanisms of insecticide resistance (dotted box). Note that this representation is skewed when few SNPs were found to be significant, *i.e.* for redox and immunity genes (cf. **Table S7**).

**Table S7**. Distribution of the SNPs (and genes) significantly associated to at least one environmental variable by *LFMM*, among different categories of the mechanisms of insecticide resistance. The number of genes is given in parentheses, next to SNP numbers.

| Mechanisms | Categories | SNPs | Genes | Latitude | Elevation | Precipitation | Temperature | Potato |
|---|---|---|---|---|---|---|---|---|
| | **ABC transporters** | 19 | 10 | 5(4) | 1(1) | 12(6) | 1(1) | 0(0) |
| | **CYPs** | 25 | 11 | 3(2) | 0(0) | 3(2) | 19(5) | 0(0) |
| **Metabolic detoxification** | **Esterases** | 11 | 5 | 1(2) | 0(0) | 3(3) | 5(2) | 2(1) |
| | **GSTs** | 1 | 1 | 0(0) | 0(0) | 0(0) | 0(0) | 1(1) |
| | **MFS** | 1 | 1 | 0(0) | 1(1) | 0(0) | 0(0) | 0(0) |
| Target-sites | **Voltage-dependent channels** | 11 | 9 | 3(2) | 1(1) | 6(5) | 0(0) | 1(1) |
| | **Known insecticide resistance genes** | 4 | 2 | 0(0) | 0(0) | 3(1) | 1(1) | 0(0) |

| Mechanisms | Categories | SNPs | Genes | Latitude | Elevation | Precipitation | Temperature | Potato |
|---|---|---|---|---|---|---|---|---|
| **Growth Factors** | **Cuticular proteins** | 5 | 3 | 1(1) | 0(0) | 4(2) | 0(0) | 0(0) |

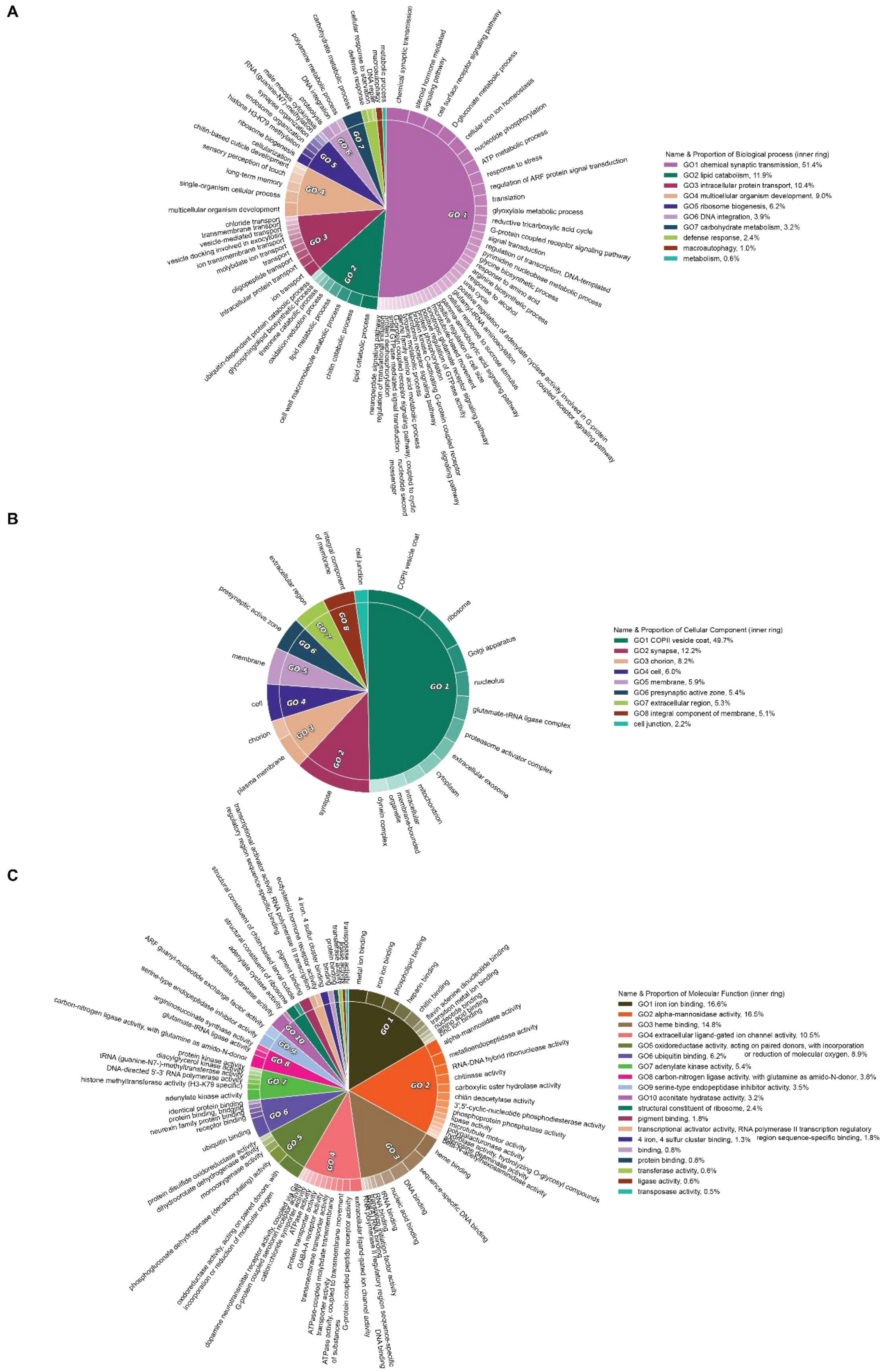

**Figure S19**. *CirGO* plot of overrepresented gene ontology terms from the *LFMM* genome scan of selection. Terms are labeled as A) biological processes, B) cellular components, and C) molecular functions. Each box along the outer ring represents a gene ontology term, and these are grouped into 'superclusters' of related terms based on semantic similarity (grouping is indicated by color and highlighted by a GO label, with the relative importance indicated by the size of the pie slice).

***PCAdapt* / *LFMM* overlap**

A total of 557 genes identified in *LFMM* were also identified as targets of selection in *PCAdapt*, encompassing 68% of the genes highlighted by *LFMM*. Of these, 29 are candidate insecticide resistance genes. We also note that an octopamine receptor (LDEC006841), which was identified as a gene associated with pest behavior [18], is a significant target in both *LFMM* and *PCAdapt*. Overlap between these two genome scan tests (**Fig. S20**) shows enrichment of gene ontology terms associated with insecticide resistance and/or stress. Terms associated with biological processes (**Fig. S27A**) include GO1 chemical synaptic transmission, oxidation-reduction process, gamma-aminobutyric acid signaling pathway, and G-protein coupled receptor signaling pathway; GO2 proteolysis and DNA repair; GO3 chloride transport, transmembrane transport, and ion transport, and GTPase activity. Terms associated with cellular components (**Fig. S27B**) include GO2 integral component of membrane and GO5 membrane, as well as GO7 synapse and GO8 presynaptic active zone. Terms associated with molecular functions (**Fig. S27C**) include pathways such as GO1 heme binding, GO2 ATPase activity coupled to transmembrane movement, GO3 GABA and G-protein coupled receptor activity, GO4 iron ion binding, and GO6 monooxygenase and oxidoreductase activity.

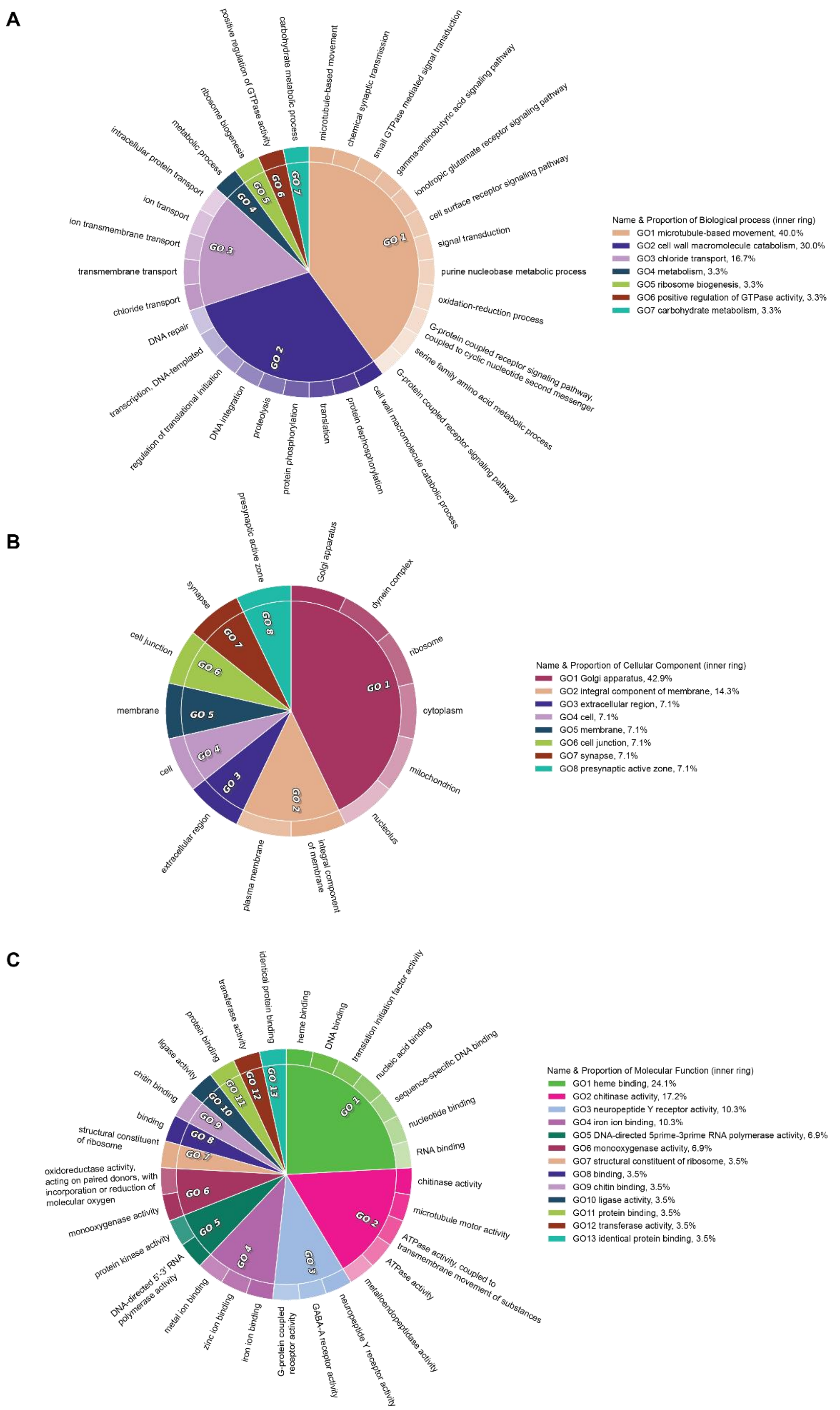

**Figure S20**. *CirGO* plot of overrepresented gene ontology terms shared in both genome scans of selection. Terms are labeled as A) biological processes, B) cellular components, and C) molecular functions. Each box along the outer ring represents a gene ontology term, and these are grouped into 'superclusters' of related terms based on semantic similarity (grouping is indicated by color and highlighted by a GO label, with the relative importance indicated by the size of the pie slice).

**Haplotype-based selection scan (*hapFLK*)**

We analyzed the first (longest) 95 genomic scaffolds of the CPB dataset using the haplotype frequency-based method *hapFLK* [13]. We identified signatures of selection by considering *p-values* < 0.01 (significance threshold at α = 0.01, 0.001 and 0.0001). Only two scaffolds (#45 and #86) did not have any significant candidate SNPs. This resulted in 1,169 selection regions with significant SNPs at α = 0.01, including 140 regions with significant SNPs at α = 0.001 and 24 regions with significant SNPs at α = 0.0001 (**Table S8**). We calculated the average length of regions at each significance threshold to ascertain if the strength of selection was correlated with haplotype block length. One region is ~35 Kb in length without gaps (scaffold 79, position 13917 to 51894), found significant on five branches (all pest populations) spanning the geographic range of CPB, and encodes the genes LDEC004928 and LDEC004929. The former is an uncharacterized membrane-bound protein homologous to *T. castaneum* TC004298 (which is 50% lethal upon knockdown) and known to be immune responsive in *Tenebrio molitor* [19], and the latter is annotated as *tyrosine-protein phosphatase 69d*, a transmembrane protein that regulates synaptic signaling in the brain of *D. melanogaster* [20]. We removed this exceptional region to calculate the average length of the remaining significant regions. Significant regions with larger p-values tended to be shorter in length (**Table S8** and **Fig. S21**).

**Table S8**. Candidate regions detected by *hapFLK*.

| α threshold | Number of regions | Average gap (from previous region) | Average length | Median length | Average number of SNPs |
|---|---|---|---|---|---|
| **0.01** | 1169 | 121386 | 1161 | 734 | 69 |
| **0.001** | 140 | 103737 | 2983 | 2695 | 186 |
| **0.0001** | 24 | 80888 | 3131 | 2736 | 186 |

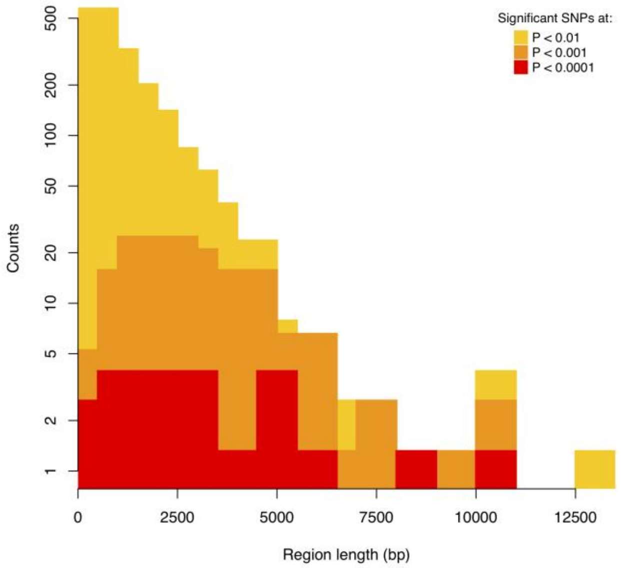

**Figure S21**. Length distribution of selection regions, containing significant SNPs for different α threshold. The *y* axis is on a log scale.

Significant changes in branch lengths at various significance thresholds ($p < 0.01$, $p < 0.001$, $p < 0.0001$) revealed where each candidate selection event was localized in the population tree (**Table S9**). Out of the 1,169 selection regions significant at $\alpha = 0.01$, 48 were not associated to any specific branch, leaving 1,121 regions mapped to the population tree (**Fig. S22**). Non-terminal branches exhibited fewer selection events than terminal branches. Western (both resistant and susceptible), Vermont (susceptible), New Jersey (susceptible), European and Michigan resistant populations displayed the highest number of selection events, with more than 600 regions in these terminal branches (at $\alpha = 0.01$). As results are elevated for New Jersey (an inbred lab population), results may be impacted by differences in the demographic history of each population. However, the overall distribution pattern of selection events did not depend on the significance level of branch support (data not shown) or on SNP significance threshold (α value; **Fig. S22B**), suggesting a slight inflation of p-values due to a violation of model assumptions can't explain these results. Indeed, previous simulation studies have shown that *hapFLK* is not sensitive to demographic bottlenecks and population admixture unless they are relatively strong [13], which does not appear to characterize the CPB samples in this study (**Figs. S3** and **S4**).

Both the number of branches associated with each selection event and the fraction of regions associated with one branch of the population tree only (singular regions) were independent of regions' significance level (SNP support) (**Table S9**). Singular regions tended to be relatively short (1.1 Kb on average) and varied by geographical region (**Figure S22**), most of them being located in the Michigan-resistant/New Jersey/Europe clade, the clade of resistant populations comprising Maine, New-York and Maryland, and the Western clade (Oregon and Idaho). While other singular events were associated with the Wisconsin susceptible and the composite "East" population (comprising all individuals sampled from Eastern US), they vanished under a more stringent level of SNP support and were recovered in *PCAdapt* analyses.

**Table S9**. Total number of regions of selection (defined as containing significant SNPs at $\alpha$ = 0.01, $\alpha$ = 0.001 and $\alpha$ = 0.0001), regions containing resistance-associated genes, average number of branches associated to each region and proportion of singular events, for branch support significant at $p < 0.01$, $p < 0.001$, $p < 0.0001$.

| SNP support (α) | Number of selection regions | Branch support significance level | | |
|---|---|---|---|---|
| | | P < 0.01 | P < 0.001% | P < 0.0001% |
| 0.01 | 1169 [24] | 8.09 (1.1%) | 5.72 (5%) | 4.23 (9.7%) |
| 0.001 | 140 [1] | 8.01 (0%) | 5.71 (3.6%) | 4.3 (7.1%) |
| 0.0001 | 24 [0] | 8.09 (0%) | 6.14 (4.3%) | 4.27 (13%) |

[ ]: number of regions containing resistance-associated genes

( ): percentage of singular events of selection (regions associated with one branch only)

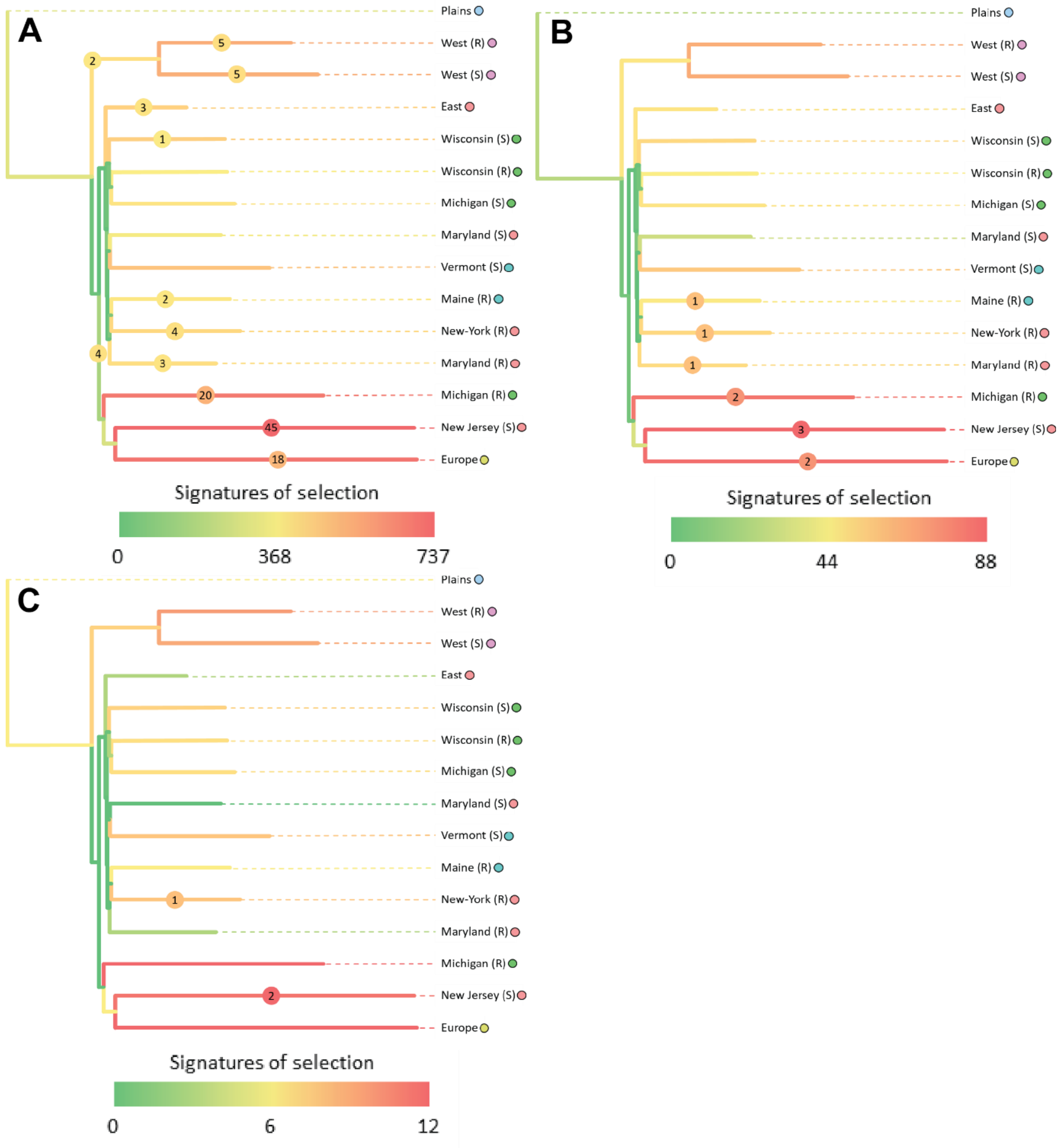

**Figure S22**. Population trees showing the distribution of significant signatures of selection at a branch association threshold $p < 0.0001$. Branch colors represent the number of selection events each branch is associated with. Colored discs on some branches show the number of singular events of selection (associated with one branch only). Colored circles next to populations' names represent their geographical distribution. Panels shows regions of selection defined at various SNP significance thresholds: A) $\alpha = 0.01$, B) $\alpha = 0.001$, C) $\alpha = 0.0001$.

**Function of Putatively Selected SNPs**

Out of 1,169 selection regions from *hapFLK*, 319 were found in 224 genes and 3.8% were singular. Only 24 of the 319 regions contained potential insecticide resistance-associated genes (representing 16 genes in total), including six ABC transporters, two esterases, one olfactory gene (an odorant binding protein), one nicotinic acetylcholine receptor, two genes associated with glutamate pathways and four growth factors (**Table S10**). Interestingly, 19 out of these 24 regions were less than 1 Kb long (**Fig. S23**) and 99.8% of the SNPs in these regions were significant at $\alpha = 0.01$. This might indicate soft selective sweeps or older signatures of selection, as recent hard sweeps create longer haplotype blocks and exhibit higher levels of significance. Three internal branches are notable in their number of selection events: 11 in the West population, five in the shared Europe-New Jersey branch, and four in the Michigan resistant population. The nine remaining internal branches showed fewer than 0.67 selection events on average. Overall, resistant populations did not exhibit more selection events than their susceptible counterparts. Most resistance-associated selection events were pinpointed in 9-to-11 branches, and only two were singular (**Fig. S24**). Many (19 out of 24) were shared between Western and Eastern lineages (who are the most genetically distinct), suggesting that convergent evolution may be prevalent among populations (**Fig. S25**). The Western populations exhibits more resistance-related selection events than other susceptible/resistant clades (**Fig. S22** and **Fig. S25**), with 20 unique selection events vs. 17 for the Wisconsin clade. Additionally, the Western "resistant" branch hosts two unique singular events, involving an esterase gene and a growth inhibitor (**Fig. S25**).

**Table S10**. Candidate regions from *hapFLK* containing potential resistance-associated genes. Growth factors include genes involved chitin synthesis and cuticle formation.

| **Processes** | **Region Number** | **Gene Number** | **SNP Number** |
|---|---|---|---|
| **Total** | 24 | 16 | 1154 |
| **Metabolic detoxification** | 14 | 8 | 713 |
| ABCs | 12 | 6 | 659 |
| CYPs | 0 | 0 | 0 |
| Esterases | 2 | 2 | 54 |
| GSTs | 0 | 0 | 0 |
| **Sensory perception** | 1 | 1 | 168 |
| Olfactory | 1 | 1 | 168 |
| Gustatory/Tactile | 0 | 0 | 0 |
| **Target-sites** | 5 | 3 | 167 |
| Glutamate | 4 | 2 | 113 |
| Sodium | 0 | 0 | 0 |
| Calcium | 0 | 0 | 0 |
| AChE | 0 | 0 | 0 |
| Nicotinic receptor | 1 | 1 | 54 |
| **Growth factors** | 4 | 4 | 106 |

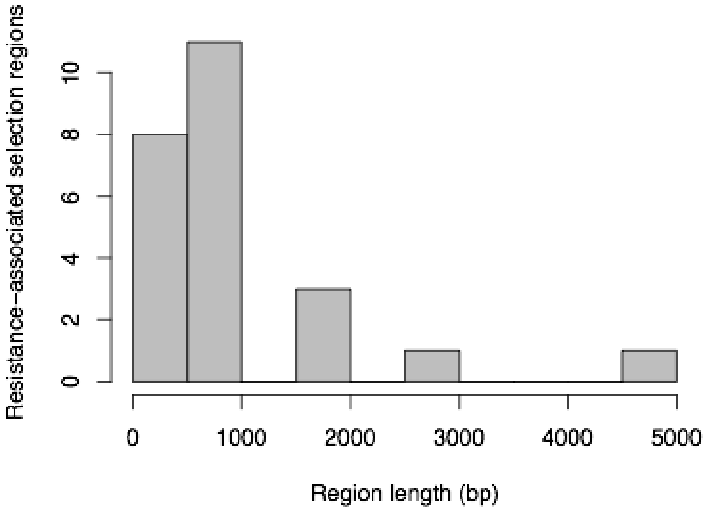

**Figure S23**. Length distribution of selection regions containing potential resistance-associated genes.

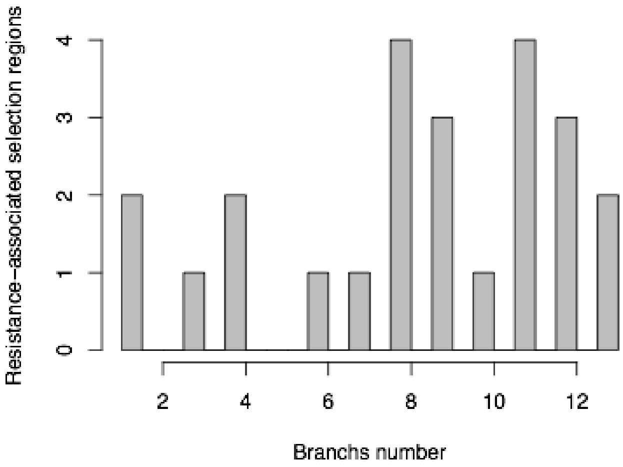

**Figure S24**. Distribution of the number of branches in which resistance-associated selection events were pinpointed.

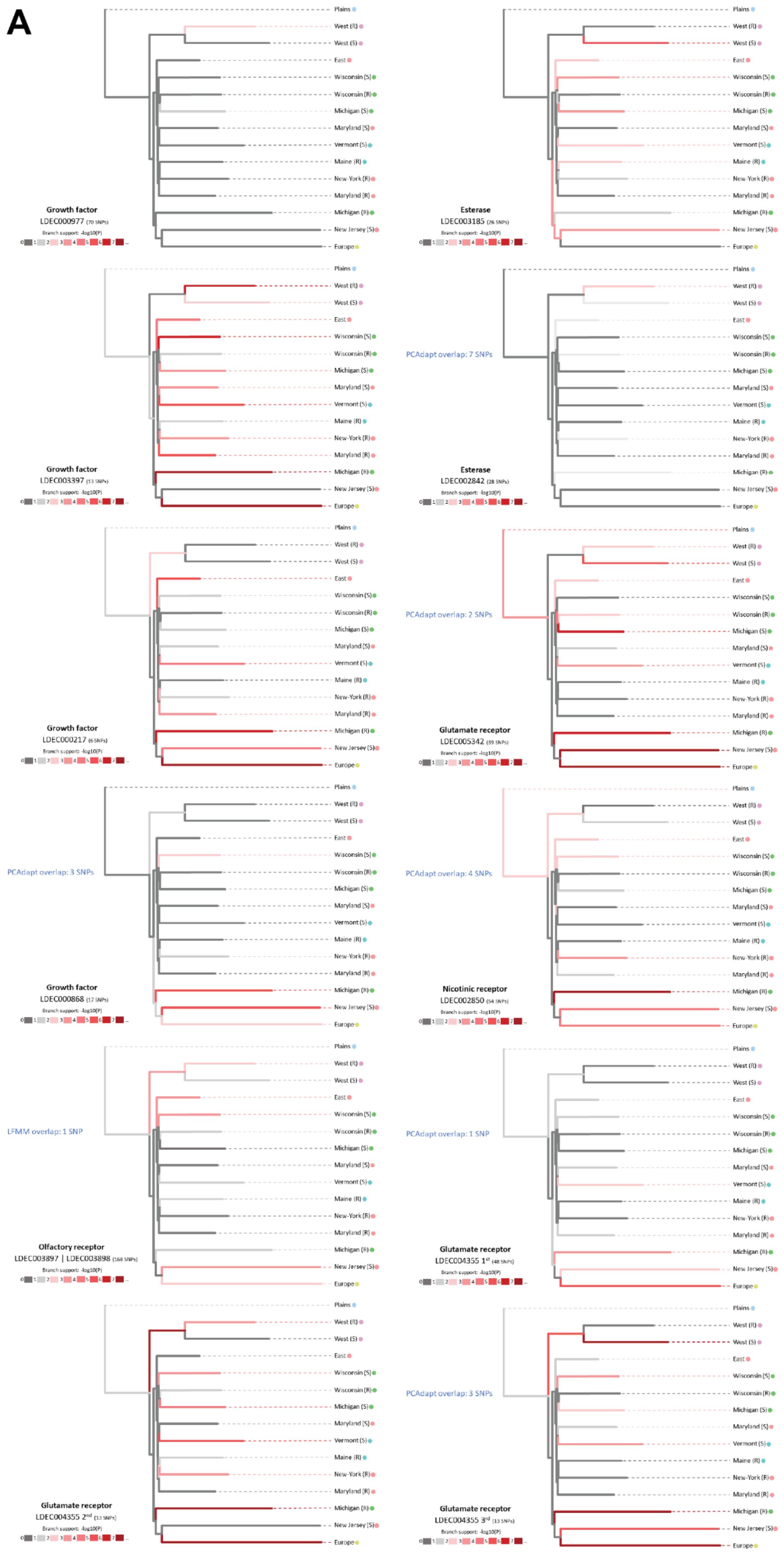
A
Plains
West (R)
West (S)
East
Wisconsin (S)
Wisconsin (R)
Michigan (S)
Maryland (S)
Vermont (S)
Maine (R)
New-York (R)
Maryland (R)
Michigan (R)
New Jersey (S)
Europe
Growth factor
LDEC000977 (70 SNPs)
Esterase
LDEC003185 (26 SNPs)
Growth factor
LDEC003397 (13 SNPs)
PCAdapt overlap: 7 SNPs
Esterase
LDEC002842 (28 SNPs)
Growth factor
LDEC000217 (6 SNPs)
PCAdapt overlap: 2 SNPs
Glutamate receptor
LDEC005342 (39 SNPs)
PCAdapt overlap: 3 SNPs
Growth factor
LDEC000868 (17 SNPs)
PCAdapt overlap: 4 SNPs
Nicotinic receptor
LDEC002850 (54 SNPs)
LFMM overlap: 1 SNP
Olfactory receptor
LDEC003897 | LDEC003898 (168 SNPs)
PCAdapt overlap: 1 SNP
Glutamate receptor
LDEC004355 1st (48 SNPs)
Glutamate receptor
LDEC004355 2nd (13 SNPs)
PCAdapt overlap: 3 SNPs
Glutamate receptor
LDEC004355 3rd (13 SNPs)
Branch support: -log10(P)
0 1 2 3 4 5 6 7 ...

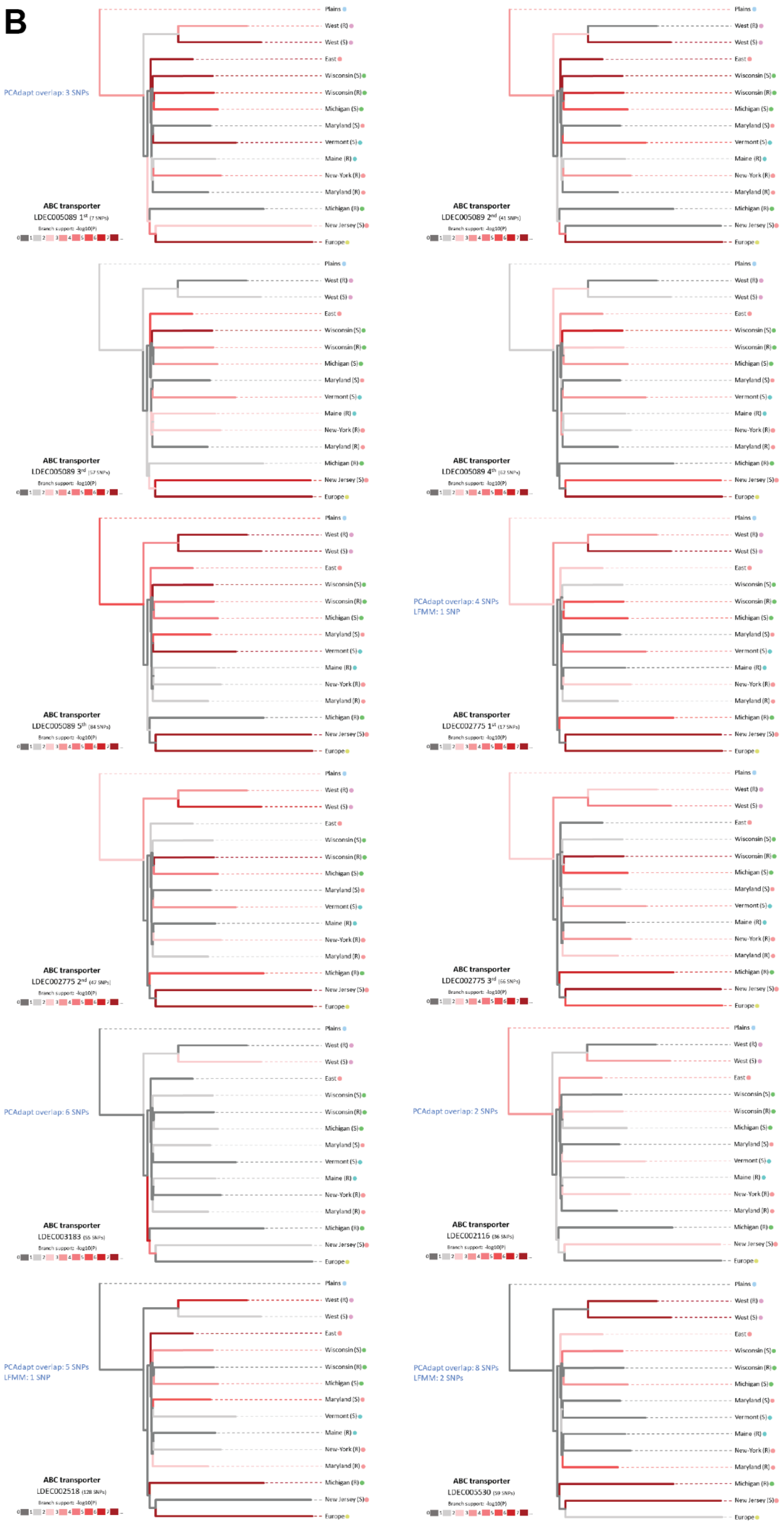
B
PCAdapt overlap: 3 SNPs
ABC transporter
LDEC005089 1st (7 SNPs)
ABC transporter
LDEC005089 2nd (41 SNPs)
ABC transporter
LDEC005089 3rd (57 SNPs)
ABC transporter
LDEC005089 4th (62 SNPs)
ABC transporter
LDEC005089 5th (84 SNPs)
PCAdapt overlap: 4 SNPs
LFMM: 1 SNP
ABC transporter
LDEC002775 1st (17 SNPs)
ABC transporter
LDEC002775 2nd (47 SNPs)
ABC transporter
LDEC002775 3rd (66 SNPs)
PCAdapt overlap: 6 SNPs
ABC transporter
LDEC003183 (55 SNPs)
PCAdapt overlap: 2 SNPs
ABC transporter
LDEC002116 (36 SNPs)
PCAdapt overlap: 5 SNPs
LFMM: 1 SNP
ABC transporter
LDEC002518 (128 SNPs)
PCAdapt overlap: 8 SNPs
LFMM: 2 SNPs
ABC transporter
LDEC005530 (59 SNPs)

**Figure S25.** Individual population trees for the 24 selection regions (containing significant SNPs at α = 0.01) matching resistance-associated genes. A) Panel showing growth factors, nicotinic acetylcholine receptors, glutamate receptors, olfactory receptors and esterases. B) Panel showing ABC transporters. Shades of red show the significance level of the region's association with each branch. Colored dots correspond to geographic regions. Dotted boxes group multiple distinct regions of selection contained in one same gene. The number of SNPs matching the genes' sequences are between brackets.

***PCAdapt* / *hapFLK* overlap**

The *hapFLK–PCAdapt* overlap (**Fig. S26**), based on a threshold of α=0.01 in *hapFLK*, consisted of 150 genes (68.5% of the significant genes). A total of 11 insecticide resistance-associated genes from *hapFLK* contained outliers from the *PCAdapt* analysis. We include one odorant binding protein (LDEC003898) among the set of 11 genes, as this could possibly be linked to insecticide resistance [16], presumably through sensory perception of leaf surfaces with insecticidal toxins.

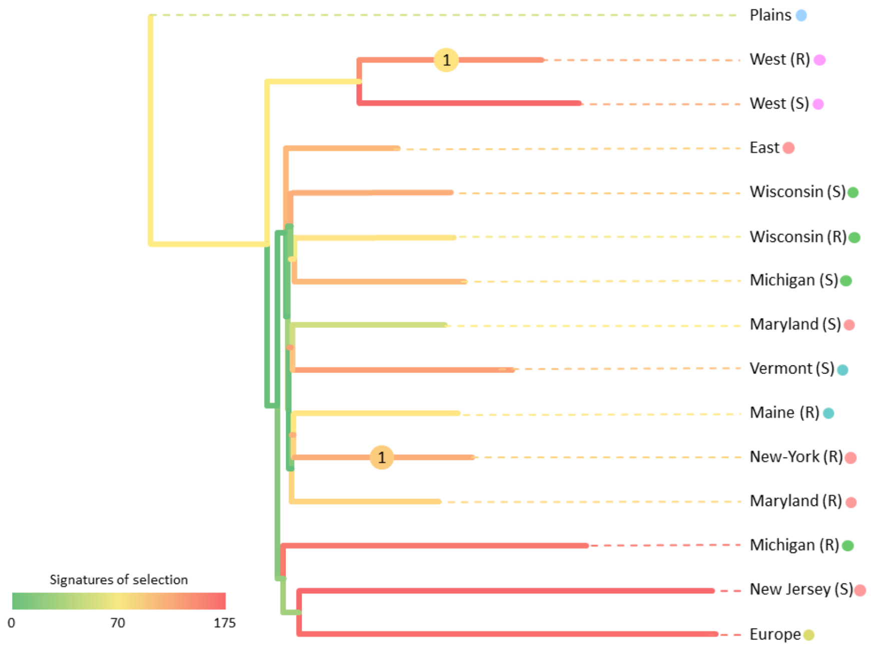

**Figure S26**. Population trees showing the distribution of significant signatures of selection in *hapFLK* (α = 0.01) that are also found as outliers in the *PCAdapt* analysis. Branch colors represent the number of selection events each branch is associated with. Colored discs on some branches show the number of singular events of selection (associated with one branch only). Colored circles next to populations' names represent their geographical distribution.

### Gene expression analysis

#### Samples and experimental treatments

In order to test for rapid regulatory evolution, we compared gene expression data from RNA sequencing (RNAseq) experiments across the geographical range of CPB, including original data from the Plains region (a Colorado population) and previously published pest CPB population samples (see **Table S11**).

**Table S11**. RNA sequencing samples used for differential gene expression analysis, grouped by study.

<table>
<tr><th>Population</th><th>N</th><th>Imidacloprid Resistance</th><th>Induction</th><th>Tissue</th><th>NCBI SRA Accession#</th><th>Reference</th></tr>
<tr><td>Holly, Colorado (CO)</td><td>6</td><td>Susceptible</td><td>No</td><td>Adult body</td><td>SRR12121893-SRR12121888</td><td>This study</td></tr>
<tr><td rowspan="2">Hermiston, OR (OR)</td><td>3*</td><td>Susceptible</td><td>No</td><td>Larvae body</td><td rowspan="4">SRR13510812-SRR13510823</td><td rowspan="4">[21]</td></tr>
<tr><td>3*</td><td>Susceptible</td><td>Yes</td><td>Larvae body</td></tr>
<tr><td rowspan="2">Riverhead (Long Island), NY (NY)</td><td>3*</td><td>Resistant</td><td>No</td><td>Larvae body</td></tr>
<tr><td>3*</td><td>Resistant</td><td>Yes</td><td>Larvae body</td></tr>
<tr><td>Long Island, New York (NY)</td><td>3</td><td>Resistant</td><td>No</td><td>Adult body</td><td>SRR6830862-SRR6830864</td><td>[22]</td></tr>
<tr><td>Arlington, Columbia County, Wisconsin (WI)</td><td>3*</td><td>Susceptible</td><td>No</td><td>Adult body</td><td rowspan="3">SRR2556962, SRR2600374, SRR2600376, SRR2600438, SRR2600681, SRR2600974, SRR2600998-SRR2601000</td><td rowspan="3">[23]</td></tr>
<tr><td>Hancock, Waushara County, Wisconsin (WI)</td><td>3*</td><td>Resistant 1st generation</td><td>No</td><td>Adult body</td></tr>
<tr><td>Hancock, Waushara County, Wisconsin (WI)</td><td>3*</td><td>Resistant 2nd generation</td><td>No</td><td>Adult body</td></tr>
<tr><td rowspan="2">Arlington, Columbia County, Wisconsin (WI)</td><td>4</td><td>Susceptible</td><td>No</td><td>Larvae body</td><td rowspan="2">SRR7127649-SRR7127664</td><td rowspan="2">[24]</td></tr>
<tr><td>4</td><td>Susceptible</td><td>Yes</td><td>Larvae body</td></tr>
<tr><td>Lab Selected for 51 generations: Long Island, New York (NY)</td><td>3*</td><td>Resistant</td><td>No</td><td>Adult body</td><td>SRR3999901-SRR3999903</td><td rowspan="2">[25]</td></tr>
<tr><td>London, Ontario, Canada (CAN)</td><td>3*</td><td>Susceptible</td><td>No</td><td>Adult body</td><td>SRR4069274-SRR4069276</td></tr>
<tr><td>French Biolabs-USDA-New Jersey Department of Agriculture colony, West Trenton, New Jersey (NJ)</td><td>6</td><td>Susceptible</td><td>No</td><td>Larvae</td><td>SRR1820770, SRR1820776, SRR1820785, SRR1820843, SRR1820867, SRR1820877</td><td rowspan="3">[26]</td></tr>
<tr><td>French Biolabs-USDA-New Jersey Department of Agriculture colony, West Trenton, New Jersey (NJ)</td><td>3</td><td>Susceptible</td><td>No</td><td>Adult midgut</td><td rowspan="2">SRR6238881-SRR6238883, SRR6238885, SRR6238887, SRR6238889</td></tr>
<tr><td>Lab Selected for 52 generations: Montcalm County, Michigan (MI)</td><td>3</td><td>Resistant</td><td>No</td><td>Adult midgut</td></tr>
</table>

*Samples with an asterisk are the number of sampled pools, where each pool comprises RNA extracted from multiple individuals simultaneously.

#### Differential expression results

We conducted a series of preliminary tests to identify the effects of different experimental conditions on gene expression. We first tested whether samples of post-diapause (1st generation, overwintered) adult beetles from a resistant population in Hancock, Wisconsin samples showed differential gene expression relative to non-diapausing (2nd generation, summer) adults. No genes were significantly differentially expressed. We then compared whether imidacloprid exposure caused an induced response relative to control samples in lab-reared larvae from Wisconsin, Oregon and Long Island. No genes were significantly differentially expressed in the populations from Oregon and Long Island, but the Wisconsin population showed 77 genes upregulated in the induced treatment and one gene downregulated. Two candidate insecticide resistance genes were significantly induced: a probable mitochondrial cytochrome p450 (LDEC021046) and a cuticle protein (LDEC015588). Third, we compared whether comparisons of life stages (larvae versus adults) resulted in differential expression by testing susceptible, lab-reared larvae against susceptible field-sampled adults from Arlington, Wisconsin. The lab-reared larvae included a control and an imidacloprid-induced treatment, which were compared separately to the field-sampled adults. In both cases, field-sampled adults showed upregulation of a large number of genes (624 in the larvae lab control vs. adult comparison, 24 in the larvae lab induction vs. adult comparison). The larvae lab control vs. adult included six candidate insecticide resistance genes: LDEC008011 (glutamate receptor NMDA 1), LDEC010662 (probable cytochrome p450 4s3), LDEC0023573 (cytochrome p450), LDEC016273 (probably cytochrome p450 305a1), and LDEC015588 (cuticle protein). The larvae lab induction vs. adult included one candidate insecticide resistance gene, the same glutamate receptor NMDA 1 (LDEC008011), which regulates ion-channels in nerve cells.

Based on these comparisons, we determined that regional population differences could be compared for adults from field collected populations irrespective of generation sampled, but lab reared larvae needed to be compared separately. For the larval comparison, we removed samples representing an induction treatment. Overall expression patterns showed considerable gene expression variance among populations. Samples group primarily by geography in the adults (**Fig. S27**), rather than imidacloprid resistance status. The larvae showed even stronger geographical differentiation (**Fig. S28**). In the adult comparison, 85 genes were differentially expressed across populations, whereas 1,312 genes were differentially expressed in the larval comparison. To refine these comparisons, we focused on significant expression differences at candidate insecticide resistance genes and transcription factors.

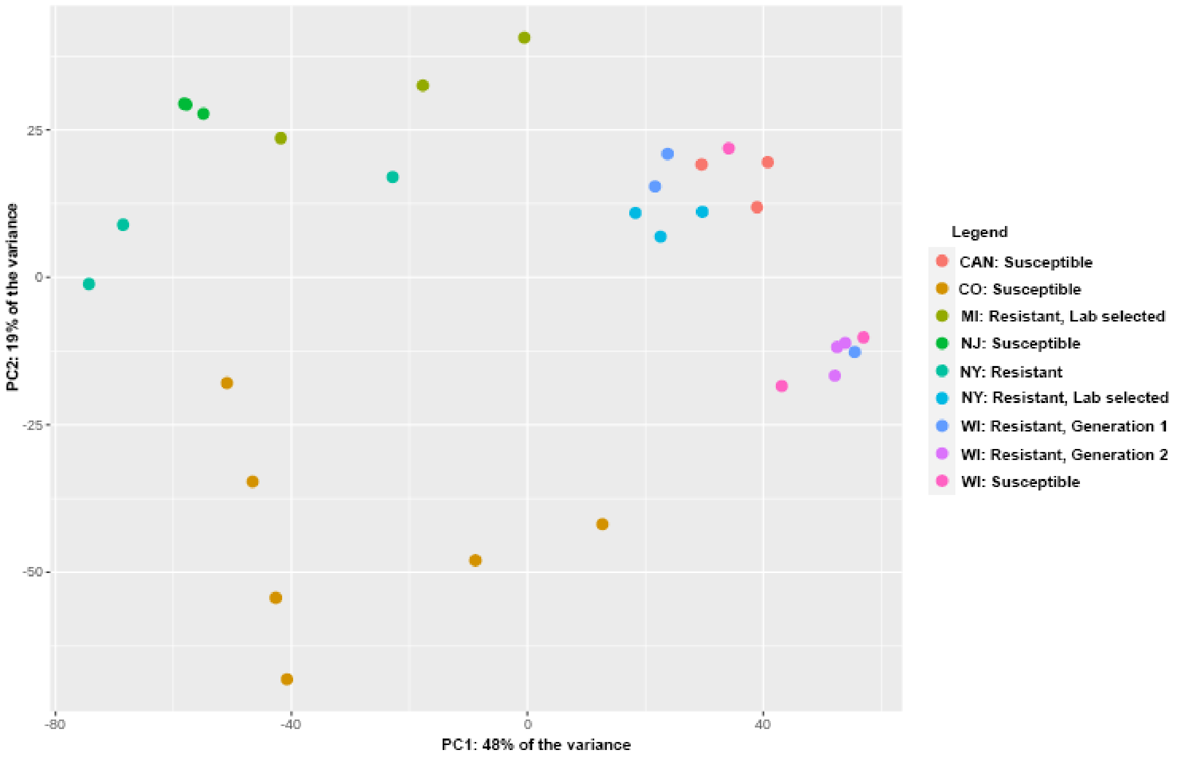

**Figure S27**. A principal component analysis of gene expression among adult CPB samples. Geographical populations show grouping by regional location, rather than imidacloprid resistance status.

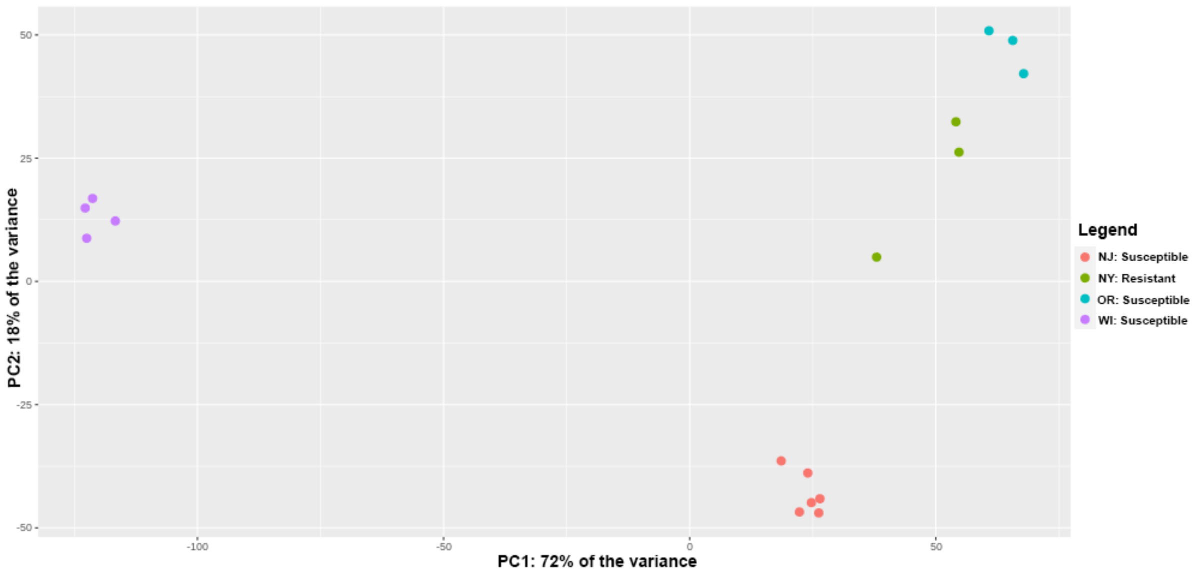

**Figure S28**. A principal component analysis of gene expression among lab-reared larval CPB samples. Geographical populations show grouping by regional location, rather than imidacloprid resistance status.

In the adult comparison, seven candidate insecticide resistance genes were found among the significantly differentially expressed genes (**Fig. S29**). This includes five cytochrome P450s (LDEC023780, LDEC017468, LDEC021334, LDEC021333, LDEC022309), one esterase (LDEC019310), and one ABC transporter (LDEC004154). Notably, these genes showed high expression levels in Wisconsin, downregulation in Canada, and more varied expression across other populations, suggesting selection alters constitutive expression levels of insecticide resistance genes uniquely across populations. No transcription factors were significantly differentially expressed in comparisons of the adult samples. A gene set enrichment analysis of the differentially expressed gene list in among-population comparisons of adults (**Figure S30**) showed enrichment of adult biological processes (tracheal system, male meiosis cytokinesis, and multicellular organism reproduction) and molecular functions related to plant digestion (serine and cysteine type peptidase activity). More notably, there was enrichment of terms associated with regulatory changes to gene networks underlying insecticide detoxification and/or stress, such as heme and ion iron binding, oxidoreductase activity and dioxygenase activity, proteolysis, transport, defense response, and integral component of the membrane.

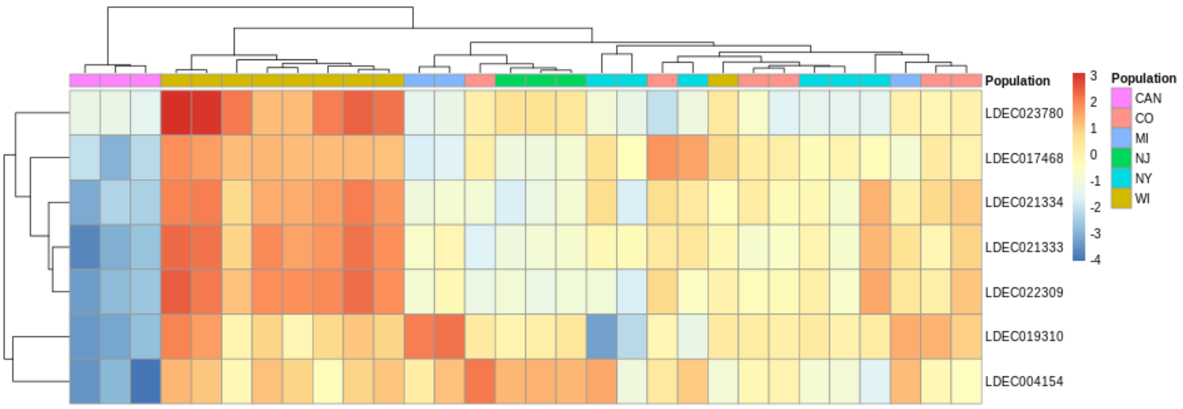

**Figure S29**. Heatmap of seven candidate insecticide resistance genes that were significantly differentially expressed in adult samples of CPB populations. Colors of expression levels correspond to log-fold change. Genes include: five cytochrome P450s (LDEC023780, LDEC017468, LDEC021334, LDEC021333, LDEC022309), one esterase (LDEC019310), and one ABC transporter (LDEC004154).

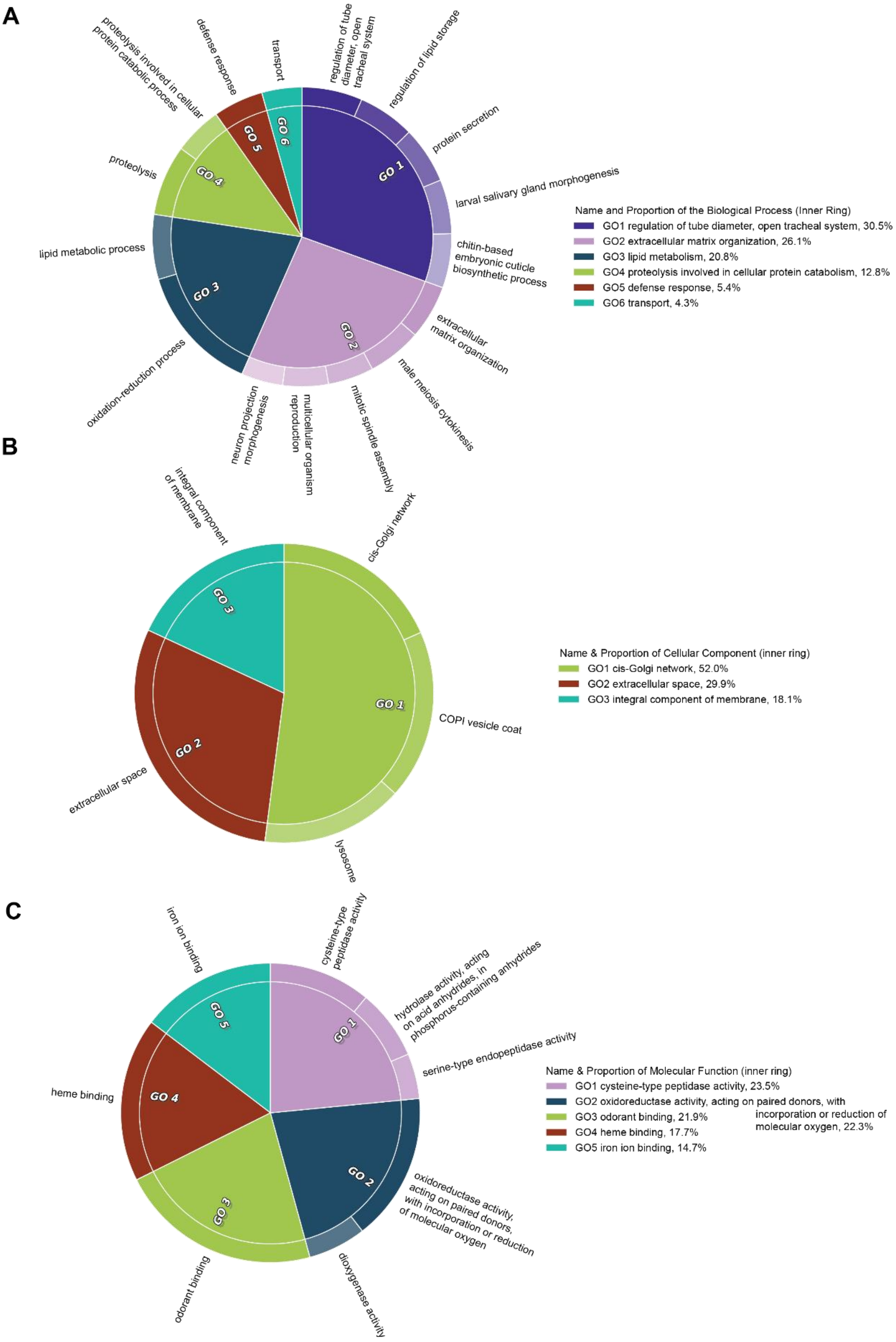

**Figure S30**. *CirGO* plot of overrepresented gene ontology terms from the adult among-population differential gene expression analysis. Terms are labeled as A) biological processes, B) cellular components, and C) molecular functions. Each box along the outer ring represents a gene ontology term, and these are grouped into 'superclusters' of related terms based on semantic similarity (grouping is indicated by color and highlighted by a GO label, with the relative importance indicated by the size of the pie slice).

The larval comparison showed similar divergence in constitutive expression levels of insecticide resistance genes across populations. Here, 84 candidate insecticide resistance genes were found among the significantly differentially expressed genes (**Table S12**). This included eight nerve cell receptors, nine ABC transporters, 15 cytochrome P450s, eight esterases, four glutathione S-transferases, four carboxylesterases, and 36 cuticle proteins. A gene set enrichment analysis of the differentially expressed gene list in among-population comparisons of larvae (**Fig. S32**) showed enrichment of larvae-specific biological processes (multicellular organism development, regulation of development, regulation of cell proliferation, and chitin-based cuticle development), cellular component (collagen trimer), and molecular functions (structural constituent of cuticle and growth factor activity). More interestingly, there was enrichment of terms associated with regulatory changes to gene networks underlying insecticide detoxification or stress, such as glutathione transferase activity, hexachlorocyclohexane metabolism, oxidoreductase and monooxygenase activity, gap junction channel activity, proteolysis, substrate-specific transmembrane transporter activity, heme and iron ion binding, innate immune response, and integral component of the membrane.

Additionally, 26 transcription factors were significantly differentially expressed in comparisons of the four larval populations (**Fig. S33**, **Table S13**). However, these appear to function in core developmental processes or broad transcriptional regulatory processes, and therefore likely indicate variation in developmental stages of the focal larval populations.

**Table S12.** Functional annotation of 84 candidate insecticide resistance genes that were significantly differentially expressed in larval samples of CPB populations.

| Gene ID | Gene Annotation | Gene ID | Gene Annotation |
|---|---|---|---|
| LDEC004025 | acetylcholine receptor subunit alpha-like 1 | LDEC010804 | flexible cuticle protein 12-like |
| LDEC009862 | glutamate receptor 2-like isoform x4 | LDEC006245 | larval cuticle LCP-30 |
| LDEC022809 | glutamate receptor kainate 1-like protein | LDEC001814 | larval cuticle protein 8-like |
| LDEC008011 | Glutamate receptor NMDA 1 | LDEC010803 | larval cuticle protein 8-like |
| LDEC015955 | voltage-dependent calcium channel type d subunit alpha-1-like protein | LDEC015102 | larval cuticle protein a2b |
| LDEC004594 | voltage-dependent t-type calcium channel subunit alpha-1g-like isoform x1 | LDEC011031 | larval cuticle protein a2b-like |
| LDEC024133 | sodium channel protein 60e-like protein | LDEC011036 | larval cuticle protein a2b-like |
| LDEC005495 | transient receptor potential channel pyrexia isoform x2 | LDEC011038 | larval cuticle protein a2b-like |
| LDEC017669 | atp-binding cassette sub-family g member 8 | LDEC001829 | larval cuticle protein lcp-17 |
| LDEC021280 | multidrug resistance-associated | LDEC000824 | larval cuticle protein lcp-30 |
| LDEC021941 | multidrug resistance-associated 1 isoform X1 | LDEC000825 | larval cuticle protein lcp-30 |
| LDEC004154 | multidrug resistance-associated protein 4 | LDEC012201 | larval pupal cuticle protein h1c |
| LDEC004155 | multidrug resistance-associated protein 4 | LDEC012202 | larval pupal cuticle protein h1c |
| LDEC021955 | probable multidrug resistance-associated lethal(2)03659 | LDEC012203 | larval pupal cuticle protein h1c |
| LDEC002119 | probable multidrug resistance-associated protein lethal 03659 | LDEC012204 | larval pupal cuticle protein h1c |
| LDEC002518 | probable multidrug resistance-associated protein lethal 03659 | LDEC012205 | larval pupal cuticle protein h1c |
| LDEC014173 | probable multidrug resistance-associated protein lethal 03659 | LDEC024732 | larval pupal cuticle protein h1c-like |
| LDEC024466 | cytochrome P450 | LDEC004412 | tweedle motif cuticular protein 2 |
| LDEC015420 | cytochrome p450 | LDEC019310 | esterase |
| LDEC019189 | cytochrome P450 [Tribolium castaneum] | LDEC023088 | esterase |
| LDEC006084 | cytochrome P450 [Tribolium castaneum] | LDEC015319 | esterase |
| LDEC004501 | cytochrome p450 6a2 | LDEC017039 | esterase |
| LDEC005460 | cytochrome p450 6bq10 | LDEC011787 | esterase-6-like protein |
| LDEC004500 | cytochrome p450 6bq8 | LDEC011788 | esterase-6-like protein |
| LDEC001002 | cytochrome P450 6k1-like | LDEC017040 | esterase-6-like protein |
| LDEC007963 | cytochrome P450 6k1-like | LDEC017041 | esterase-6-like protein |
| LDEC008391 | cytochrome p450 6k1-like | LDEC015216 | glutathione S-transferase 1 isoform x2 |
| LDEC011604 | cytochrome p450 6k1-like | LDEC022984 | glutathione S-transferase 1-1 |
| LDEC010650 | cytochrome p450 mitochondrial | LDEC001117 | glutathione S-transferase 1-like |
| LDEC010280 | probable cytochrome p450 4aa1 | LDEC001119 | glutathione S-transferase epsilon |
| LDEC010662 | probable cytochrome p450 4s3 | LDEC021156 | venom carboxylesterase-6 |
| LDEC004818 | probable cytochrome p450 mitochondrial | LDEC021157 | venom carboxylesterase-6-like |
| LDEC001016 | cuticle 8-like | LDEC022654 | venom carboxylesterase-6-like |
| LDEC001803 | cuticle protein | LDEC009957 | venom carboxylesterase-6-like |
| LDEC011039 | cuticle protein | LDEC001813 | flexible cuticle protein 12 |
| LDEC011040 | cuticle protein | LDEC010805 | flexible cuticle protein 12 |
| LDEC011041 | cuticle protein | LDEC001804 | flexible cuticle protein 12-like |
| LDEC011042 | cuticle protein | LDEC010804 | flexible cuticle protein 12-like |
| LDEC011043 | cuticle protein | LDEC006245 | larval cuticle LCP-30 |
| LDEC011468 | cuticlin-1 | LDEC001814 | larval cuticle protein 8-like |
| LDEC016622 | cuticlin-1-like protein | LDEC010803 | larval cuticle protein 8-like |
| LDEC008714 | cuticular protein | LDEC015102 | larval cuticle protein a2b |
| LDEC008715 | cuticular protein | LDEC011031 | larval cuticle protein a2b-like |
| LDEC012195 | cuticular protein | LDEC011036 | larval cuticle protein a2b-like |
| LDEC004413 | cuticular protein 4 | LDEC011038 | larval cuticle protein a2b-like |
| LDEC004414 | cuticular protein 4 | LDEC001829 | larval cuticle protein lcp-17 |
| LDEC005020 | cuticular protein analogous to peritrophins 1-b | LDEC000824 | larval cuticle protein lcp-30 |
| LDEC001813 | flexible cuticle protein 12 | LDEC000825 | larval cuticle protein lcp-30 |
| LDEC010805 | flexible cuticle protein 12 | LDEC012201 | larval pupal cuticle protein h1c |
| LDEC001804 | flexible cuticle protein 12-like | | |

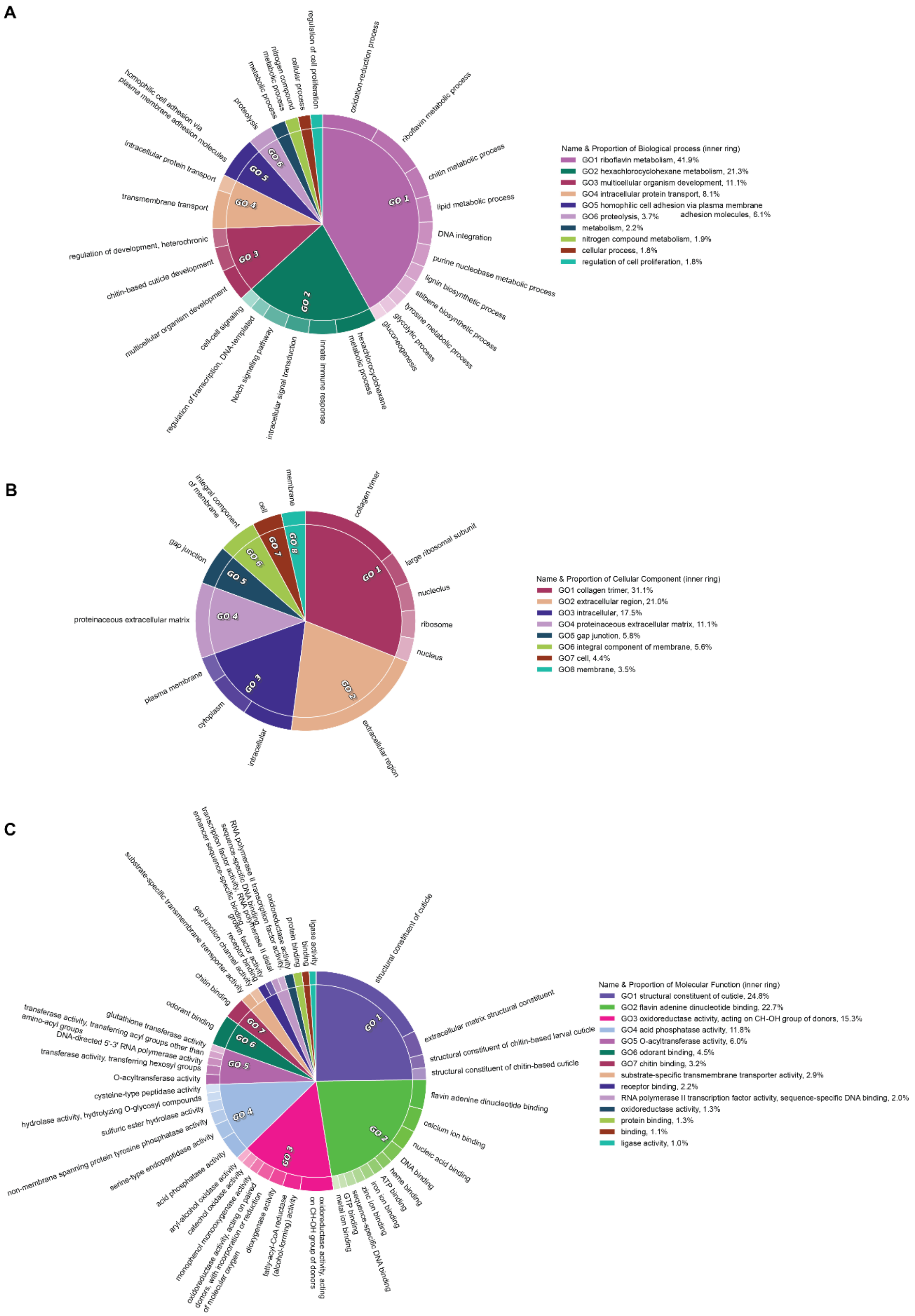

**Figure S31**. *CirGO* plot of overrepresented gene ontology terms from the larval among-population differential gene expression analysis. Terms are labeled as A) biological processes, B) cellular components, and C) molecular functions. Each box along the outer ring represents a gene ontology term, and these are grouped into 'superclusters' of related terms based on semantic similarity (grouping is indicated by color and highlighted by a GO label, with the relative importance indicated by the size of the pie slice).

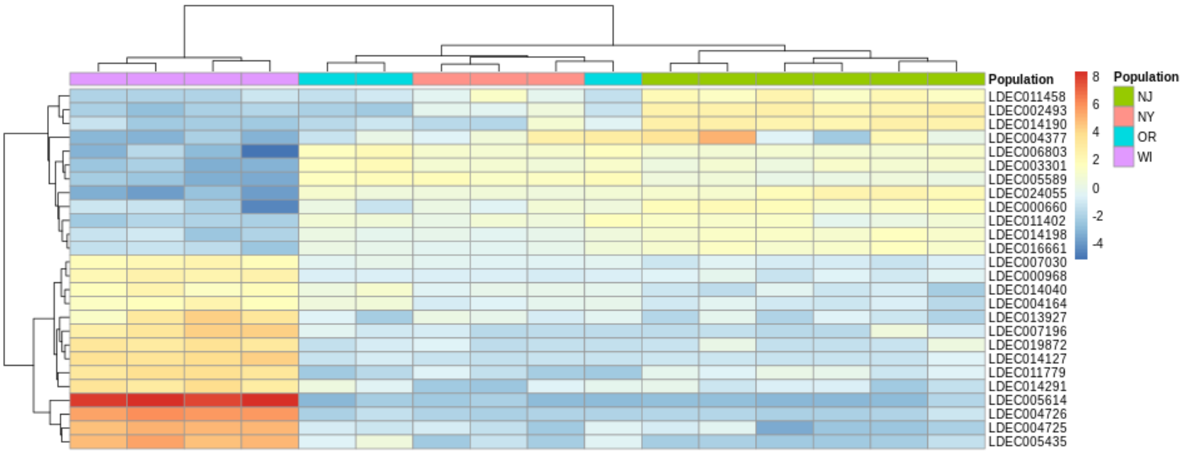

**Figure S32**. Heatmap of 26 transcription factors that were significantly differentially expressed in larval samples of CPB populations. Colors of expression levels correspond to log-fold change. See **Table S13** for the functional annotation of these genes.

**Table S13.** Functional annotation of 26 transcription factors that were significantly differentially expressed in larval samples of CPB populations.

| Gene ID | Gene Annotation | Gene ID | Gene Annotation |
|---|---|---|---|
| LDEC019872 | doublesex | LDEC014198 | homeotic protein antennapedia-like |
| LDEC006803 | transcription factor 21 | LDEC014291 | transcription factor |
| LDEC007030 | transcription initiation factor TFIID subunit 7 | LDEC000660 | homeobox protein |
| LDEC007196 | embryonic polarity dorsal isoform X1 | LDEC004164 | shootin-1 |
| LDEC002493 | knirps-related | LDEC004377 | probable basic-leucine zipper transcription factor i |
| LDEC011402 | transcription factor glial cells missing | LDEC024055 | basic helix-loop-helix transcription factor amos |
| LDEC011458 | forkhead box protein j1-a-like | LDEC004725 | protein hunchback |
| LDEC011779 | paired box protein pax-2-b | LDEC004726 | protein hunchback |
| LDEC003301 | transcription factor ap-2 | LDEC016661 | pou domain protein cf1a |
| LDEC013927 | single-minded homolog 1 isoform x5 | LDEC000968 | heat shock factor isoform x1 |
| LDEC014040 | forkhead box protein n5 isoform x4 | LDEC005435 | protein giant |
| LDEC014127 | protein distal antenna | LDEC005589 | runt |
| LDEC014190 | homeobox protein hox-b4 | LDEC005614 | protein asteroid |

#### Gene expression overlap

Only one esterase (LDEC019310) and one ABC transporter (LDEC004154) were found as significant in both the larval and adult differential expression comparisons. The esterase expression varies widely in susceptible or unexposed populations, with strong upregulation in WI and MI resistant samples. The ABC transporter appeared to have moderate to high constitutive expression in the WI samples, but varied in expression among susceptible, resistant and exposed/unexposed populations. The overlap among larvae and adult differential expression analysis gene lists (**Fig. S33**) showed that shared regulatory changes in gene networks included digestive processes (serine and cysteine-type peptidase activity, and lipid metabolism), but also pathways linked to insecticide detoxification and stress, such as oxidoreductase and dioxygenase activity, heme and iron ion binding, proteolysis and integral component of membrane. We note that the enrichment of lipid metabolism includes an interaction with oxidation-reduction processes and may be related to insecticide detoxification, rather than metabolism per se.

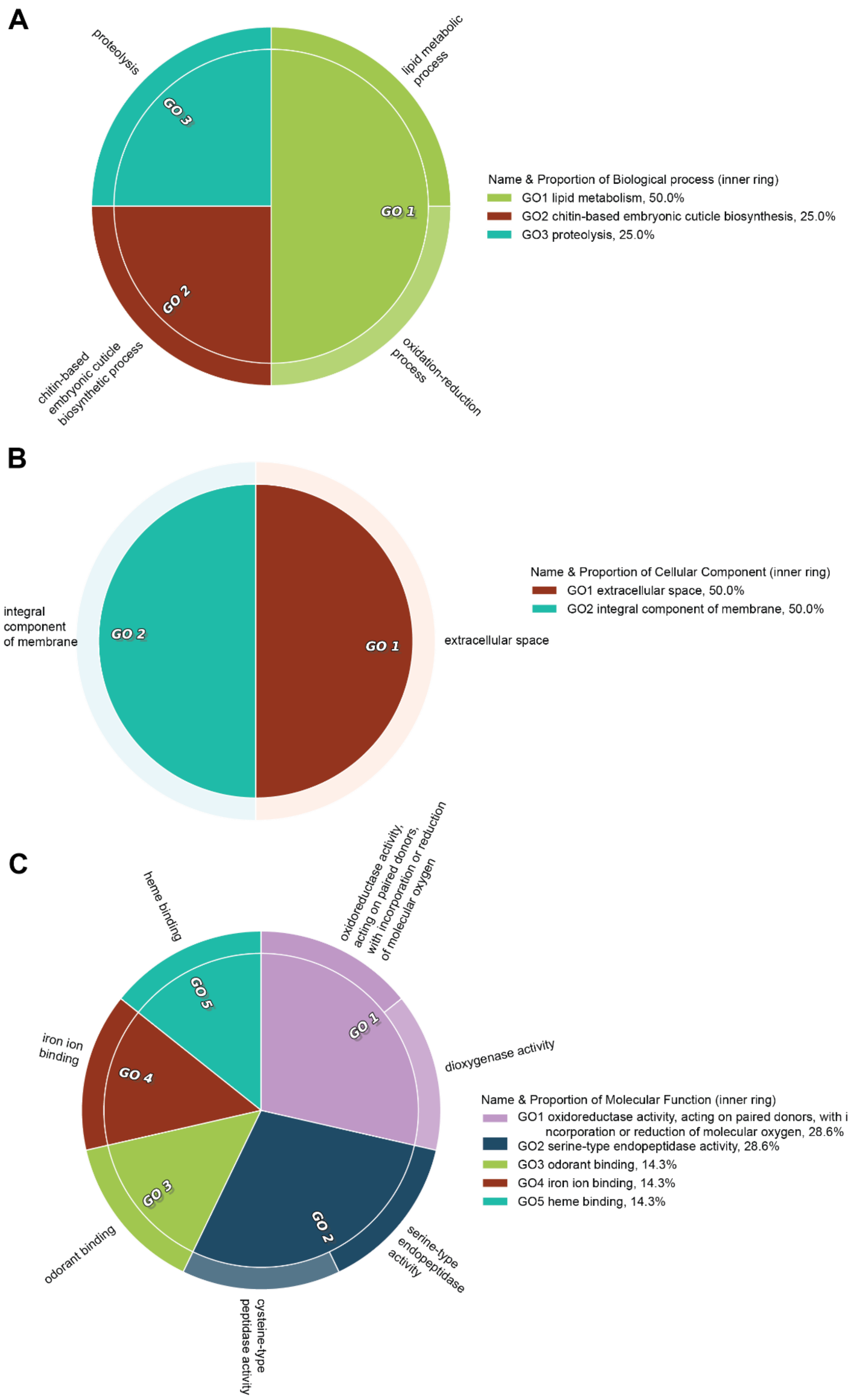

**Figure S33**. *CirGO* plot of overrepresented gene ontology terms shared between the larval and adult among-population differential gene expression analysis. Terms are labeled as A) biological processes, B) cellular components, and C) molecular functions. Each box along the outer ring represents a gene ontology term, and these are grouped into 'superclusters' of related terms based on semantic similarity (grouping is indicated by color and highlighted by a GO label, with the relative importance indicated by the size of the pie slice).

#### Gene ontology overlap among gene expression and selection gene sets

The overlap among enriched gene ontology terms generated from all four analyses, both genome-wide scans of selection (*PCAdapt* and *LFMM*), and both differential gene expression analyses (larvae and adult), resulted in a small set of shared terms (**Figure S44**). However, the six ontology terms were linked to insecticide resistance pathways. Integral component of membrane (GO:0016021) is a broad category that includes genes involved in xenobiotic detoxification, including esterases, cytochrome P-450s, ATP-transporters, as well as chemosensation, hormonal receptors, and membrane-bound ligands. Protein-coding genes in this pathway have been shown to be rapidly evolving in CPB, relative to other *Leptinotarsa* species [18]. Four ontology terms were associated with cytochrome P450 monooxygenase activity (GO:0055114 BP oxidation-reduction process; GO:0016705 MF oxioreductase activity acting on paired donors, with incorporation or reduction of molecular oxygen; GO:0020037 MF heme binding; and GO:0005506 MF iron ion binding), and have been linked to insecticide resistance in multiple mosquito species [27-30], DDT-resistant Drosophila melanogaster [31], and the spider mite *Tetranychus urticae* [32, 33].The last term (GO:0006508 BP proteolysis) involves cellular protein degradation and is a common component of the conserved cellular stress response [34], as is the aforementioned ontology term for oxidation-reduction.

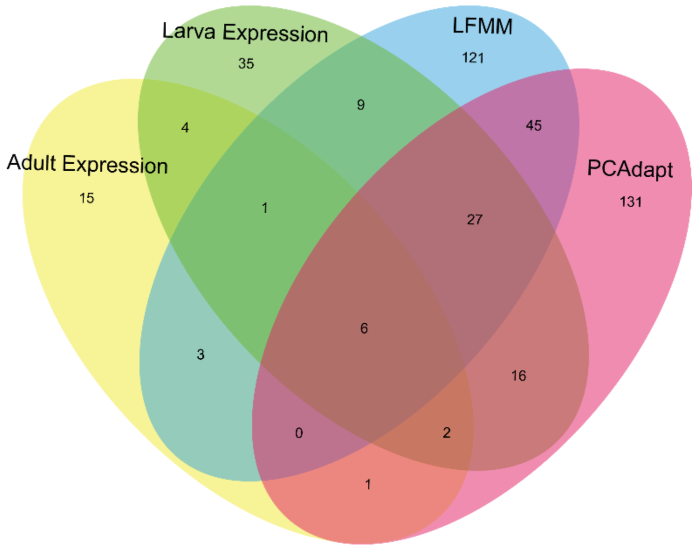

**Figure S34**. Venn diagram showing the overlap of significantly enriched gene ontology terms from the genome scan and gene expression analyses.

### Supplementary Methods

#### Bioassays for resistant samples

Based on the propensity of different populations to develop insecticide resistance and the insecticide usage history in different regions, samples in key regions were collected live for neonicotinoid pesticide bioassays. We sampled six geographically proximate pairs of resistant (R) and susceptible (S) CPB populations: Maine (R) and Vermont (S), New York (R) and New Jersey (S), Maryland (R and S), Michigan (R and S), Wisconsin (R and S), Oregon (R and S). We sequenced genomes from five beetles per population to obtain a total of 30 resistant and 30 susceptible beetle genomes. Populations were chosen based on historical precedent in the literature, and the resistance status of beetles was ascertained by topical exposure to an insecticide (imidacloprid) and from personal observations. Topical exposure was done by applying 1 µL of technical-grade imidacloprid, dissolved in acetone to achieve concentrations ranging from 50 ppm to 2000 ppm, onto the 1$^{st}$ ventral sternite of adult beetles. Beetle resistance status was ascertained three days after exposure, following established protocols [35].

The Maine (R) population came from a conventional potato field, and beetles were healthy after exposure to 500 ppm or 2000 ppm doses of imidacloprid. The Vermont (S) population came from an organic potato field, and beetles were moribund or dead after exposure to a 50 ppm dose of imidacloprid.

The New York (R) population was collected from a commercial potato field in Long Island, where CPB has rapidly and repeatedly evolved resistance to insecticides [36], and beetles were healthy after exposure to a 2000 ppm dose of imidacloprid. The New Jersey (S) population is widely used as an insecticide-susceptible reference population that has been in-colony for >20 years (Thomas Dorsey, pers. comm.), and was obtained from French labs. The $LC_{50}$ of this population has been estimated at ~59 ppm [37].

The Maryland (R) population was obtained from a commercial field in Dorchester County, and beetles survived exposure to a 2000 ppm dose of imidacloprid. The Maryland (S) population was obtained from commercial field in Prince George County, and beetles exhibited knock-down symptoms after exposure to 50 ppm imidacloprid.

The Michigan (R) population was obtained from horsenettle (*Solanum carolinense*) growing along the margins of a conventional potato field, to reduce the direct exposure to insecticides. The beetles were healthy after exposure to a 500 ppm dose of imidacloprid, but moribund after exposure to 2000 ppm. The Michigan (S) population was obtained from a conventional potato field that had recently been sprayed with an insecticide. Beetles throughout the field were either dead or moribund, and the beetles used for sequencing were collected from an unsprayed field edge and assumed to be generally “susceptible”.

The Wisconsin (R) population was obtained from a conventional potato field in an area where CPB populations are considered highly insecticide-resistant (Huseth and Groves 2013, Clements et al. 2016, Crossley et al. 2018), and beetles were healthy after exposure to 500 ppm or 2000 ppm doses of imidacloprid. The Wisconsin (S) population was obtained from a research station that has never applied neonicotinoid insecticides to potato crops. This population is highly susceptible to insecticides (Clements et al. 2017, Crossley et al. 2018), with an imidacloprid $LC_{50}$ estimated at ~27 ppm.

The Oregon (R) population is relatively insecticide-susceptible compared to Midwestern and eastern CPB populations (Olson et al. 2000, Crossley et al. 2018), but was obtained from a potato field that has experienced severe control failures since 2013. Beetles were healthy after exposure to 500 ppm or 1000 ppm doses of imidacloprid. The Oregon (S) population was obtained from conventionally-managed potato on a research station, and beetles were moribund after exposure to a 50 ppm dose of imidacloprid.

#### Genomic Resequencing, Quality Control and Variant Calling

High quality genomic DNA was isolated from beetle thoracic muscle tissue using DNeasy Blood & Tissue kits (Qiagen) and then submitted to the University of Wisconsin-Madison Biotechnology Center. DNA concentration was verified using the Qubit® dsDNA HS Assay Kit (Life Technologies, Grand Island, NY) and 1ug of each sample was sheared using a Covaris M220 Ultrasonicator (Covaris Inc, Woburn, MA, USA) to an average insert size of 550 base pairs (bp). Sizing was verified by Fragment Analyzer (Advanced Analytical Technologies, Inc., Ames, IA, USA). Libraries were prepared according the NEBNext® Ultra™ DNA Library Prep Kit for Illumina (New England Biolabs, Ipswich, MA, USA) with minor modifications. Quality and quantity of the finished libraries were assessed using the Fragment Analyzer and Qubit® dsDNA HS Assay Kit, respectively. Libraries were standardized to 2μM.  Cluster generation was performed using HiSeq PE Cluster Kit v4 cBot kits (Illumina Inc, San Diego, CA, USA). Flow cells were sequenced using paired-end, 125bp sequencing and HiSeq SBS Kit v4 (250 Cycle) (Illumina Inc.) on a HiSeq2500 sequencer. Images were analyzed using the standard Illumina Pipeline, version 1.8.2. Each sample was demutiplexed prior to downstream analysis.

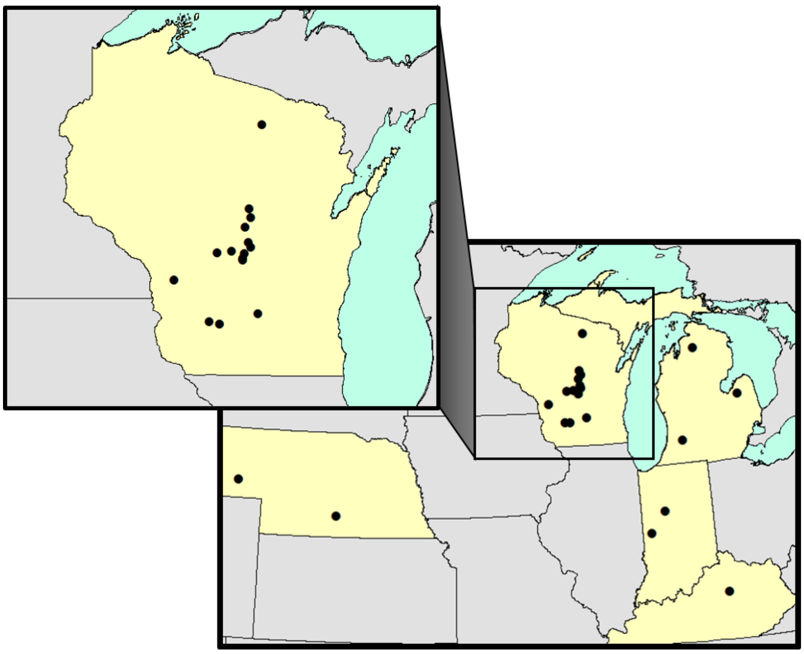

**Figure S35.** Map showing the 24 CPB populations represented in the GBS sampling of Crossley *et al*. [35], from Wisconsin (magnified), Michigan, Indiana, Kentucky and Nebraska.

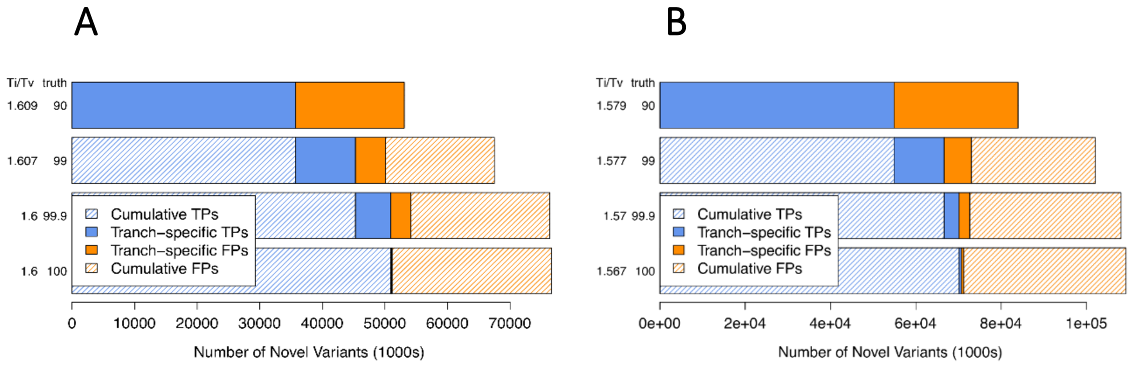

**Figure S36.** Tranche plots depicting variant probability and quality for the A) "CPB" and B) "*Leptinotarsa*" datasets. The number of novel variants is shown on the x-axis, while quality metrics, transition to transversion ratio, and the overall sensitivity, are shown on the y-axis.

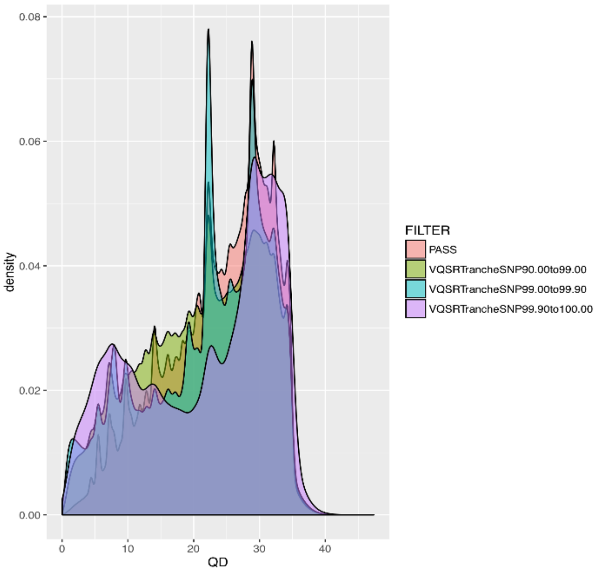

**Figure S37.** Distribution of quality by depth attribute for each inferred SNP in the "CPB" dataset following the VQSR procedure.

#### Selection Tests: *hapFLK* Method

In order to use the multi-point linkage disequilibrium model, *hapFLK* needs the number of haplotype clusters (K) to be specified. Using a custom script (https://github.com/inzilico/kselection), we subsampled the first scaffold and generated 100 datasets containing 10% of artificially, randomly-distributed missing genotypes. We then re-imputed missing alleles and genotypes of every artificial dataset using *fastPHASE* [38], for K= 5, 10, 15, 20, 25 and 40. Finally we calculated the proportion of imputation errors (= 1 - the proportion of correctly imputed alleles and genotypes) for each K value and chose the K value minimizing imputation errors, in our case K=20 (**Fig. S38**). We also compared *hapFLK* results for all aforementioned K values and found our choice of K to be robust. For K=5, 10 and 15, the amount of noise was significantly increased, making detection of candidate regions more difficult. Conversely, for K=25 and above, neither noise nor signal increased, while computational time increased greatly, becoming prohibitive at K=40.

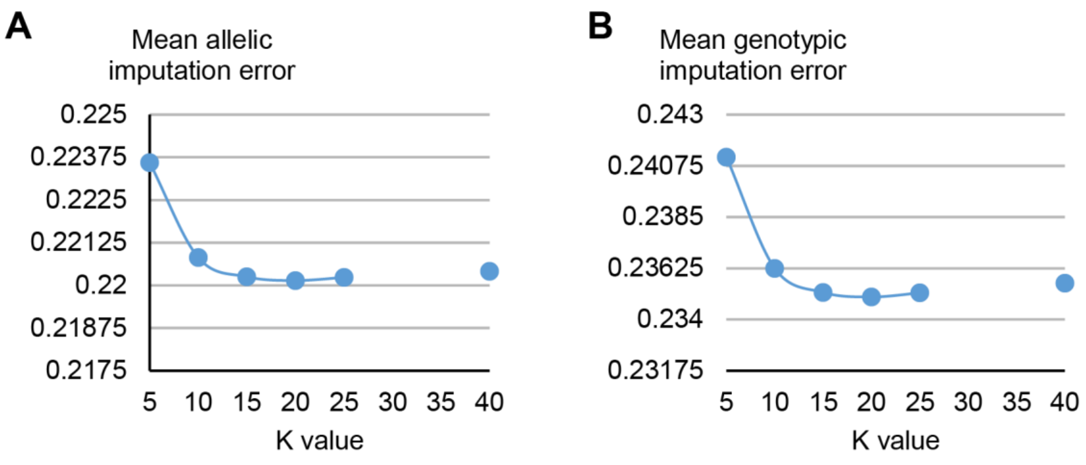

**Figure S38**. A) Allelic and B) *fastPHASE* imputation errors for various K values.

We estimated the kinship matrix and corresponding population tree (**Fig. S39**) for all our samples based on the first 95 scaffolds and then ran *hapFLK* (K=20). The population tree confirmed that Western samples have a distinct evolutionary history, as they belong to a very basal clade which shows a relatively long branch (the longest internal branch). It also confirms the strong allele frequency shifts evident in Europe, New Jersey and the Michigan resistant populations, which all display very long branches. However, the fact that these three populations cluster together might be due to an effect of long branch attraction rather than relatedness. Finally, all remaining pest populations exhibit very short internal branches and are clustered together, even though some sub-structure is still visible. Three distinct sub-clades include several resistant populations (Maryland, New-York and Maine), susceptible populations (Vermont and Maryland) and Midwestern populations.

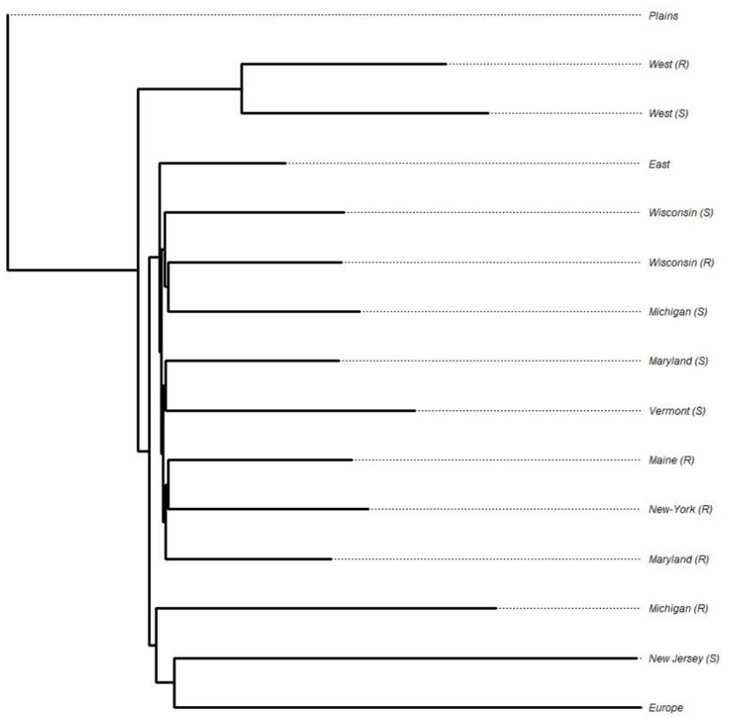

**Figure S39**. Unrooted population tree estimated on the first 95 scaffolds (~21% of the genome). Colors correspond to sampling regions.

Finally, we identified signatures of selection by considering *p-values* at several significance thresholds: α = 0.01, 0.001 and 0.0001 (*e.g.* **Fig. S40**).

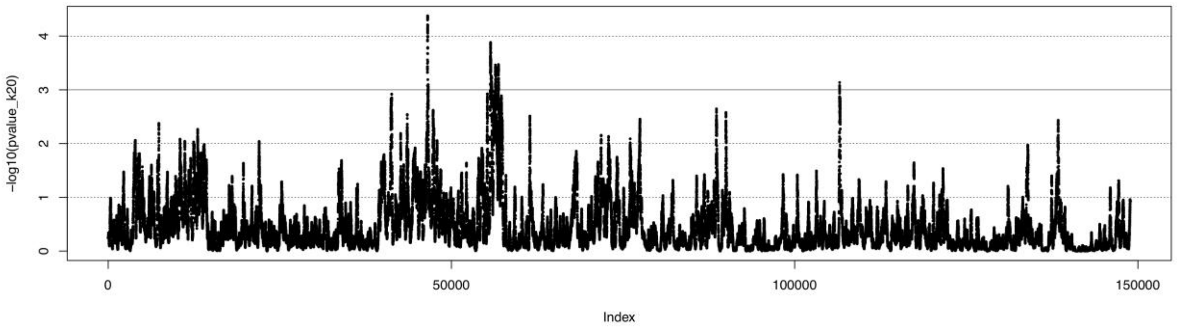

**Figure S40**. Example of *hapFLK* genome scan of the first scaffold (*p-values* are log adjusted).

### Insecticide Resistance Candidate Genes

We obtained a list of candidate insecticide resistance genes using manually and Blast2GO-annotated gene models in the *L. decemlineata* OGS v1.1, along with custom R scripts. We looked for genes associated with known mechanisms of insecticide resistance (**Table S14**): metabolic detoxification including cytochrome p450s (CYPs), esterases, Glutathione S-transferases (GSTs) and ATP-binding cassette (ABC) transporters [39]; target-site insensitivity including most of the modes of actions classified by the *Insecticide Resistance Action Committee* (IRAC; http://www.irac-online.org/modes-of-action/); and reduced cuticular penetration including genes involved chitin production or cuticle development. Gene ontology terms associated with these genes, including biological processes, cellular components, and molecular functions, are clustered using the simRel score for functional similarity and plotted using a *CirGO* plot (**Fig. S41**) [40].

**Table S14.** Candidate insecticide resistance genes in the *Leptinotarsa decemlineata* genome.

| Gene ID | Gene Annotation | Gene ID | Gene Annotation |
|---|---|---|---|
| LDEC018534 | cytochrome P450 | LDEC013136 | major facilitator superfamily domain-containing protein 6 |
| LDEC018533 | cytochrome P450 | LDEC003251 | major facilitator superfamily domain-containing protein 6 |
| LDEC019185 | cytochrome P450 | LDEC013524 | major facilitator superfamily domain-containing protein 6 |
| LDEC019186 | cytochrome P450 | LDEC014298 | major facilitator superfamily domain-containing protein 6 |
| LDEC019187 | cytochrome P450 | LDEC004274 | major facilitator superfamily domain-containing protein 6 |
| LDEC019188 | cytochrome P450 | LDEC010033 | major facilitator superfamily domain-containing protein 6-a |
| LDEC019189 | cytochrome P450 | LDEC023938 | major facilitator superfamily domain-containing protein 6-a |
| LDEC024466 | cytochrome P450 | LDEC009079 | major facilitator superfamily domain-containing protein 8 |
| LDEC019765 | cytochrome P450 | LDEC003558 | major facilitator superfamily domain-containing protein 9-like |
| LDEC019766 | cytochrome P450 | LDEC022720 | atp-binding cassette sub-family a member 3 |
| LDEC019769 | cytochrome P450 | LDEC024718 | atp-binding cassette sub-family a member 3 isoform x1 |
| LDEC020420 | cytochrome P450 | LDEC008896 | atp-binding cassette sub-family a member 3 isoform x1 |
| LDEC020421 | cytochrome P450 | LDEC023229 | atp-binding cassette sub-family a member 3 isoform x2 |
| LDEC020526 | cytochrome P450 | LDEC008893 | atp-binding cassette sub-family a member 3-like protein |
| LDEC007597 | cytochrome P450 | LDEC024249 | atp-binding cassette sub-family a member 3-like protein |
| LDEC007598 | cytochrome P450 | LDEC005667 | atp-binding cassette sub-family a member 3-like protein |
| LDEC007599 | cytochrome P450 | LDEC015008 | atp-binding cassette sub-family a member 5-like isoform x1 |
| LDEC021334 | cytochrome P450 | LDEC015007 | atp-binding cassette sub-family a member 5-like isoform x2 |
| LDEC022309 | cytochrome p450 | LDEC024415 | ATP-binding cassette sub-family B member mitochondrial |
| LDEC022856 | cytochrome p450 | LDEC019485 | ATP-binding cassette sub-family B member mitochondrial |
| LDEC022983 | cytochrome p450 | LDEC007355 | ATP-binding cassette sub-family B member mitochondrial |
| LDEC009934 | cytochrome p450 | LDEC022912 | atp-binding cassette sub-family b member mitochondrial |
| LDEC009935 | cytochrome p450 | LDEC010427 | atp-binding cassette sub-family b member mitochondrial |
| LDEC023239 | cytochrome p450 | LDEC024539 | ATP-binding cassette sub-family B member mitochondrial isoform X1 |
| LDEC023330 | cytochrome p450 | LDEC002144 | atp-binding cassette sub-family b member mitochondrial-like |
| LDEC023573 | cytochrome p450 | LDEC000891 | atp-binding cassette sub-family d member 2 |
| LDEC013540 | cytochrome p450 | LDEC007110 | ATP-binding cassette sub-family D member 3 |
| LDEC023780 | cytochrome p450 | LDEC007111 | ATP-binding cassette sub-family D member 3 |
| LDEC014467 | cytochrome p450 | LDEC001567 | ATP-binding cassette sub-family E member 1 |
| LDEC015048 | cytochrome p450 | LDEC002470 | atp-binding cassette sub-family f member 1 |
| LDEC015049 | cytochrome p450 | LDEC005120 | atp-binding cassette sub-family f member 2 |
| LDEC015052 | cytochrome p450 | LDEC023305 | atp-binding cassette sub-family f member 3 |
| LDEC015218 | cytochrome p450 | LDEC003451 | atp-binding cassette sub-family f member 3 |
| LDEC015220 | cytochrome p450 | LDEC004565 | atp-binding cassette sub-family f member partial |
| LDEC015420 | cytochrome p450 | LDEC018766 | ATP-binding cassette sub-family G member 1 |
| LDEC004499 | cytochrome p450 | LDEC002861 | atp-binding cassette sub-family g member 1 |
| LDEC004502 | cytochrome p450 | LDEC020054 | ATP-binding cassette sub-family G member 1-like |
| LDEC004503 | cytochrome p450 | LDEC002775 | atp-binding cassette sub-family g member 1-like |
| LDEC015922 | cytochrome p450 | LDEC010324 | atp-binding cassette sub-family g member 4 |

| | | | |
|---|---|---|---|
| LDEC016271 | cytochrome p450 | LDEC002867 | atp-binding cassette sub-family g member 4 |
| LDEC016272 | cytochrome p450 | LDEC005346 | atp-binding cassette sub-family g member 4 isoform x1 |
| LDEC018119 | cytochrome p450 | LDEC005530 | atp-binding cassette sub-family g member 4-like |
| LDEC005703 | cytochrome p450 | LDEC017668 | atp-binding cassette sub-family g member 5 |
| LDEC019183 | cytochrome P450 [Tribolium castaneum] | LDEC017669 | atp-binding cassette sub-family g member 8 |
| LDEC019189 | cytochrome P450 [Tribolium castaneum] | LDEC021233 | multidrug resistance 1A |
| LDEC006084 | cytochrome P450 [Tribolium castaneum] | LDEC008906 | multidrug resistance protein 1a |
| LDEC011286 | cytochrome p450 18a1 | LDEC008907 | multidrug resistance protein 1b isoform x1 |
| LDEC016277 | cytochrome p450 2l1 | LDEC008908 | multidrug resistance protein 1b isoform x1 |
| LDEC011287 | cytochrome p450 306a1 | LDEC008909 | multidrug resistance protein 1b isoform x1 |
| LDEC006897 | cytochrome P450 307a1-like | LDEC014301 | multidrug resistance protein 1b isoform x1 |
| LDEC021933 | cytochrome P450 307a1-like | LDEC021280 | multidrug resistance-associated |
| LDEC000648 | cytochrome p450 49a1 | LDEC019090 | Multidrug resistance-associated 1 |
| LDEC008545 | cytochrome p450 4c1- partial | LDEC021940 | multidrug resistance-associated 1 isoform X1 |
| LDEC009365 | cytochrome p450 4c1- partial | LDEC021941 | multidrug resistance-associated 1 isoform X1 |
| LDEC005857 | cytochrome P450 4C1-like | LDEC020972 | multidrug resistance-associated 4 |
| LDEC006858 | cytochrome P450 4C1-like | LDEC021794 | multidrug resistance-associated 4 |
| LDEC006857 | cytochrome P450 4C1-like | LDEC020530 | multidrug resistance-associated 4-like |
| LDEC023594 | cytochrome p450 4c1-like | LDEC021235 | multidrug resistance-associated 4-like |
| LDEC023726 | cytochrome p450 4c1-like | LDEC021236 | multidrug resistance-associated 4-like |
| LDEC015420 | cytochrome p450 4c1-like | LDEC021279 | multidrug resistance-associated 4-like |
| LDEC016357 | cytochrome p450 4c1-like | LDEC021627 | multidrug resistance-associated 4-like |
| LDEC017468 | cytochrome p450 4c1-like | LDEC022210 | multidrug resistance-associated protein |
| LDEC006438 | cytochrome P450 4C1-like isoform X1 | LDEC022769 | multidrug resistance-associated protein |
| LDEC003194 | cytochrome p450 4g15 | LDEC022949 | multidrug resistance-associated protein 1 |
| LDEC022711 | cytochrome p450 4g15-like | LDEC022533 | multidrug resistance-associated protein 4 |
| LDEC003603 | cytochrome p450 4g15-like | LDEC022534 | multidrug resistance-associated protein 4 |
| LDEC024738 | cytochrome p450 6a2 | LDEC009711 | multidrug resistance-associated protein 4 |
| LDEC014519 | cytochrome p450 6a2 | LDEC023815 | multidrug resistance-associated protein 4 |
| LDEC004501 | cytochrome p450 6a2 | LDEC004150 | multidrug resistance-associated protein 4 |
| LDEC016518 | cytochrome p450 6a2 isoform x1 | LDEC004151 | multidrug resistance-associated protein 4 |
| LDEC013538 | cytochrome p450 6bq10 | LDEC004152 | multidrug resistance-associated protein 4 |
| LDEC005460 | cytochrome p450 6bq10 | LDEC004153 | multidrug resistance-associated protein 4 |
| LDEC005462 | cytochrome p450 6bq10 | LDEC004154 | multidrug resistance-associated protein 4 |
| LDEC011526 | cytochrome p450 6bq11 | LDEC004155 | multidrug resistance-associated protein 4 |
| LDEC013537 | cytochrome p450 6bq11 | LDEC004858 | multidrug resistance-associated protein 4 |
| LDEC004500 | cytochrome p450 6bq8 | LDEC004859 | multidrug resistance-associated protein 4 |
| LDEC001002 | cytochrome P450 6k1-like | LDEC004861 | multidrug resistance-associated protein 4 |
| LDEC020576 | cytochrome P450 6k1-like | LDEC005086 | multidrug resistance-associated protein 4 |
| LDEC007699 | cytochrome P450 6k1-like | LDEC005089 | multidrug resistance-associated protein 4 |
| LDEC007962 | cytochrome P450 6k1-like | LDEC005090 | multidrug resistance-associated protein 4 |
| LDEC007963 | cytochrome P450 6k1-like | LDEC017374 | multidrug resistance-associated protein 4 |
| LDEC007966 | cytochrome P450 6k1-like | LDEC022138 | multidrug resistance-associated protein 4-like |
| LDEC022006 | cytochrome p450 6k1-like | LDEC022365 | multidrug resistance-associated protein 4-like |
| LDEC008391 | cytochrome p450 6k1-like | LDEC022765 | multidrug resistance-associated protein 4-like |
| LDEC011604 | cytochrome p450 6k1-like | LDEC022945 | multidrug resistance-associated protein 4-like |
| LDEC007964 | cytochrome P450 6k1-like isoform X3 | LDEC002289 | multidrug resistance-associated protein 4-like |

| | | | |
|---|---|---|---|
| LDEC011260 | cytochrome p450 6k1-like isoform x3 | LDEC023763 | multidrug resistance-associated protein 4-like |
| LDEC024396 | cytochrome P450 9e2 [Tribolium castaneum] | LDEC000857 | multidrug resistance-associated protein 4-like |
| LDEC019767 | cytochrome P450 9e2-like | LDEC003183 | multidrug resistance-associated protein 4-like isoform x1 |
| LDEC020527 | cytochrome P450 9e2-like | LDEC013093 | multidrug resistance-associated protein 4-like isoform x1 |
| LDEC006940 | cytochrome P450 9e2-like | LDEC003554 | multidrug resistance-associated protein 4-like isoform x1 |
| LDEC021333 | cytochrome P450 9e2-like | LDEC009620 | multidrug resistance-associated protein 7 |
| LDEC021882 | cytochrome P450 9e2-like | LDEC020973 | probable multidrug resistance-associated lethal(2)03659 |
| LDEC015050 | cytochrome p450 9e2-like | LDEC024636 | probable multidrug resistance-associated lethal(2)03659 |
| LDEC019333 | Cytochrome P450 mitochondrial | LDEC021234 | probable multidrug resistance-associated lethal(2)03659 |
| LDEC021899 | cytochrome P450 mitochondrial | LDEC021955 | probable multidrug resistance-associated lethal(2)03659 |
| LDEC010650 | cytochrome p450 mitochondrial | LDEC022154 | probable multidrug resistance-associated protein lethal 03659 |
| LDEC000205 | cytochrome p450 partial | LDEC002110 | probable multidrug resistance-associated protein lethal 03659 |
| LDEC009936 | cytochrome p450 partial | LDEC002111 | probable multidrug resistance-associated protein lethal 03659 |
| LDEC014468 | cytochrome p450 partial | LDEC002116 | probable multidrug resistance-associated protein lethal 03659 |
| LDEC005461 | cytochrome p450 partial | LDEC002119 | probable multidrug resistance-associated protein lethal 03659 |
| LDEC008373 | cytochrome p450-like protein | LDEC023040 | probable multidrug resistance-associated protein lethal 03659 |
| LDEC003195 | cytochrome p450-like protein | LDEC023045 | probable multidrug resistance-associated protein lethal 03659 |
| LDEC004541 | cytochrome p450-like protein | LDEC023046 | probable multidrug resistance-associated protein lethal 03659 |
| LDEC004542 | cytochrome p450-like protein | LDEC023094 | probable multidrug resistance-associated protein lethal 03659 |
| LDEC009201 | probable cytochrome p450 303a1 | LDEC002518 | probable multidrug resistance-associated protein lethal 03659 |
| LDEC009207 | probable cytochrome p450 303a1 | LDEC011867 | probable multidrug resistance-associated protein lethal 03659 |
| LDEC016273 | probable cytochrome p450 305a1 | LDEC012031 | probable multidrug resistance-associated protein lethal 03659 |
| LDEC016278 | probable cytochrome p450 305a1 | LDEC023743 | probable multidrug resistance-associated protein lethal 03659 |
| LDEC000768 | probable cytochrome p450 49a1 isoform x1 | LDEC014173 | probable multidrug resistance-associated protein lethal 03659 |
| LDEC004819 | probable cytochrome p450 49a1 isoform x1 | LDEC024128 | probable multidrug resistance-associated protein lethal 03659 |
| LDEC010280 | probable cytochrome p450 4aa1 | LDEC024132 | probable multidrug resistance-associated protein lethal 03659 |
| LDEC010662 | probable cytochrome p450 4s3 | LDEC005087 | probable multidrug resistance-associated protein lethal 03659 |
| LDEC011343 | probable cytochrome p450 6a13 | LDEC017372 | probable multidrug resistance-associated protein lethal 03659 |
| LDEC005459 | probable cytochrome p450 6a14 | LDEC017375 | probable multidrug resistance-associated protein lethal 03659 |
| LDEC022008 | probable cytochrome p450 6a14 isoform x2 | LDEC017376 | probable multidrug resistance-associated protein lethal 03659 |
| LDEC021046 | probable cytochrome P450 mitochondrial | LDEC017378 | probable multidrug resistance-associated protein lethal 03659 |
| LDEC008256 | probable cytochrome P450 mitochondrial | LDEC017818 | probable multidrug resistance-associated protein lethal 03659 |
| LDEC008257 | probable cytochrome P450 mitochondrial | LDEC017905 | probable multidrug resistance-associated protein lethal 03659 |
| LDEC013790 | probable cytochrome p450 mitochondrial | LDEC018022 | probable multidrug resistance-associated protein lethal 03659 |
| LDEC004817 | probable cytochrome p450 mitochondrial | LDEC005645 | probable multidrug resistance-associated protein lethal 03659 |
| LDEC004818 | probable cytochrome p450 mitochondrial | LDEC022532 | probable multidrug resistance-associated protein lethal 03659 isoform x1 |
| LDEC006766 | glutathione S-transferase | LDEC009692 | probable multidrug resistance-associated protein lethal 03659 isoform x1 |
| LDEC020265 | Glutathione S-transferase | LDEC012767 | probable multidrug resistance-associated protein lethal 03659 isoform x1 |
| LDEC009779 | glutathione s-transferase | LDEC005088 | probable multidrug resistance-associated protein lethal 03659 isoform x1 |

| | | | |
|---|---|---|---|
| LDEC014129 | glutathione s-transferase | LDEC017379 | probable multidrug resistance-associated protein lethal 03659 isoform x1 |
| LDEC001117 | glutathione S-transferase 1 | LDEC001030 | cuticle 19-like |
| LDEC001120 | glutathione S-transferase 1 | LDEC001031 | cuticle 19-like |
| LDEC015214 | glutathione s-transferase 1 | LDEC001029 | cuticle 7 |
| LDEC015216 | glutathione s-transferase 1 isoform x2 | LDEC001025 | cuticle 7-like |
| LDEC007489 | Glutathione S-transferase 1-1 | LDEC001021 | cuticle 8 |
| LDEC022984 | glutathione s-transferase 1-1 | LDEC001016 | cuticle 8-like |
| LDEC001117 | glutathione S-transferase 1-like | LDEC001026 | cuticle 8-like |
| LDEC001121 | glutathione S-transferase 1-like | LDEC001027 | cuticle 8-like |
| LDEC001122 | glutathione S-transferase 1-like | LDEC020438 | cuticle -like |
| LDEC001123 | glutathione S-transferase 1-like | LDEC001803 | cuticle protein |
| LDEC019759 | glutathione S-transferase 1-like | LDEC009542 | cuticle protein |
| LDEC019760 | glutathione S-transferase 1-like | LDEC011039 | cuticle protein |
| LDEC004485 | glutathione s-transferase c-terminal domain-containing protein homolog | LDEC011040 | cuticle protein |
| LDEC001116 | glutathione S-transferase epsilon | LDEC011041 | cuticle protein |
| LDEC001118 | glutathione S-transferase epsilon | LDEC011042 | cuticle protein |
| LDEC001119 | glutathione S-transferase epsilon | LDEC011043 | cuticle protein |
| LDEC024589 | glutathione S-transferase GST2 | LDEC013169 | cuticle protein |
| LDEC023323 | glutathione s-transferase isoform d | LDEC015588 | cuticle protein |
| LDEC008356 | glutathione s-transferase omega-1-like | LDEC003386 | cuticle protein 19 |
| LDEC012947 | glutathione s-transferase theta-1 | LDEC003392 | cuticle protein 19 |
| LDEC020798 | glutathione S-transferase theta-1-like | LDEC022168 | cuticle protein 19-like |
| LDEC014130 | glutathione s-transferase-like | LDEC003393 | cuticle protein 19-like |
| LDEC004455 | glutamate receptor 1 | LDEC024307 | cuticle protein 19-like |
| LDEC004457 | glutamate receptor 1 | LDEC018268 | cuticle protein 19-like |
| LDEC004458 | glutamate receptor 1 | LDEC012198 | cuticle protein 21 |
| LDEC022173 | glutamate receptor 1-like protein | LDEC015109 | cuticle protein 21-like |
| LDEC009921 | glutamate receptor 1-like protein | LDEC015110 | cuticle protein 21-like |
| LDEC018044 | glutamate receptor 1-like protein | LDEC003627 | cuticle protein 64 |
| LDEC018929 | glutamate receptor 2-like | LDEC018266 | cuticle protein 7 |
| LDEC009862 | glutamate receptor 2-like isoform x4 | LDEC016286 | cuticle protein 76-like |
| LDEC001378 | glutamate receptor delta-1 subunit | LDEC016288 | cuticle protein 76-like |
| LDEC001376 | Glutamate receptor delta-1 subunit | LDEC009667 | cuticle protein 7-like |
| LDEC023971 | glutamate receptor delta-1 subunit | LDEC003387 | cuticle protein 7-like |
| LDEC001377 | Glutamate receptor delta-2 subunit | LDEC023335 | cuticle protein 8-like |
| LDEC014994 | glutamate receptor delta-2 subunit | LDEC003397 | cuticle protein 8-like |
| LDEC015416 | glutamate receptor delta-2 subunit | LDEC002961 | cuticle protein isoform a-like |
| LDEC017754 | glutamate receptor kainate 1 | LDEC011470 | cuticlin- partial |
| LDEC018518 | Glutamate receptor kainate 1 [Tribolium castaneum] | LDEC011468 | cuticlin-1 |
| LDEC018519 | Glutamate receptor kainate 1 [Tribolium castaneum] | LDEC016323 | cuticlin-1 |
| LDEC018522 | Glutamate receptor kainate 1 [Tribolium castaneum] | LDEC016622 | cuticlin-1-like protein |
| LDEC022809 | glutamate receptor kainate 1-like protein | LDEC004243 | cuticular precursor |
| LDEC019234 | glutamate receptor kainate 2 | LDEC008714 | cuticular protein |
| LDEC019236 | glutamate receptor kainate 2 | LDEC008715 | cuticular protein |
| LDEC006963 | Glutamate receptor kainate 2 | LDEC012195 | cuticular protein |
| LDEC021982 | Glutamate receptor kainate 2 | LDEC014759 | cuticular protein |

| | | | |
|---|---|---|---|
| LDEC009920 | glutamate receptor kainate 2 | LDEC004413 | cuticular protein 4 |
| LDEC015534 | glutamate receptor kainate 2 | LDEC004414 | cuticular protein 4 |
| LDEC024033 | glutamate receptor kainate 2 | LDEC000974 | cuticular protein 4 |
| LDEC018045 | glutamate receptor kainate 2 | LDEC014174 | cuticular protein 56f |
| LDEC019311 | glutamate receptor kainate 2 isoform X1 | LDEC015105 | cuticular protein 5c |
| LDEC007558 | glutamate receptor kainate 2 isoform X1 | LDEC004066 | cuticular protein analogous to peritrophins 1-b |
| LDEC007560 | glutamate receptor kainate 2 isoform X1 | LDEC005020 | cuticular protein analogous to peritrophins 1-b |
| LDEC017418 | glutamate receptor kainate 2 isoform x1 | LDEC017902 | cuticular protein analogous to peritrophins 1-f precursor |
| LDEC023278 | glutamate receptor kainate 2 isoform x2 | LDEC014693 | cuticular protein analogous to peritrophins 1-j precursor |
| LDEC024021 | glutamate receptor kainate 2 isoform x2 | LDEC003587 | cuticular protein analogous to peritrophins 3-a2 precursor |
| LDEC015532 | glutamate receptor kainate 2 isoform x3 | LDEC003585 | cuticular protein analogous to peritrophins 3-d1 |
| LDEC012786 | glutamate receptor kainate 2 isoform x4 | LDEC010847 | cuticular protein rr-2 family |
| LDEC015313 | glutamate receptor kainate 2 isoform x8 | LDEC010849 | cuticular protein rr-2 family |
| LDEC007559 | glutamate receptor kainate 2-like | LDEC012667 | cuticular protein rr-2 family |
| LDEC022174 | glutamate receptor kainate 2-like | LDEC012668 | cuticular protein rr-2 family |
| LDEC015533 | glutamate receptor kainate 2-like | LDEC003398 | cuticular protein rr-2 motif 130 |
| LDEC023413 | glutamate receptor kainate 2-like isoform x11 | LDEC008054 | cuticular RR-2 family |
| LDEC019232 | glutamate receptor kainate 2-like isoform X2 | LDEC004412 | tweedle motif cuticular protein 2 |
| LDEC019235 | glutamate receptor kainate 2-like isoform X2 | LDEC001813 | flexible cuticle protein 12 |
| LDEC022175 | glutamate receptor kainate 2-like isoform x2 | LDEC010805 | flexible cuticle protein 12 |
| LDEC023412 | glutamate receptor kainate 2-like isoform x5 | LDEC001804 | flexible cuticle protein 12-like |
| LDEC022643 | glutamate receptor kainate 2-like protein | LDEC001805 | flexible cuticle protein 12-like |
| LDEC016633 | glutamate receptor kainate 2-like protein | LDEC010804 | flexible cuticle protein 12-like |
| LDEC016634 | glutamate receptor kainate 2-like protein | LDEC006247 | larval cuticle 8-like |
| LDEC005024 | glutamate receptor kainate 2-like protein | LDEC024474 | larval cuticle A2B-like |
| LDEC019627 | glutamate receptor kainate 5 | LDEC021054 | larval cuticle A2B-like |
| LDEC019628 | glutamate receptor kainate 5 | LDEC006245 | larval cuticle LCP-30 |
| LDEC020708 | glutamate receptor kainate 5 | LDEC006246 | larval cuticle LCP-30 |
| LDEC008011 | Glutamate receptor NMDA 1 | LDEC001825 | larval cuticle protein |
| LDEC004472 | glutamate receptor nmda 2b isoform x1 | LDEC005242 | larval cuticle protein 1 |
| LDEC004469 | glutamate receptor nmda 2b isoform x2 | LDEC001808 | larval cuticle protein 8-like |
| LDEC020195 | glutamate receptor NMDA 3A-like | LDEC001814 | larval cuticle protein 8-like |
| LDEC020795 | glutamate receptor U1 | LDEC010803 | larval cuticle protein 8-like |
| LDEC006508 | Glutamate receptor-interacting 2 | LDEC001734 | larval cuticle protein a2b |
| LDEC006510 | Glutamate receptor-interacting 2 | LDEC015102 | larval cuticle protein a2b |
| LDEC006511 | Glutamate receptor-interacting 2 | LDEC011031 | larval cuticle protein a2b-like |
| LDEC006514 | Glutamate receptor-interacting 2 | LDEC011032 | larval cuticle protein a2b-like |
| LDEC000434 | glutamate receptor-like | LDEC011033 | larval cuticle protein a2b-like |
| LDEC000294 | probable glutamate receptor | LDEC011036 | larval cuticle protein a2b-like |
| LDEC014124 | probable glutamate receptor isoform x1 | LDEC011038 | larval cuticle protein a2b-like |
| LDEC013845 | probable glutamate receptor isoform x2 | LDEC023912 | larval cuticle protein a2b-like |
| LDEC010137 | chloride channel protein 2 isoform x1 | LDEC015096 | larval cuticle protein a2b-like |
| LDEC010129 | chloride channel protein 2 isoform x2 | LDEC015097 | larval cuticle protein a2b-like |
| LDEC010130 | chloride channel protein 2 isoform x2 | LDEC015098 | larval cuticle protein a2b-like |
| LDEC010134 | chloride channel protein 2 isoform x2 | LDEC015104 | larval cuticle protein a2b-like |
| LDEC010135 | chloride channel protein 2 isoform x2 | LDEC015106 | larval cuticle protein a2b-like |

| | | | |
|---|---|---|---|
| LDEC010136 | chloride channel protein 2 isoform x2 | LDEC015107 | larval cuticle protein a2b-like |
| LDEC017002 | chloride intracellular channel exc-4 | LDEC004877 | larval cuticle protein a2b-like |
| LDEC011784 | carboxylesterase | LDEC004540 | larval cuticle protein a3a isoform x1 |
| LDEC011789 | carboxylesterase | LDEC004543 | larval cuticle protein a3a isoform x1 |
| LDEC006847 | carboxylesterase 1E | LDEC004544 | larval cuticle protein a3a isoform x1 |
| LDEC013802 | carboxylesterase 1e | LDEC018315 | larval cuticle protein a3a-like |
| LDEC021746 | Carboxylesterase 3 | LDEC001829 | larval cuticle protein lcp-17 |
| LDEC015960 | carboxylesterase 5a | LDEC001811 | larval cuticle protein lcp-30 |
| LDEC015961 | carboxylesterase 5a | LDEC000824 | larval cuticle protein lcp-30 |
| LDEC018986 | carboxylesterase CXE6 | LDEC000825 | larval cuticle protein lcp-30 |
| LDEC011822 | carboxylesterase cxe6 | LDEC012201 | larval pupal cuticle protein h1c |
| LDEC016728 | carboxylesterase cxe6 | LDEC012202 | larval pupal cuticle protein h1c |
| LDEC019310 | esterase | LDEC012203 | larval pupal cuticle protein h1c |
| LDEC019313 | esterase | LDEC012204 | larval pupal cuticle protein h1c |
| LDEC019314 | esterase | LDEC012205 | larval pupal cuticle protein h1c |
| LDEC019315 | esterase | LDEC024732 | larval pupal cuticle protein h1c-like |
| LDEC019316 | esterase | LDEC018687 | voltage-dependent calcium channel gamma-7 subunit [Cephus cinctus] |
| LDEC019607 | esterase | LDEC000112 | voltage-dependent calcium channel subunit alpha-2 delta-3 |
| LDEC019968 | esterase | LDEC016259 | voltage-dependent calcium channel subunit alpha-2 delta-3 |
| LDEC020362 | esterase | LDEC005309 | voltage-dependent calcium channel subunit alpha-2 delta-3 |
| LDEC020662 | esterase | LDEC019417 | voltage-dependent calcium channel subunit alpha-2 delta-3 isoform X1 |
| LDEC020663 | esterase | LDEC020521 | voltage-dependent calcium channel subunit alpha-2 delta-3 isoform X1 |
| LDEC020664 | esterase | LDEC014561 | voltage-dependent calcium channel subunit alpha-2 delta-3 isoform x1 |
| LDEC021268 | esterase | LDEC016260 | voltage-dependent calcium channel subunit alpha-2 delta-3 isoform x1 |
| LDEC022525 | esterase | LDEC014562 | voltage-dependent calcium channel subunit alpha-2 delta-3 isoform x2 |
| LDEC022531 | esterase | LDEC019416 | voltage-dependent calcium channel subunit alpha-2 delta-3 isoform X3 |
| LDEC009236 | esterase | LDEC013402 | voltage-dependent calcium channel type a subunit alpha-1 |
| LDEC009237 | esterase | LDEC013406 | voltage-dependent calcium channel type a subunit alpha-1 |
| LDEC023088 | esterase | LDEC013412 | voltage-dependent calcium channel type a subunit alpha-1 isoform x1 |
| LDEC023090 | esterase | LDEC013403 | voltage-dependent calcium channel type a subunit alpha-1-like protein |
| LDEC011641 | esterase | LDEC013407 | voltage-dependent calcium channel type a subunit alpha-1-like protein |
| LDEC011642 | esterase | LDEC013408 | voltage-dependent calcium channel type a subunit alpha-1-like protein |
| LDEC012570 | esterase | LDEC013410 | voltage-dependent calcium channel type a subunit alpha-1-like protein |
| LDEC012571 | esterase | LDEC013413 | voltage-dependent calcium channel type a subunit alpha-1-like protein |
| LDEC012574 | esterase | LDEC013414 | voltage-dependent calcium channel type a subunit alpha-1-like protein |
| LDEC012575 | esterase | LDEC021583 | Voltage-dependent calcium channel type D subunit alpha-1 |
| LDEC012578 | esterase | LDEC021584 | Voltage-dependent calcium channel type D subunit alpha-1 |
| LDEC013097 | esterase | LDEC015955 | voltage-dependent calcium channel type d subunit alpha-1-like protein |
| LDEC014121 | esterase | LDEC001186 | voltage-dependent L-type calcium channel subunit beta-2 isoform X9 |
| LDEC014122 | esterase | LDEC004591 | voltage-dependent t-type calcium channel subunit alpha-1g |
| LDEC014125 | esterase | LDEC004595 | voltage-dependent t-type calcium channel subunit alpha-1g isoform x2 |

| | | | |
|---|---|---|---|
| LDEC015319 | esterase | LDEC004596 | voltage-dependent t-type calcium channel subunit alpha-1g isoform x2 |
| LDEC015320 | esterase | LDEC004597 | voltage-dependent t-type calcium channel subunit alpha-1g isoform x2 |
| LDEC017038 | esterase | LDEC014585 | voltage-dependent t-type calcium channel subunit alpha-1g isoform x4 |
| LDEC017039 | esterase | LDEC004593 | voltage-dependent t-type calcium channel subunit alpha-1g-like |
| LDEC024201 | esterase | LDEC004594 | voltage-dependent t-type calcium channel subunit alpha-1g-like isoform x1 |
| LDEC018118 | esterase | LDEC020116 | NA(involved in ryanodine reception) |
| LDEC005856 | esterase [Tribolium castaneum] | LDEC016101 | nicotinic acetylcholine receptor a9 subunit |
| LDEC019309 | esterase E4-like | LDEC016788 | nicotinic acetylcholine receptor subunit a2 |
| LDEC011645 | esterase e4-like | LDEC016741 | nicotinic acetylcholine receptor subunit alpha10 |
| LDEC011785 | esterase e4-like | LDEC020580 | nicotinic acetylcholine receptor subunit alpha4 |
| LDEC011786 | esterase e4-like | LDEC007707 | nicotinic acetylcholine receptor subunit alpha4 |
| LDEC012342 | esterase e4-like | LDEC006111 | nicotinic acetylcholine receptor subunit alpha5 [Tribolium castaneum] |
| LDEC014547 | esterase e4-like | LDEC000437 | nicotinic acetylcholine receptor subunit alpha6 |
| LDEC018354 | esterase e4-like | LDEC019364 | nicotinic acetylcholine receptor subunit alpha7 |
| LDEC023390 | esterase fe4 | LDEC020581 | acetylcholine receptor subunit alpha-like |
| LDEC005269 | esterase fe4-like | LDEC022282 | acetylcholine receptor subunit alpha-like |
| LDEC005858 | Esterase P [Tribolium castaneum] | LDEC000554 | acetylcholine receptor subunit alpha-like |
| LDEC005859 | Esterase P [Tribolium castaneum] | LDEC000556 | acetylcholine receptor subunit alpha-like |
| LDEC005860 | Esterase P [Tribolium castaneum] | LDEC024051 | acetylcholine receptor subunit alpha-like |
| LDEC005861 | Esterase P [Tribolium castaneum] | LDEC004025 | acetylcholine receptor subunit alpha-like 1 |
| LDEC015418 | esterase p-like protein | LDEC017728 | acetylcholine receptor subunit alpha-like isoform x2 |
| LDEC022530 | esterase-6-like protein | LDEC011012 | acetylcholine receptor subunit alpha-type acr-16 |
| LDEC009234 | esterase-6-like protein | LDEC019366 | Acetylcholine receptor subunit beta-like 1 |
| LDEC011783 | esterase-6-like protein | LDEC021578 | acetylcholine receptor subunit beta-like 1 |
| LDEC011787 | esterase-6-like protein | LDEC002850 | acetylcholine receptor subunit beta-like 2 |
| LDEC011788 | esterase-6-like protein | LDEC019226 | transient receptor potential cation channel painless |
| LDEC011823 | esterase-6-like protein | LDEC011527 | transient receptor potential cation channel protein painless |
| LDEC012576 | esterase-6-like protein | LDEC017078 | transient receptor potential cation channel subfamily a member 1 isoform x1 |
| LDEC012577 | esterase-6-like protein | LDEC017081 | transient receptor potential cation channel subfamily a member 1 isoform x1 |
| LDEC012643 | esterase-6-like protein | LDEC018396 | transient receptor potential cation channel subfamily a member 1 isoform x1 |
| LDEC012644 | esterase-6-like protein | LDEC003773 | transient receptor potential cation channel subfamily v member 5 |
| LDEC003520 | esterase-6-like protein | LDEC013671 | transient receptor potential cation channel subfamily v member 5 isoform x2 |
| LDEC003524 | esterase-6-like protein | LDEC013670 | transient receptor potential cation channel subfamily v member 6 |
| LDEC014120 | esterase-6-like protein | LDEC013672 | transient receptor potential cation channel subfamily v member 6 |
| LDEC003999 | esterase-6-like protein | LDEC011197 | transient receptor potential cation channel trpm isoform x13 |
| LDEC004000 | esterase-6-like protein | LDEC011195 | transient receptor potential cation channel trpm-like protein |
| LDEC004001 | esterase-6-like protein | LDEC005427 | transient receptor potential channel pyrexia |
| LDEC004002 | esterase-6-like protein | LDEC017400 | transient receptor potential channel pyrexia isoform x1 |
| LDEC023955 | esterase-6-like protein | LDEC005495 | transient receptor potential channel pyrexia isoform x2 |
| LDEC015419 | esterase-6-like protein | LDEC005426 | transient receptor potential channel pyrexia-like |
| LDEC015702 | esterase-6-like protein | LDEC003213 | transient receptor potential protein |
| LDEC016727 | esterase-6-like protein | LDEC003215 | transient receptor potential protein |

| | | | |
|---|---|---|---|
| LDEC017040 | esterase-6-like protein | LDEC003216 | transient receptor potential protein |
| LDEC017041 | esterase-6-like protein | LDEC023160 | transient receptor potential-gamma protein |
| LDEC018353 | esterase-6-like protein | LDEC011476 | transient receptor potential-gamma protein |
| LDEC021156 | venom carboxylesterase-6 | LDEC011482 | transient receptor potential-gamma protein isoform x1 |
| LDEC019969 | Venom carboxylesterase-6 | LDEC003210 | transient-receptor-potential-like protein |
| LDEC022567 | venom carboxylesterase-6 | LDEC003211 | transient-receptor-potential-like protein |
| LDEC012343 | venom carboxylesterase-6 | LDEC003212 | transient-receptor-potential-like protein |
| LDEC005267 | venom carboxylesterase-6 | LDEC011942 | voltage-sensitive sodium channel |
| LDEC005268 | venom carboxylesterase-6 | LDEC006646 | sodium channel 60E |
| LDEC019608 | venom carboxylesterase-6-like | LDEC006647 | sodium channel 60E |
| LDEC006843 | venom carboxylesterase-6-like | LDEC021105 | sodium channel 60E |
| LDEC021157 | venom carboxylesterase-6-like | LDEC006645 | Sodium channel 60E |
| LDEC024628 | venom carboxylesterase-6-like | LDEC021104 | Sodium channel 60E |
| LDEC001753 | venom carboxylesterase-6-like | LDEC021106 | Sodium channel 60E |
| LDEC022145 | venom carboxylesterase-6-like | LDEC019443 | sodium channel Nach |
| LDEC022654 | venom carboxylesterase-6-like | LDEC006519 | sodium channel Nach |
| LDEC009957 | venom carboxylesterase-6-like | LDEC019444 | sodium channel Nach-like |
| LDEC002822 | venom carboxylesterase-6-like | LDEC020342 | sodium channel Nach-like |
| LDEC012572 | venom carboxylesterase-6-like | LDEC008037 | sodium channel Nach-like |
| LDEC023739 | venom carboxylesterase-6-like | LDEC008039 | sodium channel Nach-like isoform X3 |
| LDEC003521 | venom carboxylesterase-6-like | LDEC022225 | sodium channel protein 60e |
| LDEC000596 | venom carboxylesterase-6-like | LDEC012838 | sodium channel protein 60e |
| LDEC018985 | venom carboxylesterase-6-like [Tribolium castaneum] | LDEC002820 | sodium channel protein 60e-like protein |
| LDEC021266 | venom carboxylesterase-6-like isoform X2 | LDEC024133 | sodium channel protein 60e-like protein |
| LDEC007910 | Major facilitator superfamily domain-containing 12 | LDEC010412 | sodium channel protein nach |
| LDEC020788 | major facilitator superfamily domain-containing 12-like isoform X1 | LDEC017037 | sodium channel protein nach |
| LDEC007655 | major facilitator superfamily domain-containing 9-like | LDEC017502 | sodium channel protein nach |
| LDEC006948 | Major facilitator superfamily domain-containing partial | LDEC008745 | sodium channel protein nach-like |
| LDEC013021 | major facilitator superfamily domain-containing protein 1 | LDEC002517 | sodium channel protein nach-like |
| LDEC000491 | major facilitator superfamily domain-containing protein 10 | LDEC016806 | sodium channel protein nach-like |
| LDEC016787 | major facilitator superfamily domain-containing protein 12-like | LDEC011943 | sodium channel protein para-like protein |
| LDEC010032 | major facilitator superfamily domain-containing protein 6 | LDEC006711 | NA(acetylcholinesterase) |

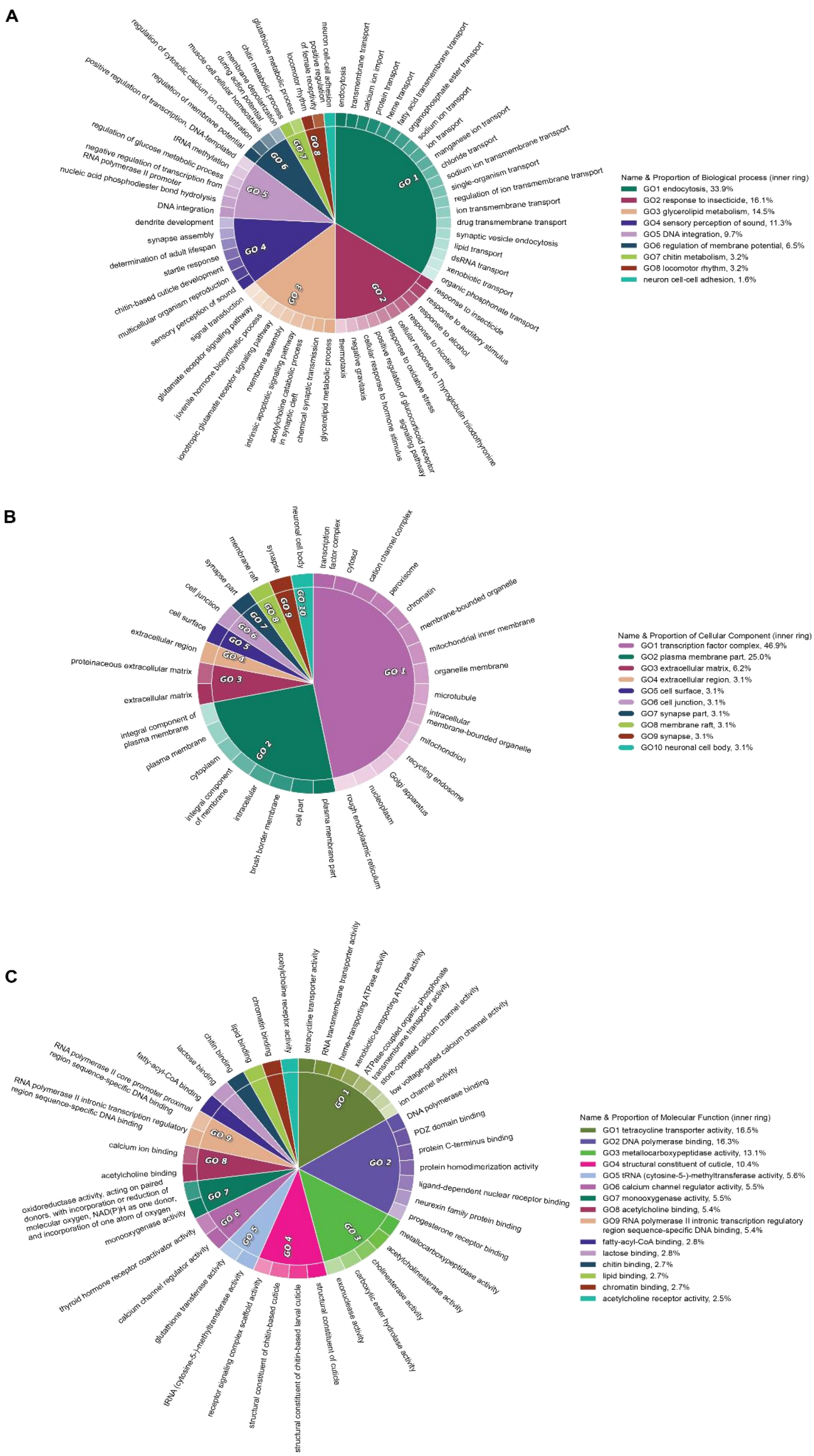

**Figure S41**. *CirGO* plot of overrepresented gene ontology terms for the list of candidate insecticide resistance genes. Terms are labeled as A) biological processes, B) cellular components, and C) molecular functions. Each box along the outer ring represents a gene ontology term, and these are grouped into 'superclusters' of related terms based on semantic similarity (grouping is indicated by color and highlighted by a GO label, with the relative importance indicated by the size of the pie slice).
